# The T cell receptor sequence influences the likelihood of T cell memory formation

**DOI:** 10.1101/2023.07.20.549939

**Authors:** Kaitlyn A. Lagattuta, Aparna Nathan, Laurie Rumker, Michael E. Birnbaum, Soumya Raychaudhuri

## Abstract

T cell differentiation depends on activation through the T cell receptor (TCR), whose amino acid sequence varies cell to cell. Particular TCR amino acid sequences nearly guarantee Mucosal-Associated Invariant T (MAIT) and Natural Killer T (NKT) cell fates. To comprehensively define how TCR amino acids affects all T cell fates, we analyze the paired αβTCR sequence and transcriptome of 819,772 single cells. We find that hydrophobic CDR3 residues promote regulatory T cell transcriptional states in both the CD8 and CD4 lineages. Most strikingly, we find a set of TCR sequence features, concentrated in CDR2α, that promotes positive selection in the thymus as well as transition from naïve to memory in the periphery. Even among T cells that recognize the same antigen, these TCR sequence features help to explain which T cells form immunological memory, which is essential for effective pathogen response.

## Introduction

Early in T cell development, stochastic genome rearrangement on chromosomes 7 and 14 defines each T cell with its own T cell antigen receptor (TCR)^1^. In the thymus and periphery, T cell differentiation depends critically on TCR activation^2–8^. One prominent example is the *PLZF*^high^ innate-like transcriptional fate, which is nearly guaranteed when V(D)J recombination selects *TRAV1-2* and *TRAJ33*, *TRAJ20*, or *TRAJ12*^9^.

Given the central role of the TCR in T cell activation and differentiation, we^10^ and others^11–16^ have identified differences in TCR sequence (e.g. hydrophobicity, V gene selection) using bulk sequencing of flow-sorted T cell populations. However, bulk sequencing obscures the pairing of TCR ɑ and β chains. Furthermore, flow sorting requires predefining T cell states for investigation, and may miss important transcriptional heterogeneity of T cells.

Now, single-cell sequencing assays enable joint profiling of the TCR and transcriptome. In contrast to bulk sequencing methods, single cell technology anchors ɑ and β TCR reads to individual cells, allowing reconstruction of the full TCR ɑ/β heterodimer. Moreover, genome-wide transcriptional analysis can comprehensively define T cell states. Early methods to jointly analyze TCR and transcriptional data^17, 18^ have suggested TCR sequence similarity may correspond to similarity in transcriptional state.

To statistically define the relationship between TCR sequence and T cell state (TCS), we analyze 820,873 T cells with quality-controlled ɑβTCR and transcriptional profiling at single-cell resolution from six published datasets (**Table 1, Supplementary Table 1**). Rather than pre-specifying transcriptional states, we use paired dimensionality reduction to uncover relevant transcriptional states in an unbiased fashion. Our results define four TCR scoring functions that quantify the transcriptional fate predisposition conferred by the TCR. We apply these scoring functions to better understand thymic selection as well as cell state variation within antigen-specific T cell populations.

**Table 1.** Datasets analyzed in this study

| Name | Reference | Cohort attributes | Number of T cells after quality control | Purpose |
| --- | --- | --- | --- | --- |
| Dataset 1 | COMBAT<sup>19</sup> | 77 individuals with COVID-19, 23 with sepsis, 12 with influenza, 10 healthy controls | 282,639 | Twin TCR analysis, developing TCR scoring functions |
| Dataset 2 | Ren et al.<sup>20</sup> | 117 individuals with COVID-19, 17 healthy controls | 211,780 | Developing TCR scoring functions |
| Dataset 3 | Stephenson et al.<sup>21</sup> | 89 individuals with COVID-19, 26 healthy controls | 144,175 | validating TCR scoring functions |
| Dataset 4 | Domínguez Conde et al.<sup>22</sup> | 8 organ donors | 30,702 | validating TCR scoring functions |
| Dataset 5 | Boutet et al.<sup>23</sup> | 4 healthy individuals | 136,546 | Assessing TCR scoring functions within antigen-specific populations |
| Dataset 6 | Suo et al.<sup>24</sup> | 6 prenatal samples | 13,930 | Assessing TCR scoring functions in the thymus |

## Results

### T cell transcriptional state annotation

To construct an accurately annotated reference of T cells, we used Dataset 1 (from the COMBAT consortium^19^, **Table 1**) with 371,621 T cells from 122 individuals with multimodal Total-seq profiling of mRNA and surface proteins. We clustered the cells in two ways: first agnostic to protein expression (clusters A1-A9, **Figure 1a, Supplementary Figure 1**), and second, incorporating traditional surface markers via a linear multimodal strategy^25^ (clusters B1-B9, **Supplementary Figure 2**). Cluster A9, representing the *PLZF*^high^ innate-like TCS, accounted for nearly all canonical MAIT or NKT TCRs (**Figure 1b-c**). Clusters B1-B9 delineated CD4, CD8, central memory (CM), and effector memory (EM) states based on a curated list of 10 surface proteins including CD45RO and CD45RA (**Supplementary Table 2, Supplementary Figure 2, Methods**). To standardize cell state definitions across datasets, we projected all additional datasets (**Table 1**) into these two embeddings and transferred annotations via k-nearest-neighbors classification (k=5, **Methods**).

**Figure 1.**
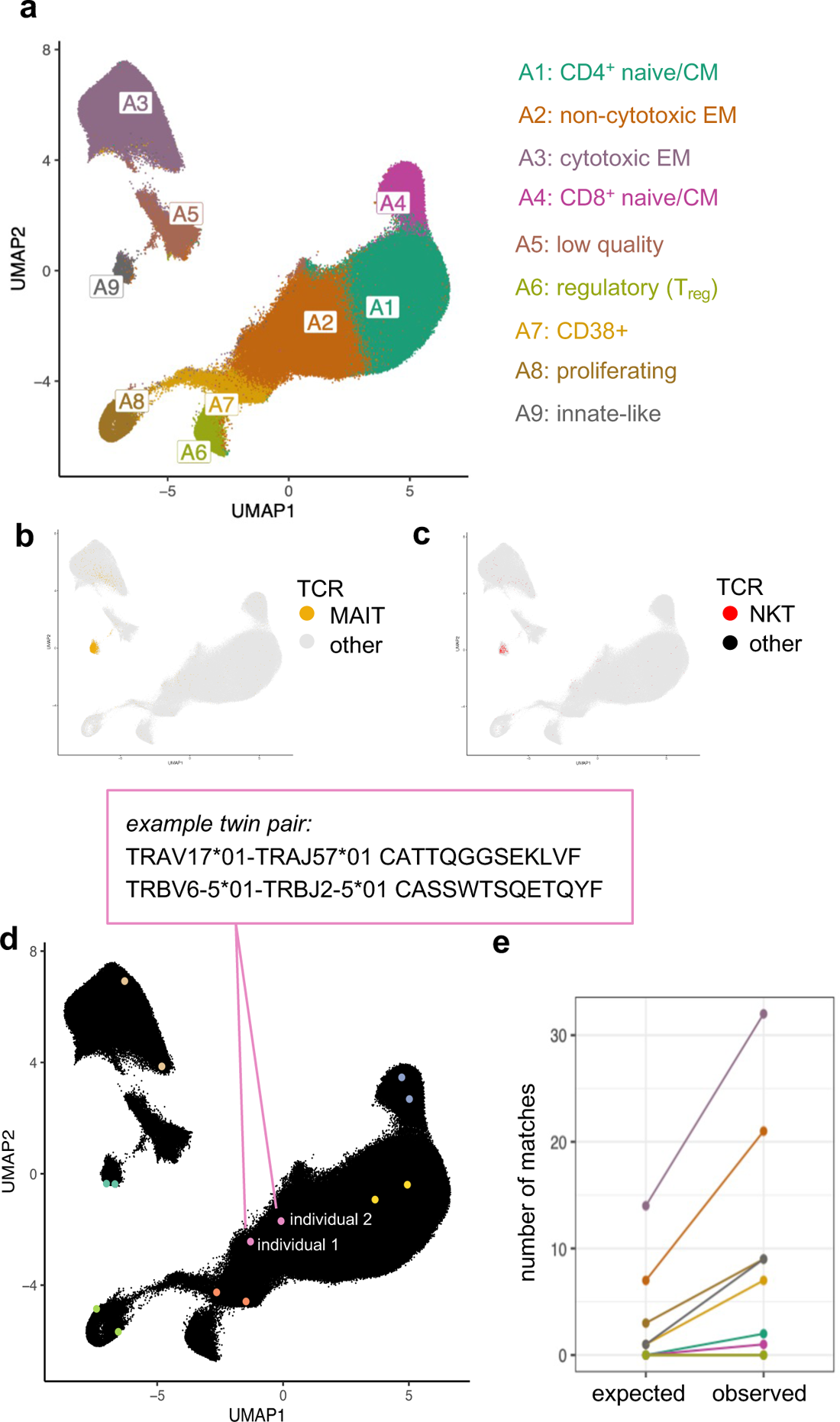
**(a)** UMAP of T cells from Dataset 1 based on the first 20 batch-corrected principal components (PCs) of gene expression. Cells are colored by transcriptional cluster, named according to expression of marker genes and proteins (Supplementary Figure 1). **(b)** T cells from Dataset 1, colored yellow if their paired TCR sequence includes the canonical genes for MAIT cells (*TRAV*1-2* with *TRAJ*20, *33,* or *\*12*, and *TRBV*6* or *\*20*) or **(c)** colored red if their paired TCR sequence includes the canonical genes for NKT cells (*TRAV10* with *TRAJ18* and *TRBV25-1*) or gray otherwise. **(d)** Seven example TCR-twin cell pairs from Dataset 1 are highlighted in distinct colors. Within a twin pair, TCRs with matching sequences were observed in two different individuals. **(e)** Number of transcriptional cluster matches for TCR twin cell pairs in each cluster. Real observed counts are compared to expected counts based on random sampling of T cells without regard for TCR sequence.

### Transcriptional fate matching between T cells with the same TCR sequence from different individuals

We first used Dataset 1 (**Table 1**) to assess whether perfect TCR sequence matching raised the likelihood of T cell state matching. Within an individual, identical TCR sequences likely reflect an expanded T cell clone, comprised of cells that tend to be transcriptionally similar (**Supplementary Figure 3**). To avoid clonally related cells, we focused on identical TCR sequences sampled from two different individuals. We identified 115 pairs of “TCR twins:” two T cells from two different individuals with the same TCR ɑ and β amino acid sequences (**Figure 1d**, **Supplementary Table 3**, **Methods**). We assessed whether their transcriptional states (A1-A9) were concordant more often than expected by chance. Under the null, we would expect 25 (of 115) transcriptionally concordant TCR twin pairs (see **Methods**). Instead, we observed 80 (*P* = 6.1 x 10^-28^, binomial test, **Figure 1e**, **Methods**). To assess if this enrichment was explained by MHC(-like) restriction, we repeated our analysis with MAIT and NKT TCRs removed, and observed an enrichment in concordant states (*P* = 2.3 x 10^-21^). Partitioning into CD4^+^ and CD8^+^ populations did not obviate enrichment either (*P* = 4.0 x 10^-9^, *P* = 0.00018). Finally, recognizing that SARS-CoV-2 infection is a potential confound, we filtered to individuals that were PCR-negative for SARS-CoV-2 but continued to observe enrichment (13 matches versus 6 expected by chance, *P* = 0.00048). These results suggest a consistent influence of TCR sequence on T cell fate in unrelated individuals.

This analysis is limited to TCRs found in more than one individual (“public TCRs”), which comprise <0.01% of the TCRs in this cohort. Public TCRs have distinct structural features^26^, and could demonstrate transcriptional state matching due to recognition of common antigens. We hypothesized, however, that these results were driven to some extent by distinct biophysical features of the TCR sequences. If so, similar, but not identical, TCR sequences would also promote similar T cell fates.

### A multidimensional approach to uncover TCR sequence features that guide T cell fate

To extend our study to private TCRs, we converted each TCR sequence into a vector of biophysical features. Consistent with other numerical representations of the TCR^17, 27^, we translated each amino acid residue in both the α and β chains of the TCR into five Atchley factors^28^. These factors correspond to hydrophobicity, size, charge, secondary structure, and heat capacity. Applying this to α and β complementarity-determining and framework regions of the TCR yielded 1190 biophysical features. Excluding invariant positions, framework regions, and adding interaction terms between adjacent residues yielded 1225 TCR features (**Supplementary Table 4**, **Methods**).

We aimed to identify TCR sequence effects on TCS in an unbiased fashion, without restricting to preselected cell states. To find generalizable associations, we combined Dataset 1 with a second published dataset (“Dataset 2”, **Table 1, Supplementary Figure 4, Methods**), resulting in 573,082 quality-controlled T cells from 256 individuals. We applied regularized Canonical Correlation Analysis (rCCA) (**Figure 2a-c, Methods**). Each axis identified by rCCA denotes a weighted sum of gene expression PC scores (a transcriptional state) that correlates with a weighted sum of TCR sequence features.

**Figure 2.**
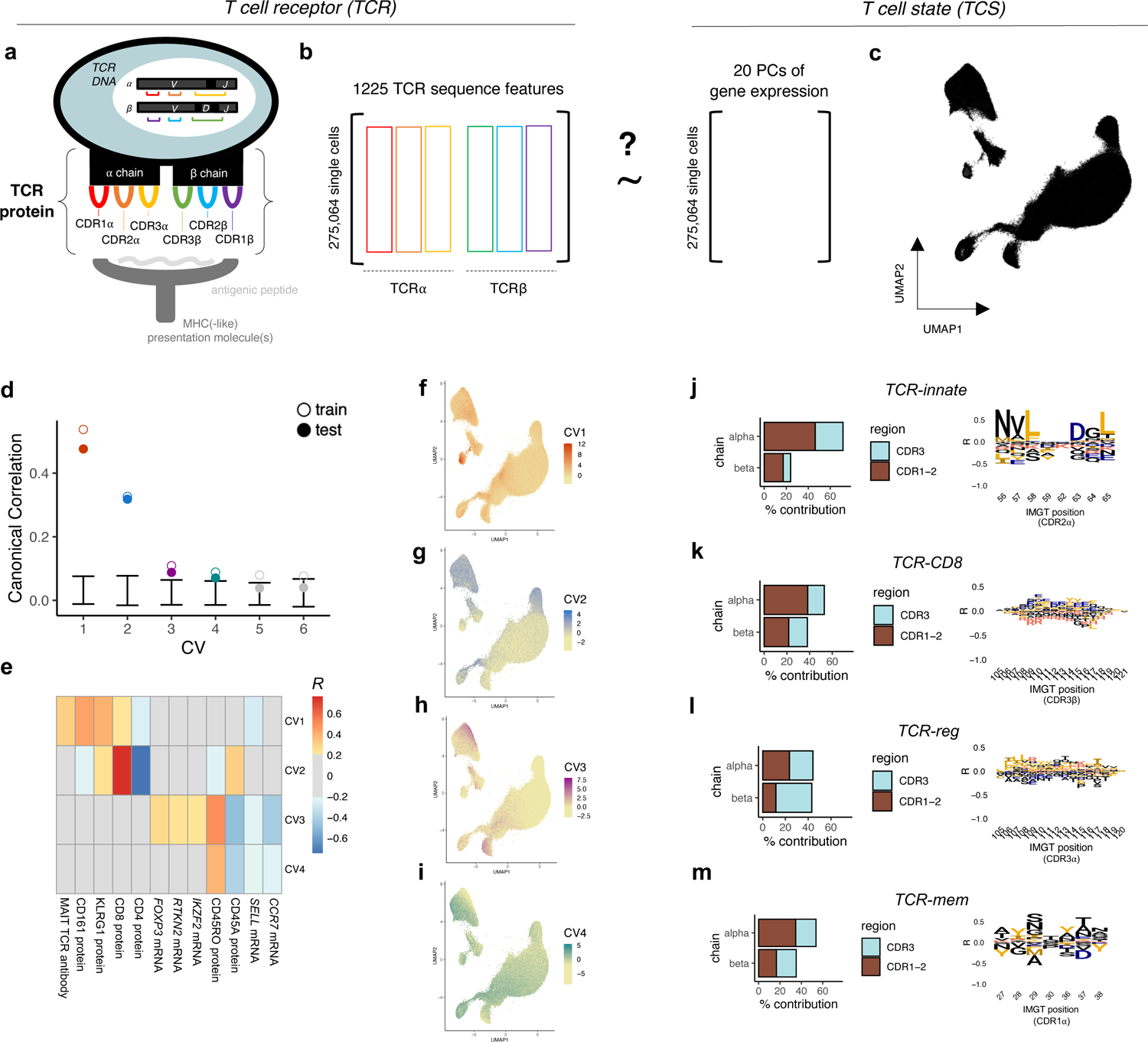
**(a)** Schematic of a T cell recognizing antigen through the T cell antigen receptor (TCR), an αβ heterodimeric surface protein encoded by V, D, and J genes that are stochastically rearranged in each T cell. Black regions in between V, D, and J genes represent non-templated nucleotide insertions. The TCR surface protein contains six complementarity-determining regions (CDRs) that extrude toward the presented antigen: CDR1α, CDR2α, CDR3α, CDR1β, CDR2β, and CDR3β. **(b)** Our paired dimensionality reduction approach to uncover ways in TCR sequence variation impacts T cell transcriptional fate. We applied regularized Canonical Correlation Analysis (rCCA) to paired measurements in each cell: 1) TCR sequence, represented by 1225 quantitative features from the six CDRs 2) T cell state, represented by 20 batch-corrected principal components (PCs) of gene expression. **(c)** A visual projection (UMAP) of T cell states captured by 20 batch-corrected PCs of gene expression in Dataset 1. **(d)** Canonical correlations between TCR and T cell state detected in training and held-out testing observations. Error bars denote the minimum and maximum canonical correlations observed in 1000 permutations of the data. **(e)** Heatmap depicting Pearson correlation between genes and proteins in Dataset 1 and the continuous T cell state identified by each canonical variate (CV). **(f-i)** UMAP of Dataset 1 T cells, colored by T cell state scores for CV1, CV2, CV3, and CV4, respectively. **(j-m)** Visualization of TCR-innate, TCR-CD8, TCR-reg, and TCR-mem scoring function. Left: Percent contribution from each TCR region toward the TCR scoring function; right: amino acid contributions to the TCR scoring function, visualized as marginal correlations to each amino acid in a select TCR region. Other TCR regions displayed in SFig12.

We mitigated technical confounding in our rCCA implementation. First, to prevent confounding by variable clone size, we selected one cell at random to represent each clone. We confirmed that results were invariant to the specific set of cells sampled (**Supplementary Figure 5a**, **Methods**). To mitigate overfitting, we added a ridge regularization to the covariance matrix for each of the inputs to rCCA, using 5-fold cross-validation to tune both lambda penalty values. To assess overfitting, we randomly assigned 68 donors to a validation test set, such that ∼30% of clones (29.3%) were held out from training.

### Regularized canonical correlation analysis identifies four T cell fates informed by TCR sequence

We observed canonical correlations between TCR sequence and TCS descending from *R* = 0.55. To assess the statistical significance of each canonical correlation, we permuted our data 1000 times and re-applied rCCA (**Methods**). We observed empirical *P* values < 0.001 for both training and held-out testing data for the first four canonical variates (CVs) (**Figure 2d**).

We observed that framework regions and interactions between non-adjacent residues did not increase correlations between TCR and TCS (**Supplementary Figure 5b**). Training separately in each dataset confirmed that the TCR - TCS associations learned in Dataset 1 replicated in Dataset 2 and vice versa (**Supplementary Figure 6, Methods**).

To interpret the four continuous T cell states identified by rCCA, we examined CVs1-4 in terms of cell scores and expression correlates (**Figure 2e, Supplementary Table 5, Methods**). Cells scoring highest on CV1 localized to transcriptional cluster A9 (**Figure 2f**), the innate-like, *PLZF*^high^ transcriptional fate for canonical MAIT and NKT TCRs (**Figure 1b-c, Supplementary Note 1, Supplementary Figure 7**). CV2 tracked closely with CD8 versus CD4 surface expression (**Figure 2e**), delineating CD4+ T versus CD8+ T populations (**Figure 2g, Supplementary Figure 8**). These results point to families of peptide presentation molecules as the primary source of covariation between TCR sequence and T cell state. Indeed, it is well-established that unconventional (MR1, CD1d), MHC class I, and MHC class II families each prefer biophysically distinct ɑβTCR sequences^9,13, 14, 29–33.^

In addition to these known relationships, rCCA proposed novel connections between TCR sequence and TCS. CV3 highlighted TCR sequence similarity between *FOXP3*-expressing CD4+ regulatory T (T_reg_) cells and KIR+*HELIOS*+ CD8 T cells (**Figure 2h, Supplementary Figure 9**), which have recently been described as human CD8+ T_regs_^34^. This suggests that the same TCR sequence features may promote suppressive functional states in both the CD4 and CD8 compartments. Most strikingly, CV4 appeared to capture TCR sequence differences between naive and memory T cells (**Figure 2i, Supplementary Figure 10a**). Surface protein measurements indicated that both effector memory (EM) and central memory (CM) CD4+ and CD8+ T cells scored highly on CV4 (**Supplementary Figure 10b-c**). This raises the intriguing possibility that some sequence features render the TCR more generally prone to activation.

### TCR scoring functions quantify TCR sequence features that inform T cell fate

The continuous T cell states defined by rCCA nominated four contrasts in T cell state for further study: *PLZF*^high^ vs. other, CD8T vs. CD4T, T_reg_ vs. T_conv_, and memory vs. naïve. For each of these recognizable T cell fate decisions, we used logistic regression on the same observations from Datasets 1 and 2 to learn a more precise predictive weighting scheme on the 1225 TCR sequence features (**Supplementary Figure 11, Supplementary Tables 6-7, Methods**). We named each predictive weighting scheme, or TCR scoring function, by the T cell state of interest: “TCR-innate’” to predict innate-like *PLZF*^high^ state, “TCR-CD8” to predict CD8+ state, “TCR-reg” to predict T_reg_ state, and “TCR-mem” to predict memory state.

To interpret each of these TCR scoring functions, we examined relative contributions from each CDR and amino acid residue (**Methods**). TCR-innate was driven predominantly by *TRAV* gene selection (CDR1α-CDR2α, **Figure 2j, Supplementary Figure 12a**). As expected from previous studies^13, 14^, TCR-CD8 high sequences were depleted for positive charge in the junctional mid-region of CDR3 (**Figure 2k, Supplementary Figure 12b**). TCR-reg reflected increased hydrophobic CDR3β residues in CD4 T_regs_ (**Supplementary Figure 12c**), consistent with previous reports^10, 12^. Paired αβ TCR sequencing data revealed that enrichment for hydrophobic amino acids extended to CDR3α (**Figure 2l**). For TCR-mem, variance components highlighted the importance of the *TRAV* gene encoding CDR1α and CDR2α (**Figure 2m, Supplementary Figure 12d**). CDR2α position 63 made the strongest contribution to TCR-mem, such that TCRs with a negatively charged residue at CDR2α p63 exhibited 20% greater odds of memory formation compared to TCRs with a positively charged residue at CDR2α p63 (**Supplementary Figure 13**). *TRAV* gene associations to memory state were generally concordant between the CD4 and CD8 lineages (*R* = 0.67, *P* = 1.8 x 10^-6^), with 10 significantly associated genes (*P* < 5.7 x 10^-4^ = 0.05/(44 genes x 2 lineages = 88 tests)) in both lineages (**Supplementary Table 8, Methods**). These results are in contrast to previous reports describing minimal difference between naïve and memory T cell V gene usage^35, 36^. But, these previous studies used bulk TCRβ sequencing, which does not capture *TRAV*. We assessed TCR-reg and TCR-mem separately in CD4+ and CD8+ T cells, and observed that these TCR scoring functions were equally applicable to both lineages (**Supplementary Figure 14, Methods**).

### TCR scoring functions generalize across individuals

We considered the possibility that the associations between TCR sequence and T cell state were driven by a subset of individuals. We first stratified each dataset by clinical status (COVID, Sepsis, Influenza, none of the above), and used mixed-effects logistic regression to calculate *β_TCRscore,_* the association between each TCR score and its target T cell state (log odds ratio for T cell state per standard deviation increase in TCR score, see **Methods**). In each clinical stratum, we observed a statistically significant positive association for each TCR scoring function (24 tests, maximum *P =* 1.1 x 10^-3^, **Figure 3a-d, Supplementary Table 9**). Reassured that our TCR scoring functions were not driven by clinical subset, we considered the possibility of an unknown individual-level mediator, such as *HLA* genotype. We computed *β_TCR-innate_, β_TCR-CD8,_ β_TCR-reg_*, and *β_TCR-mem_* within each individual’s T cells separately (**Supplementary Table 10**), and estimated the proportion of individuals for whom our TCR score does not raise the odds of its target T cell state (the local False Sign Rate^37^). Random effects meta-analysis indicated a near-zero proportion for each of our four TCR scoring functions (<1e-6, **Supplementary Figure 15, Supplementary Table 11, Methods**). We concluded that since our TCR scoring functions were robust to inter-individual variation, they should generalize to unseen samples.

**Figure 3.**
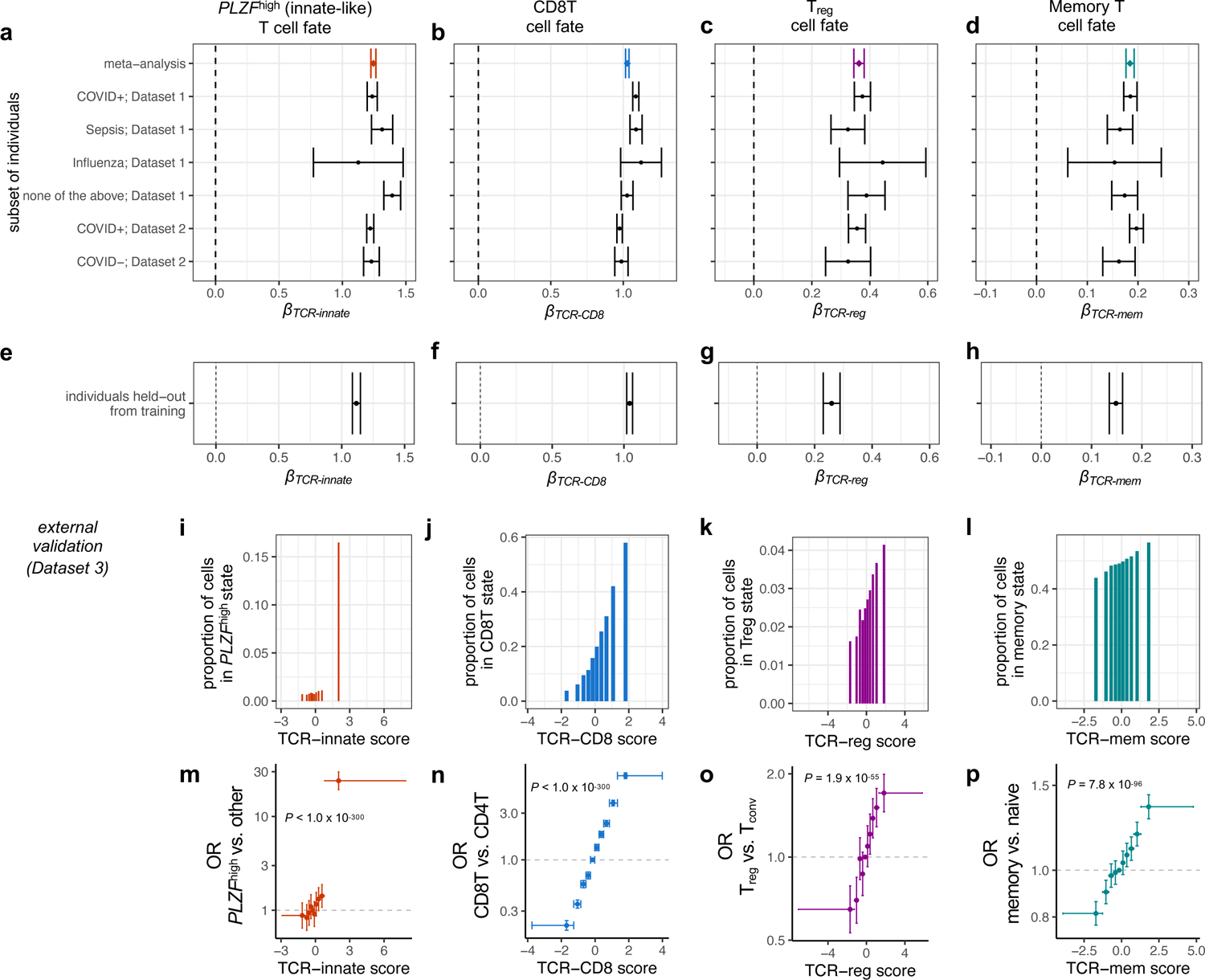
**(a)** Forest plot depicting the association between TCR-innate and *PLZF*^high^ T cell state for T cells from each subset of individuals. *βTCR-innate* is computed via mixed-effects logistic regression, with TCR-innate scaled to mean 0 variance 1. Error bars denote 95% confidence intervals. Meta-analytic *βTCR-innate* is estimated by fixed-effects inverse-variance-weighted meta-analysis. Remaining forest plots depict the same association tests for **(b)** TCR-CD8 and CD8T cell state, **(c)** TCR-reg and Treg cell state, and **(d)** TCR-mem and memory T cell state. **(e-h)** 95% confidence intervals for *βTCR-innate, βTCR-innate, βTCR-innate,* and *βTCR-innate*, respectively, in the individuals from Dataset 1 and Dataset 2 held out from TCR score training. **(i-l)** Proportion of T cell clones from an external validation dataset (Dataset 3) observed in each T cell state of interest within each decile of its corresponding TCR score. **(m)** Each point represents a decile of the TCR-innate score in Dataset 3, with a horizontal bar spanning from its minimum to maximum value. We compute the odds ratio (OR, y-axis) for *PLZF*^high^ T cell state for T cells in each decile compared to T cells in the fifth decile. 95% CIs (error bars) and *P* values computed via mixed-effects logistic regression. **(n-p)** Dataset 3 T cells as in (i), for CD8T, Treg, and memory T cell states, and their corresponding TCR scores.

We next applied our TCR scoring functions to data outside of our training set. In the 30% of Dataset 1 and Dataset 2 T cell clones held out from training, we observed replication of each TCR scoring function (**Figure 3e-h, Methods**). In an external dataset of peripheral blood T cells (“Dataset 3”, **Table 1, Supplementary Figure 16**), we observed a consistent and statistically significant increase in odds of the T cell state of interest for each TCR scoring function (**Figure 3i-p, Supplementary Table 12**). Compared to T cells in the lowest TCR-mem decile, T cells in the highest TCR-mem decile had a 66% greater odds of being observed in a memory state (OR = 1.66, 95% CI = [1.57 - 1.76], *P* = 7.9 x 10^-66^, **Supplementary Table 13**). These results indicate substantial role for TCR-mem shaping the odds of T cell memory formation.

### Alternative TCR scoring schemes

We next benchmarked our TCR scoring functions against existing TCR metrics. We previously developed a Treg TCR scoring function, “TiRP,” using the TCR β chain alone^10^. With the additional information of the α chain, TCR-reg clearly outperformed TiRP (*β_TiRP_* = 0.14, 95% CI = [0.10 – 0.18]; *β_TCR-reg_* = 0.29, 95% CI: [0.25 – 0.33]; Dataset 3 CD4T cells). Amino acid interaction strength^38^ (AAIS) has been postulated to estimate a TCR’s average affinity to pMHC^39^, but this has not been directly tested. In Dataset 3, increasing AAIS corresponded to an increase in the odds of memory state only when applied to CDR3 amino acids (*β_AAIS_* = 0.02, 95% CI = [0.003 – 0.03], *P* = 0.008). The effect size for AAIS was minimal compared to TCR-mem (*β_TCR-mem_* = 0.13, 95% CI: [0.12 – 0.15]), however. Including AAIS as a covariate did not substantially change the estimated effect size of TCR-mem (conditional *β_TCR-mem_* = 0.13, 95% CI: [0.12-0.14], heterogeneity *P* = 0.44). TCR-reg and TCR-mem clearly outperform these alternative TCR scoring functions, by capturing both α and β TCR sequence features that promote recognition in the context of the TCR-pMHC interface.

We next wanted to assess if more complex models would provide better TCR scoring functions. For each of the four T cell states of interest, we trained a binary classifier using a Convolutional Neural Network (CNN), which enables the detection of TCR amino acid motifs and possibly nonlinear effects (**Methods**). However, this deep learning approach provided no substantial benefit in discovery or external validation data (**Supplementary Figure 17, Supplementary Tables 14-15**).

### Untranslated products of V(D)J recombination do not affect T cell fate

Because stochastic V(D)J recombination precedes T cell fate decisions, TCR sequence associations to T cell state likely reflect causal effects of V(D)J recombination. However, a causal pathway that begins with V(D)J recombination and ends with T cell state likely includes several important biological mediators. To better understand these mediators, we decomposed V(D)J recombination into three products: (1) DNA-level excisions and insertions, (2) amino acid changes in the surface TCR, and (3) antigen recognition. To isolate (1) from (2), we analyzed nonproductive V(D)J recombination sequences that are not translated into surface TCR proteins. Then, to distinguish (3) from (2), we examined the TCR sequences and T cell states of antigen-labeled single cells.

Nonproductive TCR sequence transcripts can be detected when an out-of-frame V(D)J recombination event on one chromosome is followed by an in-frame V(D)J recombination event on the other^40^. Due to stop codons and frameshift errors, these nonproductive TCRs represent V(D)J genome rearrangements that are not translated into surface antigen receptors^41^. To assess whether these DNA-level changes are sufficient to produce the observed effects on T cell state, we applied our TCR scoring functions to nonproductive TCR sequences (“Dataset 4”, **Table 1, Supplementary Figure 18**). We observed no evidence of association for any of the four TCR scoring functions (*P* > 0.05). Down-sampling did not obviate associations for productive TCRs, confirming that the lack of association was not due to reduced statistical power (**Supplementary Figure 19, Supplementary Table 16, Methods**). We concluded that the DNA-level excisions and insertions from V(D)J recombination are in general not sufficient to affect T cell state; recombination products must be expressed at the protein level.

### TCR sequences that recognize the same antigen can be poised for different fates

Having localized the association signal to the TCR protein, we next wondered whether the key mediator might be each TCR’s cognate pMHC. If so, TCR sequence would direct T cell fate only insofar as it encourages recognition of pMHCs that elicit particular types of T cell differentiation. Our TCR scores would be no more than ‘TCR fingerprint’ mirrors of the mediating pMHCs, and would fail to associate with transcriptional state among T cells with the same antigen specificity.

To assess evidence for this pMHC-centric model, we downloaded public antigen profiling data^23^ (“Dataset 5”, **Table 1**). Briefly, 80 million CD8+ T cells from the peripheral blood of four human donors were exposed to 44 pMHC barcoded Dextramers. Multimodal sequencing then assayed the ɑβTCR sequence, transcriptome-wide expression, and pMHC Dextramer counts for each Dextramer-positive cell. Using Symphony and k-nearest neighbors, we assigned T cell states based on our multimodal T cell reference (**Figure 4a**).

**Figure 4.**
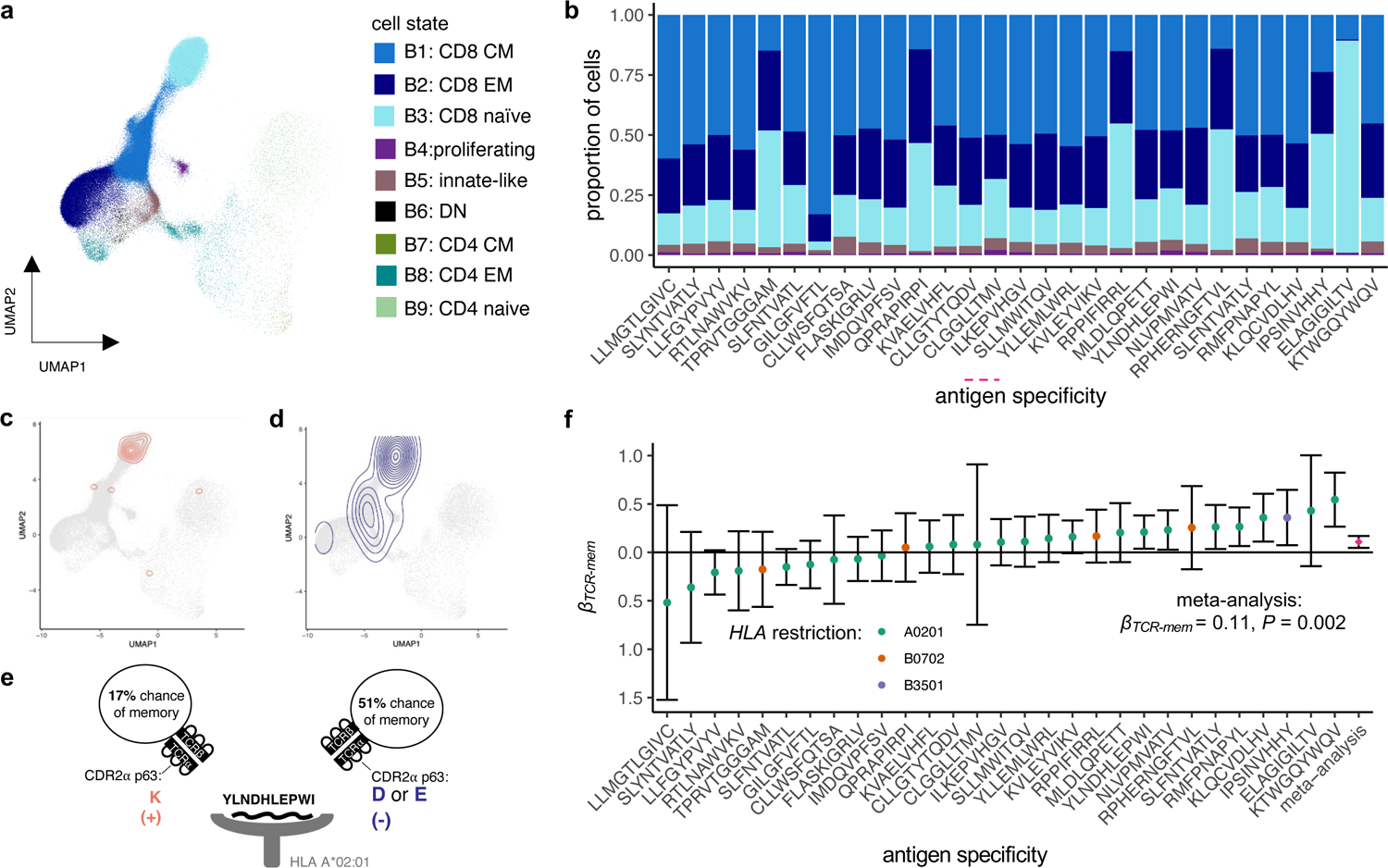
**(a)** T cells from Dextramer-labeled dataset (Dataset 5) projected onto UMAP coordinates of a low-dimensional transcriptional space defined by Dataset 1, with strong influence from Total-seq antibody counts for CD4, CD8, CD45RO, and CD45RA (Supplementary Figure 2). **(b)** Cell state proportions among each of the antigen-specific populations. **(c)** T cells from Dataset 5 as in (a); the distribution of YLDNHLEPWI-specific T cells with a positively charged residue at CDR2α position 63 is highlighted in red. **(d)** The distribution of YLDNHLEPWI-specific T cells with a negatively charged residue at CDR2α position 63 is highlighted in blue. **(e)** Schematic depicting the two types of YLDN-specific T cells contrasted in (c) and (d), respectively. **(f)** TCR-mem effect size (*βTCR-mem*) within each antigen-specific population, quantified as the natural logarithm of the memory vs. naive log(OR) per unit increase in TCR-mem. Meta-analytic *βTCR-mem*, 95% CI, and *P* value are computed via random-effects meta-analysis.

Following custom normalization of Dextramer UMI counts (**Supplementary Note 2, Supplementary Figures 20-22, Supplementary Table 17**), we observed a mixture of transcriptional states within each antigen-specific population (**Figure 4b, Supplementary Figure 23**). This transcriptional heterogeneity is consistent with other tetramer-sorted scRNAseq studies^42–44^, and offers evidence against the notion that antigen specificity fully determines T cell transcriptional phenotype. We wondered if TCR-mem might help to explain which T cells within each antigen-specific population exhibited a memory transcriptional state.

Within each antigen-specific population of cells, we tested the association between TCR-mem and memory state using a single cell per TCR clone (**Methods**). We observed *β* >0 for the majority of antigens (19/29), including seven antigen-specific populations with a nominally significant (one-tailed *P* < 0.05) result (**Supplementary Table 18**). For example, only some T cells capable of recognizing self-antigen BCLX peptide YLDNHLEPWI had successfully reached a memory transcriptional state. T cells which had successfully transitioned from naïve to memory bore TCRs with significantly higher TCR-mem scores (logistic regression *β_TCR-mem_* = 0.21, *P* = 0.02). CDR2α position 63, the strongest contributor to TCR-mem (**Supplementary Figure 13d-e),** delineated two functionally distinct types of YLND-specific T cells: 17% of those with negatively charged residue at p63 had reached a memory state, compared to 51% for those with a positively charged residue at p63 (**Figure 4c-e**).

We conducted a meta-analysis across antigens, given the lack of power within most antigen-specific populations (median TCRs per antigen: 196, range: 1 - 641). We observed a significant effect of TCR-mem on memory state, adjusted for antigen specificity (**Figure 4f**, logistic regression *β_TCR-mem_* = 0.11, *P* = 0.002, **Methods**). We observe minimal evidence for a difference in *β_TCR-mem_* before and after adjusting for antigen specificity (*P* = 0.27, **Supplementary Table 19, Methods**), and minimal heterogeneity in *β_TCR-mem_* across antigen-specific populations (*I^2^ =* 29.8%, *H^2^* = 1.42*, Q =* 39.9*, P* = 0.07). TCR-mem associations hold after adjusting for Dextramer staining intensity (**Methods**) as a proxy for TCR-pMHC affinity, which is thought to contribute to memory T cell development^45^. These results suggest that TCR-mem sequence features predispose pMHC recognition in general, regardless of the cognate antigen.

### Thymic selection pressures on the TCR sequence continue in the periphery

TCR activation results from recognition of a compound structure: an antigenic peptide presented on an MHC(-like) molecule^46, 47^. Given the consistent association of TCR-mem to memory state across antigenic peptides (**Figure 4d**), we hypothesized that TCR-mem reflects affinity to the underlying MHC(-like) molecule. Specifically, to promote recognition of many different pMHCs, such affinity would be focused on elements of MHC that are conserved across MHC genes and alleles. This TCR property, referred to as “generic MHC reactivity”, has been theorized based on thymic development in TCR-transgenic mice^48^.

If TCR-mem reflects generic MHC reactivity, then TCR-mem sequence features should also help to explain which T cells survive thymic positive selection. Only T cells with a sufficient signaling response to pMHC survive thymic positive selection, which is marked by progression from a double-positive (DP) phenotype to a single-positive (SP) phenotype. Thus, if TCR-mem reflects generic MHC reactivity, SP T cells should have higher TCR-mem compared to DP T cells, 90%^2,3^ of which never progress to the SP stage.

Thus, we compared TCR-mem scores between SP and DP prenatal T cells (7,264 clones, “Dataset 6”, **Table 1, Methods**). TCR-mem was designed to describe differences between naïve and memory TCRs in the periphery. We observed, however, that the same TCR weighting scheme also described differences between DP and SP TCRs (**Figure 5a**, *β_TCR-mem_* = 0.14, *P* = 1.2 x 10^-7^, **Supplementary Table 20)**. Strikingly, the TCR-mem difference between SP and DP thymic T cells was statistically indistinguishable from the TCR-mem difference between memory and naïve peripheral T cells (heterogeneity *P* = 0.49). Thus, TCR differences between peripheral naive and memory T cells appear to echo TCR filtering by thymic positive selection (**Figure 5b**). The influence of TCR-mem persists in the absence of foreign and peripheral antigens, suggesting that TCR-mem reflects generic pMHC reactivity^48^. While thymic selection imposes a minimum threshold for pMHC reactivity, TCRs that survive this threshold by a wider margin appear more likely to reach a memory T cell state in the periphery.

**Figure 5.**
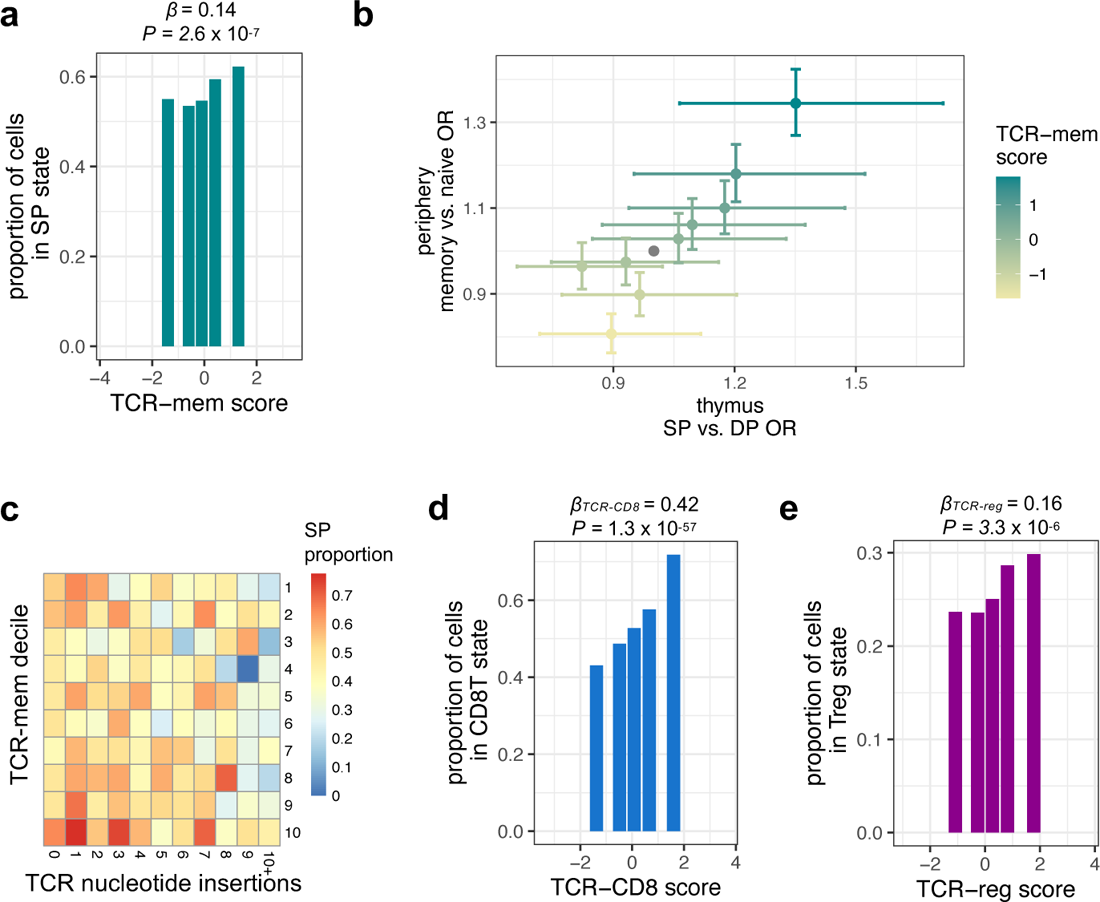
**(a)** Proportion of SP thymocytes in Dataset 6 within each decile of the TCR-mem score. *β* and *P* are computed via mixed effects logistic regression, predicting SP status based on TCR-mem score (Methods). **(b)** Each point represents a TCR-mem decile, plotted according to its association with positive selection in the thymus (Dataset 6, x-axis) and to its association with transitioning from naïve to memory in the periphery (Dataset 3, y-axis). ORs and 95% CIs are estimated via mixed-effects logistic regression. **(c)** Heatmap of SP proportion for each combination of TCR-mem score decile and number of TCR nucleotide insertions in Dataset 6. **(d)** Proportion of SP thymocytes in Dataset 6 observed in a CD8 T cell state within each decile of the TCR-CD8 score. *βTCR-CD8* and *P* are computed via mixed effects logistic regression, predicting CD8 T cell state based on TCR-CD8 score. **(e)** Proportion of SP thymocytes in Dataset 6 observed in a Treg cell state within each decile of the TCR-reg score. *βTCR-reg* and *P* are computed via mixed effects logistic regression, predicting Treg cell state based on TCR-reg score.

An alternative possibility we considered is that higher TCR-mem T cells had developed earlier in life, and therefore accrued more opportunities to transition their transcriptional state. This time-related confound would require systematic shifts in V(D)J recombination with human age. However, we observed no relationship between age and TCR-mem (**Supplementary Figure 24**) or *β_TCR-mem_* (**Supplementary Figure 15d**).

Prenatal TCR sequences from Dataset 6 allowed us to further extend our age-related line of inquiry. Some prenatal T cells lack *DNTT* expression, precluding non-templated insertion of TCR nucleotides and resulting in systemically shorter TCR sequences^49^. To identify these age-related TCRs, we applied IGoR^50^ to infer the number of nucleotide insertions in each thymic TCR (**Supplementary Figure 25**). We observed that additional nucleotide insertions corresponded to a decrease in the odds of thymic positive selection, but this effect did not account for effect of TCR-mem. Controlling for the number of nucleotide insertions, TCR-mem actually demonstrated a stronger effect on the odds of positive selection (**Figure 5c**, conditional *β_TCR-mem_* = 0.20, 95% CI [0.12-0.29], *P* = 2.3 x 10^-6^, compared to unconditional *β_TCR-mem_* = 0.14, 95% CI [0.09 – 0.19], *P* = 2.6 x 10^-7^, **Supplementary Table 20, Methods**). Thus, TCR-mem associations cannot be explained by developmental timepoint.

In contrast to TCR-mem, TCR-innate was not applicable to thymic data. Thymic T cells expressing canonical MAIT and NKT TCRs had not yet reached the innate-like, *PLZF*^high^ transcriptional fate (**Supplementary Figure 26**). This is consistent with previous reports showing that *PLZF*^high^ fate acquisition is dependent on peripheral antigen recognition^51–53^. TCR-reg and TCR-CD8, however, were applicable to thymic data (**Figure 5d-e**, *β_TCR-CD8_* = 0.42, *P* = 1.3 x 10^-57^; *β_TCR-reg_* = 0.16, *P* = 3.3 x 10^-6^, **Methods, Supplementary Table 20**), indicating that these TCR-TCS associations are not dependent on peripheral antigen recognition. Evidently, TCR sequence features shape T cell differentiation outcomes in both the thymus and periphery, influencing which T cells are able to generate an effective immune response.

## Discussion

In this study, we define four TCR scoring functions that estimate the TCR’s contribution to four T cell fates using a novel and unbased statistical approach. These scoring functions are robust across numerous genetic and clinical contexts; they explain differential transcriptional states even among T cells that recognize the same antigen.

Our unsupervised, quantitative approach allows us to understand the relative strength of previously observed TCR-TCS connections. The most deterministic relationship belongs to MAIT cells. Structural studies have shown that MHC-like molecule MR1 buries small metabolite antigens so that they do not contact TCR^54, 55^. Consequently, MAIT TCRs need only recognize MR1, which marks a highly distinct transcriptional population of *PLZF*^high^ innate-like T cells. This tight link from ɑβ TCR sequence to T cell fate is unusual; if there were other relationships of similar magnitude, we contend that our model would have identified them.

We observe elevated CDR3 hydrophobicity in KIR+*HELIOS*+ CD8 T cells, consistent with previous reports^18^. Schattgen et al. hypothesized that this CD8 population may be “MHC-independent, noncanonical, or self-specific.” Our framework unifies this observation with CD4 investigations^10, 15, 16^: KIR+CD8+ T cells, which may functionally represent human CD8 T_regs_^34^, seem to arise from the same TCR selection processes as CD4 T_regs_. In general, hydrophobic and aromatic (F, L, I, C, Y, W) junctional CDR3 residues (both α and β) may increase a T cell’s likelihood of recognizing self-antigens, driving *FOXP3* Treg fate in the case of CD4 co-receptors and KIR+*HELIOS*+ fate in the case of CD8 coreceptors.

We take particular interest in TCR-mem, because it describes TCR sequence features that are generally advantageous for reaching a memory T cell state. Antigen specificity does not appear to confer this advantage; even within antigen-specific populations, higher TCR-mem corresponds to a greater likelihood of memory formation. We speculate that conserved docking modes^56^ between TCR and pMHC make some TCR sequence features generally conducive to activation. TCR-mem aligns with thymic positive selection, extending previous observations that T cells with high self-reactivity (CD5^high^) also have higher reactivity to foreign antigens^57–59^. With TCR-mem, we have precisely identified the TCR sequence determinants of this “peripheral selection”^60^ of the T cell repertoire. For a randomly sampled antigen, T cells with higher TCR-mem are expected to be more effective in generating an immune response.

There are several limitations to our study. First, our study is restricted to ɑβ TCR sequences by virtue of the standard custom primer set for V(D)J amplification. The usage of gamma-delta, rather than ɑβ TCR genes, has clear effects on transcriptional state^61^ that merit further study. Second, our approach to defining a standard set of TCR sequence features excluded T cells in which more than one α or one more than one β chain was detected. We expect dual-alpha chain T cells follow similar relationships between TCR sequence and T cell state, but demonstrating this would require separate analysis approaches. Lastly, the consistency of our TCR scoring functions across different individuals and different antigen specificities does not preclude genetic or antigen-specific effects on T cell differentiation. Future studies should quantify these nuances, as antigen screening technologies gain increasingly higher throughput.

It is now clear that the TCR sequence conveys two types of information: transcriptional fate bias as well as antigen specificity. Both types of information may be crucial to the immune response in autoimmunity, cancer, and infection. TCR features may play a particularly important role in influencing T cell fate for T cells that recognize autoantigens, which should be relatively anergic. The analysis of TCR repertoire data, which generally aims to cluster TCRs by antigen specificity alone, would benefit from integrating information about T cell state. The therapeutic design of TCRs may need to consider not only recognition of the antigenic target, but also differentiation into an effective T cell state.

### Supplementary Note 1

Given the alignment of MAIT and NKT cells on rCCA canonical variate 1 (CV1), we wondered whether the shared transcriptional fate of MAIT and NKT cells was related to biophysical similarity in TCR sequence. We calculated the biophysical distance between canonically-defined MAIT and NKT TCRs as the sum of mean absolute differences in Atchley factors per CDR residue in Dataset 1 (**Methods**). We compared this biophysical distance to background distribution generated by randomly selecting 1000 times a different set of V and J genes as canonical MAIT and TCR genes (**Methods**). Biophysical distance between true MAIT and NKT TCRs fell within this background distribution, indicating that canonical MAIT and NKT TCRs are no more biophysically similar than would be expected by chance (empirical *P* value = 0.22, **Supplementary Figure 7b**). Indeed, two distinct types of TCRs—corresponding to canonical MAIT and NKT sequences – are apparent upon principal component analysis of TCR sequence features within transcriptional cluster A9 (**Supplementary Figure 7c**). From these analyses, we concluded that biophysical similarity in TCR sequence does not explain the shared transcriptional fate of MAIT and NKT cells.

We next examined the shared transcriptional states of MAIT and NKT cells more closely. To construct a MAIT transcriptional state score, we estimated the optimal linear combination of batch-corrected gene expression PC scores to predict canonical MAIT TCR usage via logistic regression in Dataset 1. We followed the same procedure for canonical NKT TCR usage. These two cell state scores demonstrated similar UMAP distributions, concentrated in transcriptional cluster A9 (**Supplementary Figure 7d-f**). Both the MAIT transcriptional score and the NKT transcriptional score were well-described by rCCA CV1 (**Supplementary Figure 7g**), corresponding to increased expression of MAIT and NKT transcripts such as *PLZF* (**Supplementary Figure 7h**), and NKG7 (**Supplementary Figure 7i**). From these analyses, we concluded that MAIT and NKT TCRs share highly similar transcriptional states in peripheral blood, represented by transcriptional cluster A9 in these data.

### Supplementary Note 2

While each T cell in the Dextramer dataset (“Dataset 5”, **Table 1**) should, in theory, recognize only one of pMHC Dextramers, in practice individual cells have non-zero UMI counts for more than one of the 44 pMHC. To stringently remove background Dextramer counts, we fit a negative binomial regression for each of the 44 Dextramers, estimating contributions from technical factors:

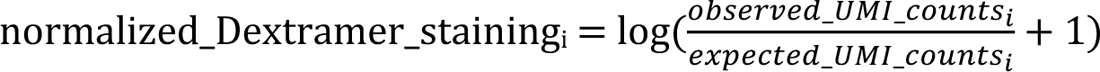

*NCj_UMI_i_* denotes the count of UMIs for negative control (NC) Dextramer j collected for cell i. We include β_TCR_TCR_exp_i_ and β_CD3_CD3_exp_i_ to correct for TCR expression level, as more TCRs expressed at the cell surface should render more opportunities to bind Dextramer. TCR_exp_i_ equals the log(CP10K + 1) normalized expression for CDR3 UMIs (alpha and beta summed) for cell i; CD3_exp_i_ equals the CLR-normalized expression of the CD3 protein for cell i. We include CD8_exp_i_ (CLR-normalized expression of the CD8 protein for cell i) to adjust for the possibility that greater co-receptor expression renders more opportunities to bind Dextramer. We used R package “MASS” (v7.3.54) to fit the negative binomial regressions.

To subtract UMI counts attributable to these technical factors we replaced each cell’s raw Dextramer UMI count with an expression of its negative binomial regression residual:

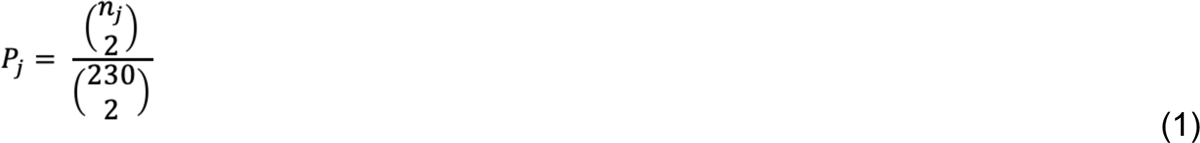

where expected_UMI_counts_i_ equals the negative binomial prediction based on technical factors for cell i. For most Dextramers, we observed a bimodal distribution of normalized Dextramer staining values, distinguishing a background population from an antigen-binding population with Dextramer UMI counts than could be attributed to technical factors. T cells in this putative antigen-specific population were less likely to have other Dextramers with non-zero UMI counts (**Supplementary Figures 19-20**). By careful visual inspection, we set a normalized Dextramer staining threshold for each antigen-specific population to distinguish binders from non-binders (**Supplementary Table 17**).

If T cells assigned to the same antigen specificity based on Dextramer UMI counts really do share antigen specifiity, they should have similar TCR sequences. Thus, we turned to TCR sequences to validate our inferences about antigen specificity. We observed significant TCR sequence conservation within our inferred antigen-specific populations (**Supplementary Figure 21-22, Methods**).

## Methods

### Data acquisition

We acquired published datasets from the following online sources:

**Supplementary Table 1.** Data sources.

| Name | Reference | URL | Data type | File | Date of download |
| --- | --- | --- | --- | --- | --- |
| Dataset 1 | COMBAT<sup>19</sup> | https://doi.org/10.5281/zenodo.6120249 | Sample metadata | CBD-KEY-CLINVAR.tar.gz | 2022-06-23 |
|  |  |  | RNA counts | COMBAT-CITESeq-DATA.h5ad | 2022-06-23 |
|  |  |  | ADT counts | COMBAT-CITESeq-DATA.h5ad | 2022-06-23 |
|  |  |  | TCR | CBD-KEY-CITESEQ-VDJ-T.tar.gz | 2022-06-23 |
| Dataset 2 | Ren et al.<sup>20</sup> | https://www.ncbi.nlm.nih.gov/geo/query/acc.cgi?acc=GSE158055 | Sample metadata | GSE158055_sample_metadata.xlsx | 2021-11-21 |
|  |  |  | RNA counts | GSE158055_covid19_counts.mtx.gz | 2021-11-21 |
|  |  |  | Cell barcodes | GSE158055_covid19_barcodes.tsv.gz | 2021-11-21 |
|  |  |  | Gene names | GSE158055_covid19_features.tsv.gz | 2021-11-21 |
|  |  |  | TCR | GSE158055_covid19_BCR_TCR.tar.gz | 2021-11-21 |
|  |  |  | Cell annotations | GSE158055_cell_annotation.csv.gz | 2021-11-21 |
| Dataset 3 | Stephenson et al.<sup>21</sup> | https://www.ebi.ac.uk/biostudies/arrayexpress/studies/E-MTAB-10026 | Sample metadata | covid_portal_210320_with_raw.h5ad | 2021-03-22 |
|  |  |  | RNA counts | covid_portal_210320_with_raw.h5ad | 2021-03-22 |
|  |  |  | ADT counts | covid_portal_210320_with_raw.h5ad | 2021-03-22 |
|  |  |  | TCR | TCR_merged-Updated | 2023-03-28 |
| Dataset 4 | Domínguez Conde et al.<sup>22</sup><br>*\*only blood and secondary lymphoid tissue* | tissueimmune cellatlas.org | Sample metadata | conde_t-cells.h5ad | 2022-08-02 |
|  |  |  | RNA counts | conde_t-cells.h5ad | 2022-08-02 |
|  |  | European Nucleotide Archive:<br>ftp.sra.ebi.ac.uk/vol1/run/ERR924 | TCR |  | 2022-08-02 |
| Dataset 5 | Boutet et al.<sup>23</sup> | https://www.10xgenomics.com/resources/datasets/cd-8-plus-t-cells-of-healthy-donor-1-1-standard-3-0-2 | Sample metadata | vdj_v1_hs_aggregated_donor[X]/vdj_v1_hs_aggregated_donor[X]_binarized_matrix.csv<br>for [X] 1-4 | 2022-08-30 |
|  |  |  | Dextramer counts | vdj_v1_hs_aggregated_donor[X]/vdj_v1_hs_aggregated_donor[X]_binarized_matrix.csv<br>for [X] 1-4 | 2022-08-30 |
|  |  |  | ADT counts | vdj_v1_hs_aggregated_donor[X]/vdj_v1_hs_aggregated_donor[X]_filtered_feature_bc_matrix.tar.gz<br>for [X] 1-4 | 2022-08-30 |
|  |  |  | RNA counts | vdj_v1_hs_aggregated_donor[X]/vdj_v1_hs_aggregated_donor[X]_filtered_feature_bc_matrix.tar.gz<br>for [X] 1-4 | 2022-08-30 |
|  |  |  | TCR | vdj_v1_hs_aggregated_donor[X]/vdj_v1_hs_aggregated_donor[X]_all_contig_annotations.csv<br>for [X] 1-4 | 2022-08-30 |
| Dataset 6 | Suo et al.<sup>24</sup> | https://developmental.cellatlas.io/fetal-immune | Sample metadata | PAN.A01.v01.raw_count.20210429.NKT.embedding.h5ad | 2022-12-10 |
|  |  |  | RNA counts | PAN.A01.v01.raw_count.20210429.NKT.embedding.h5ad | 2022-12-10 |
|  |  |  | TCR | PAN.A01.v01.raw_count.20210429.NKT.embedding.abTCR.h5ad | 2022-12-10 |

### Sequencing read alignment

Sequencing data were deposited and downloaded as aligned sequencing reads (raw UMI counts for genes, ADTs, or TCR contigs) for Datasets 1-4, and Dataset 6. For TCRs in Dataset 4, we downloaded .fastqs from the European Nucleotide Archive (ENA). We used cellranger vdj version 5.0.1 to align reads to reference GRCh38 5.0.0.

### Quality control and normalization for gene and protein expression

For all single-cell datasets, we removed cells with UMIs for less than 500 genes and cells with greater than 10% of UMIs derived from mitochondrial genes. We then normalized UMIs for each gene in each cell to log(UMI counts per 10,000) and normalized UMIs for each surface protein in each cell via the centered-log-ratio (CLR) transformation.

### Unimodal dimensionality reduction

To identify clusters A1-A9, we applied a unimodal dimensionality reduction pipeline to Dataset 1, using mRNA information alone. After normalizing gene expression, we used the variance-stabilizing transformation (VST) to select the 200 most variable genes in each sample, excluding TCR genes. The union of variable genes across samples totaled 6358 genes. To conduct principal component analysis (PCA), we used R package “irlba” (v2.3.3) on normalized expression of these 6358 genes, each scaled to mean 0 variance 1. We then used Harmony^62^ (R package v1.0) to remove batch effects from the per-cell principal component scores. We applied Harmony to three batch variables: ‘scRNASeq_sample_ID’, ‘Institute’, and ‘Pool_ID.’ We used θ values 1, 0.5 and 0.5 for these batch variables, respectively. We used UMAP to visualize the resultant cell embedding in two dimensions, computed through Symphony^63^ (R package v1.0). We built a shared-nearest-neighbor (SNN) graph of cells based on the first 20 batch-corrected gene expression PCs, using R package “singlecellmethods” (v0.1.0, parameters: prune_snn = 1/25, nn_k = 10, nn_eps = 0.5). To identify transcriptional clusters, we applied Louvain clustering to the SNN graph at multiple resolutions, through the *RunModularityClustering* function from R package “Seurat” (v3.2.2). Clusters A1-A9 were identified by clustering resolution 0.5.

### Multimodal dimensionality reduction

To identify clusters B1-B9, we applied a multimodal dimensionality reduction pipeline^64^ to Dataset 1, incorporating both mRNA and surface protein information. We used the VST to select the 100 most variable genes in each sample, excluding TCR genes. The union of variable genes across samples totaled 4423 genes. We applied Canonical Correlation Analysis (CCA) to paired mRNA and protein measurements, using the expression of these 4423 genes as input matrix 1 and the expression of 10 surface proteins as input matrix 2 (R package “CCA” v1.2.1). To emphasize traditional T cell populations, we selected 10 surface proteins critical to distinguish between CD4, CD8, central memory (CM) and effector memory (EM) T cells (**Supplementary Table 2**). Input gene expression was log10CPK-normalized and scaled to mean 0 variance 1; input protein expression was CLR-normalized and scaled to mean 0 variance 1. As in our unimodal dimensionality reduction pipeline, we then removed batch effects from the per-cell mRNA-based canonical variate (CV) scores, constructed a UMAP and SNN graph based on the 10 batch-corrected CVs, and identified cell clusters. Clusters B1-B9 were identified through clustering at resolution 4.0, and collapsing clusters with similar marker expression (**Supplementary Figure 2q-r**).

### Standardizing T cell states across Datasets

To annotate T cell states in a consistent manner across datasets, we conducted reference mapping with Symphony^63^ (v1.0). Our unimodal T cell state reference (**Supplementary Figure 1**) and multimodal T cell state reference (**Supplementary Figure 2**) were constructed from Dataset 1 (T cells in peripheral blood). To annotate T cell states in Datasets 2-5, we projected cells into these references. Symphony’s reference mapping includes correction for batch variables, which we specified for each Dataset. For all datasets, we corrected for the individual’s donor ID. For Dataset 2, we additionally corrected for batch variables “Sample.type” and “PMID.” For Dataset 3, we additionally corrected for batch variable “Site.” For Dataset 4, we additionally corrected for batch variable “Chemistry,” and conducted Symphony mapping separately for each tissue.

For each projected cell, we used the R package “class” (v7.3.17) to identify the five nearest neighbor cells from Dataset 1. We transferred the majority cell state label among these five to the projected cell.

We did not follow this process for Dataset 6, because prenatal thymic tissue includes progenitor T cell states not observed in peripheral blood. Instead, we used T cell state annotations provided by the authors^24^.

### Quality control for TCR sequence data

TCR sequence data deposited for Dataset 2 included only cells with exactly one TCRɑ chain and exactly one TCRβ chain, with Vɑ, Jɑ, Vβ, and Jβ gene names resolved. In keeping with this quality control by Ren et al., we filtered all other scRNAseq-TCR datasets to include only TCRs with exactly one TCRɑ chain, exactly one TCRβ chain, and Vɑ, Jɑ, Vβ, and Jβ gene names resolved. For analyses purely focused on cell state, such as dimensionality reduction and clustering, we did not apply TCR-based filtering.

### Defining TCR clonotypes

To define the TCR clonotype for each cell, we concatenated the IMGT Vɑ, Jɑ, Vβ, and Jβ gene names with the CDR3ɑ amino acid sequence and CDR3β amino acid sequence. To be considered a “TCR-twin,” TCRs from two different individuals were required to match exactly for each of these TCR components. To be considered part of the same expanded clone (**Supplementary Figure 3**), TCRs had to match exactly for each of these TCR components, and be sampled from the same individual. We recognize that it is optimal to identify expanded clones by nucleotide rather than amino acid sequence, because different ancestral V(D)J nucleotide recombinations can converge to the same amino acid sequence. However, only amino acid sequences were made publicly available for the TCRɑ chain for Dataset 1. In practice, because amino acid convergence is rare, TCR clones are often identified via amino acid sequence^19^.

### TCR twin analysis

To identify “TCR twins” in Dataset 1, we counted the number of individuals observed for each TCR clonotype. The vast majority of TCR clonotypes (234131/234265, 99.9%) were observed in only one individual from Dataset 1, and would be considered “private TCRs.” 134 TCR clonotypes were observed in more than one individual (“public TCRs”), and we analyzed the 115 TCR clonotypes that were observed in exactly two individuals.

We next assigned a transcriptional state to each TCR twin member. 115 TCR clonotypes, each observed in two individuals, implies 230 twin members. For 163/230 twin members (70%), we observed no evidence of clonal expansion (there was only one cell sampled from the individual with the TCR clonotype of interest), and we used the cluster identity of its single cell. For 37 twin members (16%), we observed clonal expansion within a single transcriptional cluster (resolution 0.5, **Figure 1a**), and we used this cluster to annotate the TCR twin member. For 30 twin members (13%), we observed clonal expansion across multiple clusters, and we used the transcriptional cluster that contained the greatest number of constituent cells.

We then counted how many of the 115 TCR twins were assigned the same transcriptional cluster in both individuals. To assess statistical significance, we conducted a binomial test with *N* =115 trials and *P_null_* = the probability of concordant transcriptional clusters by chance. To calculate *P_null_*, we summed the probabilities of randomly drawing two observations with the same cluster assignment for each of the nine transcriptional clusters (A1-A9, **Figure 1a**). For transcriptional cluster *j* with *n_j_* observations, this probability can be computed as:

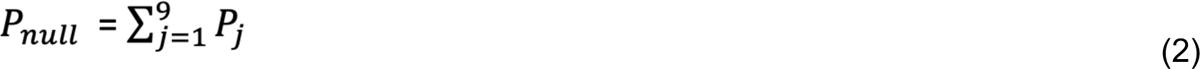

*P_null_* can then be found by summation:

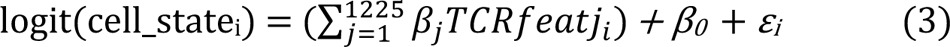

### TCR featurization

Cellranger vdj output provides amino acid sequence for CDR3 regions, but IMGT gene names only for other regions of the TCR. To translate IMGT gene names into CDR1 and CDR2 amino acids, we downloaded amino acid sequences for each gene from the following IMGT sources:

| gene | URL | Date of download |
| --- | --- | --- |
| *TRAV* | https://www.imgt.org/IMGTrepertoire/Proteins/proteinDisplays.php?species=human&latin=Homo%20sapiens&group=TRAV | 03-18-2022 |
| *TRBV* | https://www.imgt.org/IMGTrepertoire/Proteins/proteinDisplays.php?species=human&latin=Homo%20sapiens&group=TRBV | 03-18-2022 |

As this process requires the *TRAV* and *TRBV* gene name to be known, we removed all TCR sequences for which the V gene is unresolved. To focus on functional TCRs, we removed V genes listed as pseudogenes in IMGT. We used TCR position numbers from IMGT to describe the location of each TCR amino acid. To maintain a tractable number of TCR positions, we consider only TCR sequences with CDR3a length ranging from 10 to 17 amino acids and CDR3b length ranging from 11 to 18 amino acids.

To convert each TCR amino acid into a vector of quantitative features, we translated each TCR amino acid into its five Atchley factors^28^. This process results in 290 TCR features, extracted from the 58 TCR amino acid positions that comprise CDR1a, CDR2a, CDR3a, CDR1b, CDR2b, and CDR3b. We also computed the percentage of each CDR loop occupied by each of the 20 amino acids, excluding Glycine as a reference. We also included six TCR features corresponding to the length (in amino acids) of each of the CDR loops. Finally, for each pair of adjacent residues within a TCR chain, we multiplied each possible combination of the five Atchley factors, resulting in 25 interactive TCR features per pair of adjacent residues. The resultant 1225 TCR sequence features is listed in **Supplementary Table 4**. For all analyses, we scaled each TCR feature such that it would have mean 0 and variance 1 in the training dataset. Because the number of amino acids in a TCR sequence varies from cell to cell, shorter TCR sequences contain gaps at some IMGT positions. We fill these entries with the value 0, following the scaling transformation.

### Developing TCR scoring functions

For our purposes, a TCR scoring function takes an amino acid TCR sequence as input and returns a numeric value proportional to the odds of that TCR being observed in the T cell state of interest. This transformation is accomplished by a set of TCR sequence feature weights, which can be learned from our training observations (70% of clones in Dataset 1 and Dataset 2). After training, the TCR sequence feature weights are fixed, and the resultant TCR scoring function is applied to new data (i.e. Datasets 3-6).

We considered three methods to construct TCR scoring functions:

### Method 1: Regularized Canonical Correlation Analysis (rCCA)

By identifying axes of covariation between TCR sequence features and T cell state features across cells, rCCA produces a series of correlated TCR and T cell state scoring functions. rCCA is unsupervised, in that it does not require the analyst to pre-specify T cell states of interest. Instead, it identifies continuous T cell states, which may not exactly align with preexisting T cell state definitions. We applied rCCA via R package “mixOmics” (v6.19.1), tuning ridge penalties via 5-fold cross-validation. For ease of interpretation, we reversed the sign of CV1 scores and CV3 scores.

### Method 2: Logistic Regression with Ridge Penalty

Results from rCCA clearly pointed to four recognizable T cell states (**Figure 2f-i**). rCCA-based TCR scoring functions are optimized to predict the continuous-valued T cell states identified by rCCA, rather than recognizable and reproducible T cell state distinctions. Thus, to enhance biological interpretability, we translated the continuous T cell state values identified by rCCA into binary contrasts: 1) whether the cell belonged to transcriptional cluster A9 (innate-like T), 2) whether the cell belonged to transcriptional clusters B1-B3 (CD8T), 3) whether the cell belonged to the union of transcriptional cluster A5 or transcriptional cluster C15 (CD4^+^ T_reg_ and CD8^+^ T_reg_, respectively, **Supplementary Figure 9**), and 4) whether the cell did not belong to transcriptional clusters B3 or B9 (memory T). We fit a logistic regression to each of these four binary T cell state contrasts:

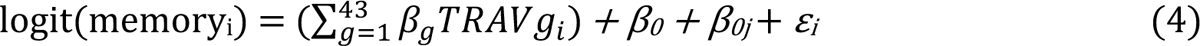

We used the same 1225 TCR sequence features as predictors, and the same set of training observations as in Method 1. To handle collinearity between these TCR sequence features, we implemented ridge regularization via R package “glmnet” (v4.0.2), using 5-fold cross-validation within the training set to select the optimal penalty weight. We removed innate-like cells (transcriptional cluster A9) from regressions focused on CD8T, T_reg_, or memory T cell fate.

### Method 3: Convolutional Neural Network (CNN) Binary Classifier

Methods 1 and 2 only consider linear effects of TCR sequence features and interactive effects limited to adjacent TCR residues. To assess whether nonlinear and/or motif-based TCR sequence feature effects would improve T cell state predictions, we implemented a Convolutional Neural Network (CNN) binary classifier for each of the four T cell state contrasts described above. In the CNN framework, each TCR sequence is represented by a 5 x 58 matrix, corresponding to 5 Atchley factors at each of the 58 residues comprising CDR1a, CDR2a, CDR3a, CDR1b, CDR2b, and CDR3b. After applying convolution and average pooling to this matrix, we concatenate the resultant hidden layer to the TCR score learned through logistic regression. This ensures that the CNN does not have to re-learn linear effects, and can instead focus on potential nonlinear and motif-based effects. We implemented these CNN binary classifiers via python package “torch” (v1.4.0), using the same training and testing split (within Dataset 1 and Dataset 2) as in Method 1 and 2. We used the Adam optimizer and the binary cross-entropy with logits loss function (“BCEWithLogitsLoss” from python package torch), and iterated over a grid of hyperparameters corresponding to batch size, learning rate, and size of hidden layers. For each of these hyperparameter combinations for each of the four cell state contrasts (3 x 2 x 3 x 4 = 72 models), we trained the CNN over 300 epochs and observed minimal impact of hyperparameters on classification performance (AUC, **Supplementary Table 14**) on the Dataset 3, which was never used to fit model parameters. We thus selected hyperparameters corresponding to the least complex model (10 nodes in hidden layer, learning rate 0.0003, and batch size of 256). We used early stopping^65^ to mitigate overfitting, training only until classification performance stopped improving in the held-out testing data (**Supplementary Figure 17a-d**).

### Benchmarking

To benchmark these different TCR scoring functions methods, we applied them to a dataset external to training (Dataset 3), and scaled each type of TCR score to have mean 0 variance 1 prior to association testing via equation (5).

### Applying TCR scoring functions to new data

To apply the TCR scoring functions to new data, we extracted the same 1225 TCR sequence features from the new data, multiplied each TCR feature value by its respective weight learned in from Dataset 1 and 2, and took the sum of these 1225 products. For each TCR feature value in new data, we subtracted its mean value in Datasets 1-2 and divided the result by its standard deviation in Datasets 1-2. Similarly, for each summation over TCR feature value-weight products, we subtracted the Dataset 1-2 mean and divided the result by the Dataset 1-2 standard deviation. Thus, one unit of TCR-mem in any dataset corresponds to one standard deviation of TCR-mem in Datasets 1-2.

### Variance Components Analysis for TCR-innate, TCR-CD8, TCR-reg, and TCR-mem

To estimate the relative contributions of different regions of the TCR to the TCR scoring functions (**Figure 2j-m**), we partitioned the 1225 TCR sequence features into their six complementarity determining loops (CDR1a, CDR2a, CDR3a, CDR1b, CDR2b, CDR3b) and fit a ridge-regularized linear regression model to predict each of the four TCR scores based on each CDR loop (4 TCR scores x 6 CDR loops = 24 regressions). We used the same set of training observations as in Method 1. To handle collinearity between TCR sequence features, we implemented ridge regularization via R package “glmnet” (v4.0.2), using 5-fold cross-validation within the training set to select the optimal penalty weight. This process results in 6 CDR-specific scoring functions for each of the four overall TCR scoring functions. To estimate the variance explained in overall TCR score by each of the CDR-specific TCR scoring functions, we conducted stepwise linear regression, iterating through the CDR regions in the following order: CDR1a, CDR1b, CDR2a, CDR2b, CDR3a, CDR3b. This order attributes all possible variance that can be explained by the V gene region to the V gene region, and attributes remaining contributions to CDR3.

### Association testing between TRAV gene usage and T cell state

To test for associations between *TRAV* gene usage and memory T cell state, we fix a mixed-effects logistic regression:

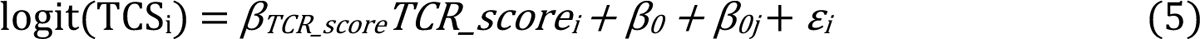

where memoryi equals 0 if cell *i* belongs transcriptional cluster B3 or B9, and 1 otherwise. We removed cells belonging to transcriptional cluster A9 (innate-like). *TRAV*_gi_ equals 1 if cell i expresses *TRAV* gene *g*, and 0 otherwise. We used *TRAV9-2* as a reference. *β0* is the global intercept; *β0j* is a modification to the global intercept fit for each individual *j*. As in our other analyses, we used one cell selected at random to represent each expanded clone. We fit this regression twice; once for all CD8T cells in the training subset of Dataset 1 and Dataset 2, and once for all CD4T cells in the training subset of Dataset 1 and Dataset 2. To assess concordance between *TRAV* gene effects in CD4T versus CD8T cells, we plotted β_g_ estimates for CD8T cells against β_g_ estimates for CD4T cells. We observed two clear outliers with wide standard errors (*TRAV9-1* and *TRAV18*, **Supplementary Table 8**), and removed these prior to estimating the correlation between β_g_ estimates in CD4T compared to CD8T cells.

### Association testing between TCR and T cell state: ꞵ_TCR-innate_, ꞵ_TCR-CD8,_ ꞵ_TCR-reg,_ and ꞵ_TCR-mem_

To test for an association between a TCR score and T cell state (TCS) contrast of interest, we used mixed-effects logistic regression. Using R package “lme4” (v1.1.23), we fit the following regression:

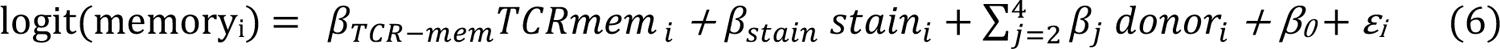

Such that each observation *i* represents one T cell from individual *j*. Consistent with our input to rCCA, we used one cell selected at random to represent each expanded clone. *TCSi* equals 1 if cell *i* is observed in the T cell state of interest; 0 otherwise. *β0* is the global intercept; *β0j* is a modification to the global intercept fit for each individual *j*. *XTCR_score* represents TCR score values, which are standardized to have mean 0 and variance 1 in the training observations from Dataset 1 and Dataset 2.

To estimate *ꞵ_TCR-innate_, ꞵ_TCR-CD8,_ ꞵ_TCR-reg,_ and ꞵ_TCR-mem_* separately for each individual from Dataset 1 and Dataset 2 (**Supplementary Figure 15**), we followed the same procedure with the removal of the *β0j* term. Removing *β0j* reduces the mixed-effects logistic regression to a logistic regression, requiring R package “stats” (v3.6.3) rather than “lme4” for parameter estimation. We included only individuals with at least 100 cells including at least 10 cells that matched the cell state of interest and at least 10 cells that did not match the cell state of interest in the regression. Then, to estimate the distribution of *ꞵ_TCR-innate_* across individuals, we used to these per-individual estimates as input to R function “rma” (from R package “metafor”, v4.0.0), using the maximum likelihood estimator of heterogeneity (method=”ML”).

### Analysis of nonproductive TCRs

To test whether our TCR scoring functions explained T cell state only when applied to the surface-expressed TCR, we analyzed nonproductive TCRs from blood and secondary lymphoid tissue (Dataset 4). We collected all contigs annotated as high confidence, full length, and nonproductive by cellranger vdj (v5.0.1). Many nonproductive TCRs are short in length due to a stop codon; accordingly, we lifted our minimum CDR3 amino acid requirement. Because nonproductive TCRs are lowly expressed, it is uncommon for both a nonproductive alpha chain and a nonproductive beta chain to be detected in the same cell. Thus, to attain statistical power, we did not require cells to have both a nonproductive alpha and a nonproductive beta chain.

Instead, for each of our TCR scoring functions, we created a version that used only TCR alpha chain information and a version that used only TCR beta chain information (4 x 2 = 8 single-chain TCR scoring functions). As before, we used productive TCRs from Dataset 1 and Dataset 2 to train TCR feature weights. We then applied these single-chain TCR scoring functions to nonproductive TCRs from Dataset 4. We removed clonal expansion from Dataset 4 as before, using productive TCR sequences to define T cell clones and selecting one cell at random to represent each expanded clone. We then tested for associations to T cell state as outlined by equation (5). To confirm that lack of association for nonproductive TCRs was not due to statistical power or dataset, we down-sampled productive TCRs from Dataset 4 to match the sample size of nonproductive TCRs and repeated association testing (**Supplementary Figure 19, Supplementary Table 16**).

### Assessing TCR-mem within antigen-specific populations

Inferring antigen specificities from Dextramer UMI counts in Dataset 5 required custom normalization method detailed in **Supplementary Note 2**. Within each group of T cells assigned the same antigen specificity, we fit the following logistic regression (stats v3.6.3):

*Memory_i_* denotes takes the value 1 if T cell *i* is observed in a memory state (clusters B1, B2, B4, **Figure 4a**) and value 0 if T cell *i* is observed in a naïve state (cluster B3, **Figure 4a**). As in previous analyses, we chose one cell at random to represent each expanded clone. To adjust for affinity between the TCR and Dextramer in question, we include *stain_i_* as a covariate, equal to the extent of Dextramer staining following our custom normalization (see **Supplementary Note 2**). For antigen-specific populations that span more than one of the four donors in Dataset 5, we include ∑_j=2_^4^β_j_ donor_i_ to adjust for donor-specific effects. We required that the antigen-specific population have at least 10 distinct TCR clones. Altogether, this process yielded 29 estimates of β_TCR-mem._ for 29 groups of T cells, each specific to a different Dextramer.

We next wanted to understand the typical value of β_TCR-mem_ in any antigen-specific T cell population. With the 29 antigen-specific β_TCR-mem_ estimates and their standard errors, we conducted a random-effects meta-analysis (R package “metafor”, v4.0.0). We used the maximum likelihood estimator of heterogeneity (method=”ML”).

## Data availability

Data analyzed in this study were previously deposited in online databases (**Supplementary Table 1**).

## Code availability

An R package to apply our TCR scoring functions to new data is available at https://github.com/immunogenomics/tcrpheno. Custom analysis scripts for this manuscript are available at https://github.com/immunogenomics/tcrpheno_analysis.

## Acknowledgments

We thank M. Kwun, D.A. Rao and A.H. Jonsson for helpful scientific conversations regarding this work. K.A.L. is supported by award number T32GM007753 from the National Institute of General Medical Sciences. S.R. is supported by NIH grants U19-AI111224-01, P01AI148102-01A1, U01-HG009379-04S1, 1R01AR063759 and UH2-AR067677.

## Ethics declarations

Competing interests

The authors declare no competing interests.

## References

1. Chi, X., Li, Y. & Qiu, X. V(D)J recombination, somatic hypermutation and class switch recombination of immunoglobulins: mechanism and regulation. Immunology 160, 233–247 (2020).

2. Klein, L., Kyewski, B., Allen, P. M. & Hogquist, K. A. Positive and negative selection of the T cell repertoire: what thymocytes see (and don’t see). Nat. Rev. Immunol. 14, 377–391 (2014).

3. Merkenschlager, M. et al. How many thymocytes audition for selection? J. Exp. Med. 186, 1149–1158 (1997).

4. Yun, T. J. & Bevan, M. J. The Goldilocks conditions applied to T cell development. Nature immunology vol. 2 13–14 (2001).

5. Nakayama, T. & Yamashita, M. The TCR-mediated signaling pathways that control the direction of helper T cell differentiation. Semin. Immunol. 22, 303–309 (2010).

6. Hogquist, K. A. & Jameson, S. C. The self-obsession of T cells: how TCR signaling thresholds affect fate “decisions” and effector function. Nat. Immunol. 15, 815–823 (2014).

7. Kaech, S. M. & Cui, W. Transcriptional control of effector and memory CD8+ T cell differentiation. Nat. Rev. Immunol. 12, 749–761 (2012).

8. Pauken, K. E. et al. TCR-sequencing in cancer and autoimmunity: barcodes and beyond. Trends Immunol. 43, 180–194 (2022).

9. Treiner, E. et al. Selection of evolutionarily conserved mucosal-associated invariant T cells by MR1. Nature 422, 164–169 (2003).

10. Lagattuta, K. A. et al. Repertoire analyses reveal T cell antigen receptor sequence features that influence T cell fate. Nat. Immunol. 23, 446–457 (2022).

11. DerSimonian, H., Band, H. & Brenner, M. B. Increased frequency of T cell receptor V alpha 12.1 expression on CD8+ T cells: evidence that V alpha participates in shaping the peripheral T cell repertoire. J. Exp. Med. 174, 1287 (1991).

12. Stadinski, B. D. et al. Hydrophobic CDR3 residues promote the development of self-reactive T cells. Nat. Immunol. 17, 946–955 (2016).

13. Li, H. M. et al. TCRβ repertoire of CD4+ and CD8+ T cells is distinct in richness, distribution, and CDR3 amino acid composition. J. Leukoc. Biol. 99, 505–513 (2016).

14. Carter, J. A. et al. Single T Cell Sequencing Demonstrates the Functional Role of αβ TCR Pairing in Cell Lineage and Antigen Specificity. Front. Immunol. 10, 1516 (2019).

15. Daley, S. R. et al. Cysteine and hydrophobic residues in CDR3 serve as distinct T-cell self-reactivity indices. J. Allergy Clin. Immunol. 144, 333–336 (2019).

16. Kasatskaya, S. A., et al. Functionally specialized human CD4+ T-cell subsets express physicochemically distinct TCRs. Elife 9, (2020).

17. Zhang, Z., Xiong, D., Wang, X., Liu, H. & Wang, T. Mapping the functional landscape of T cell receptor repertoires by single-T cell transcriptomics. Nat. Methods 18, 92–99 (2021).

18. Schattgen, S. A. et al. Integrating T cell receptor sequences and transcriptional profiles by clonotype neighbor graph analysis (CoNGA). Nat. Biotechnol. 40, 54–63 (2022).

19. COvid-19 Multi-omics Blood ATlas (COMBAT) Consortium. Electronic address: & COvid-19 Multi-omics Blood ATlas (COMBAT) Consortium. A blood atlas of COVID-19 defines hallmarks of disease severity and specificity. Cell 185, 916–938.e58 (2022).

20. Ren, X. et al. COVID-19 immune features revealed by a large-scale single-cell transcriptome atlas. Cell 184, 5838 (2021).

21. Stephenson, E. et al. Single-cell multi-omics analysis of the immune response in COVID-19. Nat. Med. 27, 904–916 (2021).

22. Domínguez Conde, C., et al. Cross-tissue immune cell analysis reveals tissue-specific features in humans. Science 376, eabl5197 (2022).

23. Boutet, S. C. et al. Scalable and comprehensive characterization of antigen-specific CD8 T cells using multi-omics single cell analysis. The Journal of Immunology 202, 131–134 (2019).

24. Suo, C. et al. Mapping the developing human immune system across organs. Science 376, eabo0510 (2022).

25. Nathan, A. et al. Single-cell eQTL models reveal dynamic T cell state dependence of disease loci. Nature 606, 120–128 (2022).

26. Venturi, V., Price, D. A., Douek, D. C. & Davenport, M. P. The molecular basis for public T-cell responses? Nat. Rev. Immunol. 8, 231–238 (2008).

27. Lu, T. et al. Deep learning-based prediction of the T cell receptor-antigen binding specificity. Nat Mach Intell 3, 864–875 (2021).

28. Atchley, W. R., Zhao, J., Fernandes, A. D. & Drüke, T. Solving the protein sequence metric problem. Proc. Natl. Acad. Sci. U. S. A. 102, 6395–6400 (2005).

29. Tilloy, F. et al. An Invariant T Cell Receptor α Chain Defines a Novel TAP-independent Major Histocompatibility Complex Class Ib–restricted α/β T Cell Subpopulation in Mammals. J. Exp. Med. 189, 1907–1921 (1999).

30. Porcelli, S., Morita, C. T. & Brenner, M. B. CDlb restricts the response of human CD4−8−T lymphocytes to a microbial antigen. Nature 360, 593–597 (1992).

31. Miescher, G. C., Howe, R. C., Lees, R. K. & MacDonald, H. R. CD3-associated alpha/beta and gamma/delta heterodimeric receptors are expressed by distinct populations of CD4-CD8-thymocytes. J. Immunol. 140, 1779–1782 (1988).

32. Lantz, O. & Bendelac, A. An invariant T cell receptor alpha chain is used by a unique subset of major histocompatibility complex class I-specific CD4+ and CD4-8-T cells in mice and humans. J. Exp. Med. 180, 1097–1106 (1994).

33. Teh, H. S. et al. Thymic major histocompatibility complex antigens and the alpha beta T-cell receptor determine the CD4/CD8 phenotype of T cells. Nature 335, 229–233 (1988).

34. Li, J., et al. KIR+CD8+ T cells suppress pathogenic T cells and are active in autoimmune diseases and COVID-19. Science 376, eabi9591 (2022).

35. Elhanati, Y., Sethna, Z., Callan, C. G., Jr, Mora, T. & Walczak, A. M. Predicting the spectrum of TCR repertoire sharing with a data-driven model of recombination. Immunol. Rev. 284, 167–179 (2018).

36. Rubelt, F. et al. Individual heritable differences result in unique cell lymphocyte receptor repertoires of naïve and antigen-experienced cells. Nat. Commun. 7, 11112 (2016).

37. Stephens, M. False discovery rates: a new deal. Biostatistics 18, 275–294 (2017).

38. Zeldovich, K. B., Berezovsky, I. N. & Shakhnovich, E. I. Protein and DNA sequence determinants of thermophilic adaptation. PLoS Comput. Biol. 3, e5 (2007).

39. Egorov, E. S. et al. The Changing Landscape of Naive T Cell Receptor Repertoire With Human Aging. Front. Immunol. 9, 1618 (2018).

40. Janeway, C. A., Jr, Travers, P. & Walport, M. The rearrangement of antigen-receptor gene segments controls lymphocyte development.: The Immune System … (2001).

41. Manfras, B. J., Terjung, D. & Boehm, B. O. Non-productive human TCR β chain genes represent VDJ diversity before selection upon function: insight into biased usage of TCRBD and TCRBJ genes and diversity of CDR3 region length. Hum. Immunol. 60, 1090–1100 (1999).

42. Zhang, W. et al. A framework for highly multiplexed dextramer mapping and prediction of T cell receptor sequences to antigen specificity. Sci Adv 7, (2021).

43. Chiang, D. et al. Single-cell profiling of peanut-responsive T cells in patients with peanut allergy reveals heterogeneous effector TH2 subsets. J. Allergy Clin. Immunol. 141, 2107– 2120 (2018).

44. Rajasekaran, K. et al. Tetramer-aided sorting and single-cell RNA sequencing facilitate transcriptional profiling of antigen-specific CD8+ T cells. Transl. Oncol. 27, 101559 (2023).

45. Solouki, S. et al. TCR Signal Strength and Antigen Affinity Regulate CD8+ Memory T Cells. J. Immunol. 205, 1217–1227 (2020).

46. Rock, K. L., Reits, E. & Neefjes, J. Present Yourself! By MHC Class I and MHC Class II Molecules. Trends Immunol. 37, 724–737 (2016).

47. Van Rhijn, I., Godfrey, D. I., Rossjohn, J. & Moody, D. B. Lipid and small-molecule display by CD1 and MR1. Nat. Rev. Immunol. 15, 643–654 (2015).

48. Scott-Browne, J. P., White, J., Kappler, J. W., Gapin, L. & Marrack, P. Germline-encoded amino acids in the alphabeta T-cell receptor control thymic selection. Nature 458, 1043– 1046 (2009).

49. Deibel, M. R., Jr et al. Expression of terminal deoxynucleotidyl transferase in human thymus during ontogeny and development. J. Immunol. 131, 195–200 (1983).

50. Marcou, Q., Mora, T. & Walczak, A. M. High-throughput immune repertoire analysis with IGoR. Nat. Commun. 9, 561 (2018).

51. Berzins, S. P., Cochrane, A. D., Pellicci, D. G., Smyth, M. J. & Godfrey, D. I. Limited correlation between human thymus and blood NKT cell content revealed by an ontogeny study of paired tissue samples. Eur. J. Immunol. 35, 1399–1407 (2005).

52. Martin, E. et al. Stepwise development of MAIT cells in mouse and human. PLoS Biol. 7, e54 (2009).

53. Leeansyah, E., Loh, L., Nixon, D. F. & Sandberg, J. K. Acquisition of innate-like microbial reactivity in mucosal tissues during human fetal MAIT-cell development. Nat. Commun. 5, 3143 (2014).

54. López-Sagaseta, J. et al. The molecular basis for Mucosal-Associated Invariant T cell recognition of MR1 proteins. Proc. Natl. Acad. Sci. U. S. A. 110, E1771–8 (2013).

55. Patel, O. et al. Recognition of vitamin B metabolites by mucosal-associated invariant T cells. Nat. Commun. 4, 2142 (2013).

56. Zareie, P. et al. Canonical T cell receptor docking on peptide–MHC is essential for T cell signaling. Science 372, eabe9124 (2021).

57. Ernst, B., Lee, D. S., Chang, J. M., Sprent, J. & Surh, C. D. The peptide ligands mediating positive selection in the thymus control T cell survival and homeostatic proliferation in the periphery. Immunity 11, 173–181 (1999).

58. Mandl, J. N., Monteiro, J. P., Vrisekoop, N. & Germain, R. N. T cell-positive selection uses self-ligand binding strength to optimize repertoire recognition of foreign antigens. Immunity 38, 263–274 (2013).

59. Fulton, R. B. et al. The TCR’s sensitivity to self peptide--MHC dictates the ability of naive CD8+ T cells to respond to foreign antigens. Nat. Immunol. 16, 107 (2015).

60. Rocha, B. & von Boehmer, H. Peripheral selection of the T cell repertoire. Science 251, 1225–1228 (1991).

61. Parker, M. E. & Ciofani, M. Regulation of γδ T Cell Effector Diversification in the Thymus. Front. Immunol. 11, 42 (2020).

62. Korsunsky, I. et al. Fast, sensitive and accurate integration of single-cell data with Harmony. Nat. Methods 16, 1289–1296 (2019).

63. Kang, J. B. et al. Efficient and precise single-cell reference atlas mapping with Symphony. Nat. Commun. 12, 5890 (2021).

64. Nathan, A. et al. Multimodally profiling memory T cells from a tuberculosis cohort identifies cell state associations with demographics, environment and disease. Nat. Immunol. 22, 781–793 (2021).

65. Amari, S., Murata, N., Muller, K. R., Finke, M. & Yang, H. H. Asymptotic statistical theory of overtraining and cross-validation. IEEE Trans. Neural Netw. 8, 985–996 (1997).

