## Supplementary Figures for "The T cell receptor sequence influences the likelihood of T cell memory formation"

### Supplementary Figure 1.

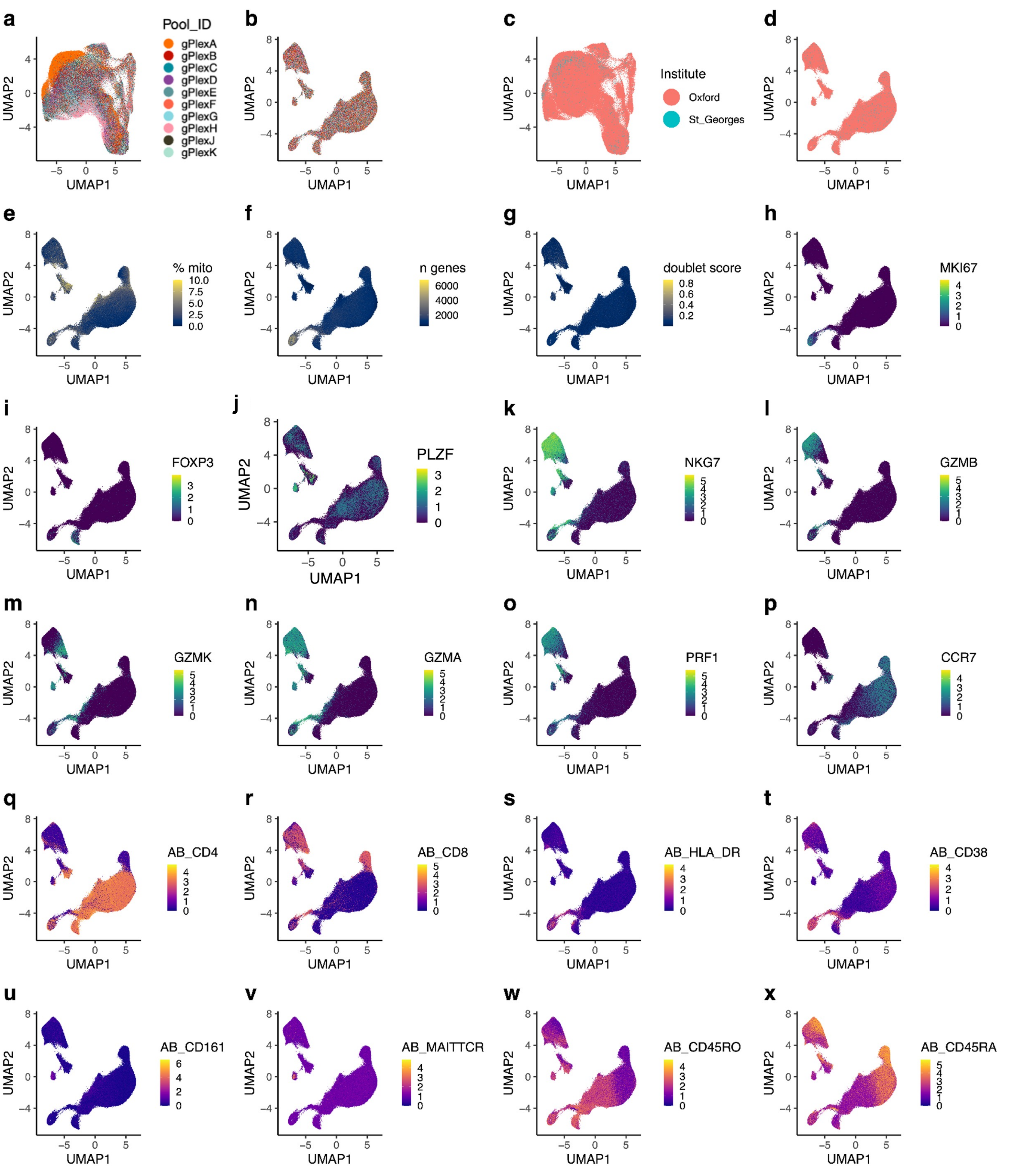


**Supplementary Figure 1. (a)** UMAP of Dataset 1 T cells based on the first 20 principal components (PCs) of gene expression, prior to batch correction. Pool_ID indicates sequencing library plex (Fluidigm Cell-ID 20-Plex Pd Barcoding Kit). **(b)** Dataset 1 T cells colored as in (a), now arranged in a UMAP following batch-correction by Harmony. **(c)** Dataset 1 T cells prior to batch correction, colored by hospital recruitment site. **(d)** Dataset 1 T cells following batch correction, colored by hospital recruitment site. **(e)** Dataset 1 T cells colored by percentage of UMIs aligned to mitochondrial transcripts. **(f)** Dataset 1 T cells colored by the number of unique genes with nonzero counts. **(g)** Dataset 1 T cells colored by Scrublet doublet scores. **(h) - (p)** Dataset 1 T cells colored by log(CP10K + 1) normalized expression of marker transcripts. **(q) - (x)** Dataset 1 T cells colored by centered-log-ratio (CLR) normalized TotalSeq UMIs.

### Supplementary Figure 2

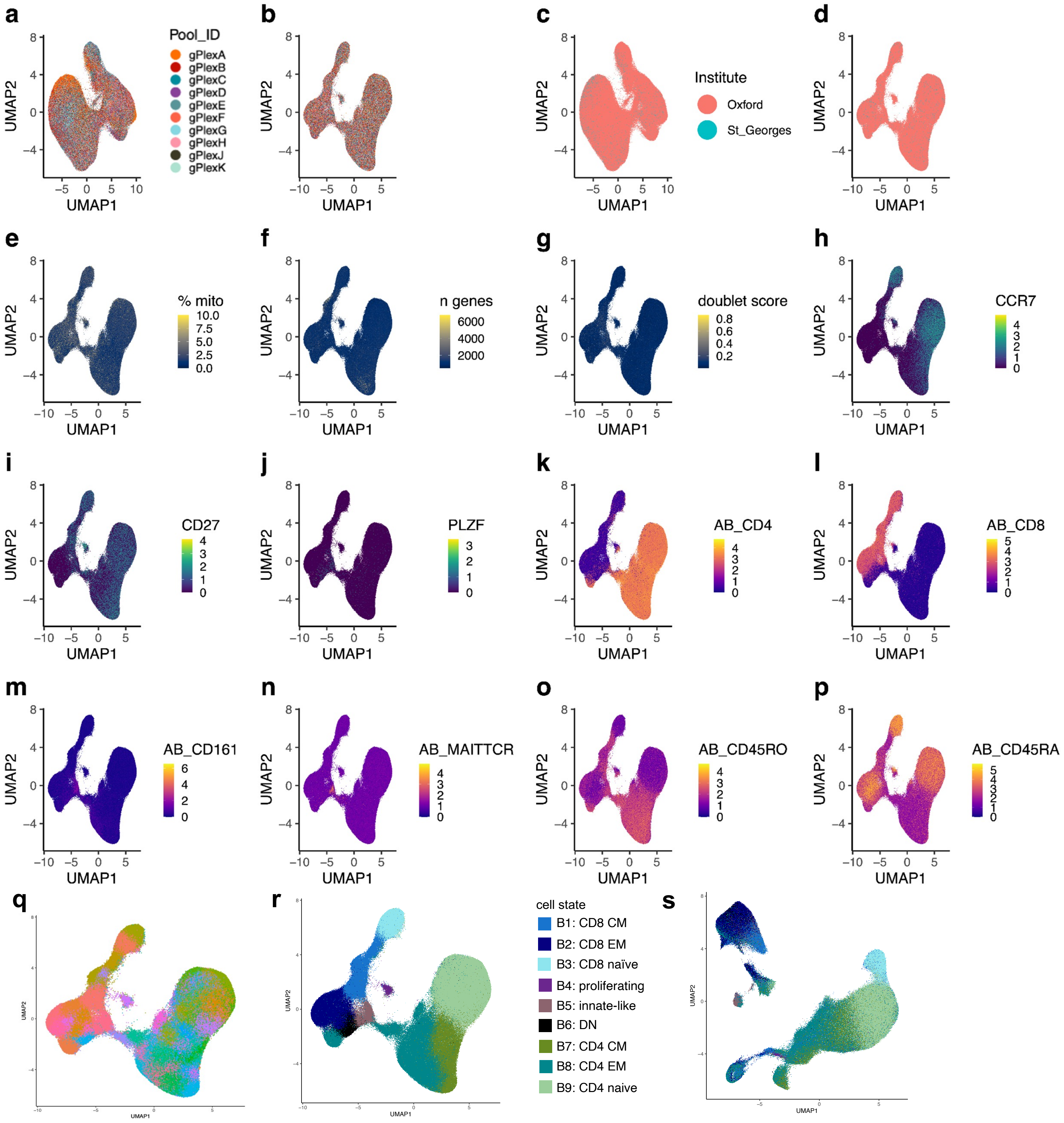


**Supplementary Figure 2. (a)** UMAP of Dataset 1 T cells, based on the first 10 gene expression-based canonical variates from canonical correlation analysis (CCA) applied to the scaled and normalized expression of 4423 variable genes and 10 surface proteins (Supplementary Table 1) relevant to CD4, CD8, central memory (CM), and effector memory (EM) distinctions (Methods). Pool_ID indicates sequencing library plex (Fluidigm Cell-ID 20-Plex Pd Barcoding Kit). **(b)** Dataset 1 T cells colored as in (a), following batch-correction by Harmony. **(c)** UMAP as in (a), colored by hospital recruitment site. **(d)** UMAP as in (b), colored by hospital recruitment site. **(e)** Dataset 1 T cells colored by percentage of UMIs aligned to mitochondrial transcripts. **(f)** Dataset 1 T cells colored by the number of unique genes with nonzero counts. **(g)** Dataset 1 T cells colored by Scrublet doublet scores. **(h) - (j)** Dataset 1 T cells colored by log(CP10K + 1) normalized UMI counts of marker transcripts. **(k) - (p)** Dataset 1 T cells colored by centered-log-ratio (CLR) normalized expression of TotalSeq UMIs. **(q)** Dataset 1 T cells assigned to 60 discrete clusters by Louvain clustering, implemented via Seurat::RunModularityClustering at resolution 4.0. **(r)** 60 Louvain clusters from (q) collapsed into 9 T cell state annotations (B1-B9). **(s)** Dataset 1 T cells colored by annotations B1-B9, rearranged into their original protein-agnostic UMAP (Supplementary Figure 1a).

#### Supplementary Figure 3

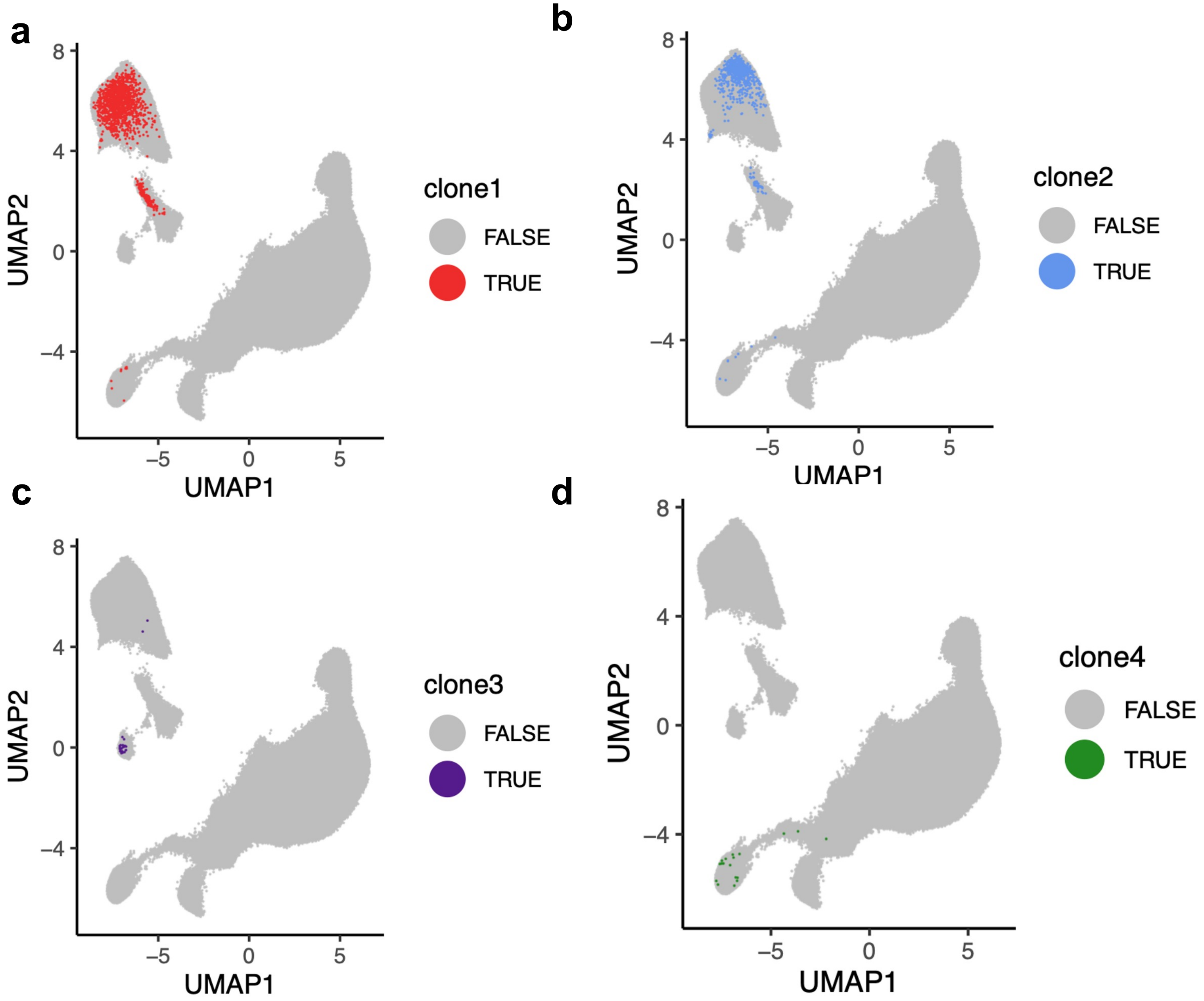


**Supplementary Figure 3. (a)** UMAP of T cells from Dataset 1, colored red if they belong to clone1 ("TRAV3*01 TRA_CAVRKLIF TRAJ4*01 -TRBV15*01 TRB_CATSRGQGRGVETQYF TRBJ2-5*01" from donor H00067) and grey otherwise. **(b)** UMAP of T cells from Dataset 1, colored blue if they belong to clone2 ("TRAV30*05 TRA_CGTDYRRDNYGQNFVF TRAJ26*01 -TRBV6-5*01 TRB_CASSYGGGPTEAFF TRBJ1-1*01" from donor S00045) and grey otherwise. **(c)** UMAP of T cells from Dataset 1, colored purple if they belong to clone3 ("TRAV10*01 TRA_CVVSALGGNNRLAF TRAJ7*01 -TRBV27*01 TRB_CASSPALSPEAFF TRBJ1-1*01" from donor H00070) and grey otherwise. **(d)** UMAP of T cells from Dataset 1, colored green if they belong to clone4 ("TRAV19*01 TRA_CALSEGSYNTDKLIF TRAJ34*01 -TRBV14*01 TRB_CASSQDRRTQYF TRBJ2-3*01" from donor N00006) and grey otherwise.

### Supplementary Figure 4

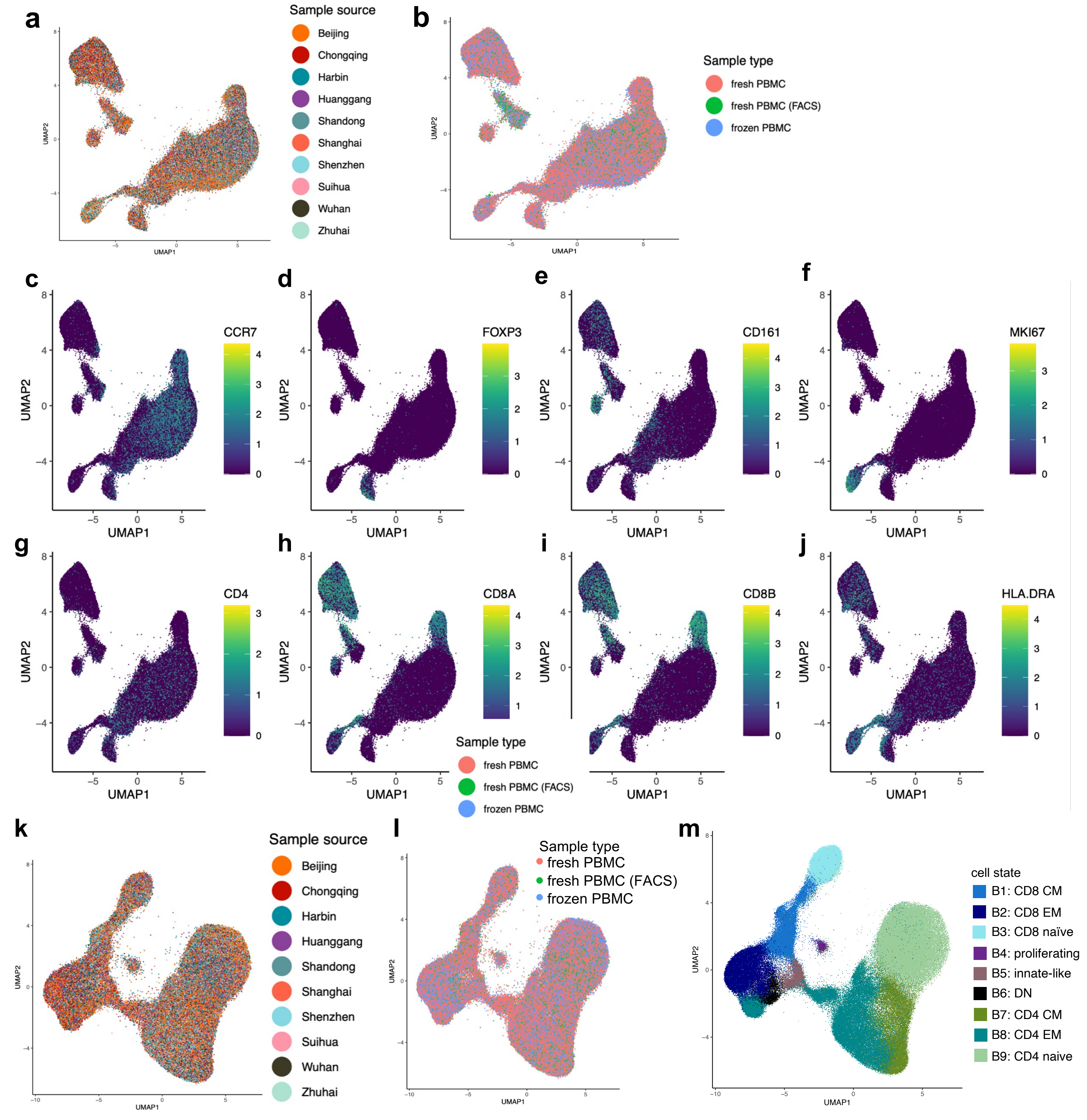


**Supplementary Figure 4. (a-b)** Dataset 2 T cells, projected into reference A (Supplementary Figure 1) by Symphony. Successful batch correction is evident by integration of cells from different collection sites (a) and different sample processing types (b). **(c-j)** Dataset 2 T cells projected into reference A, colored by log(CP10K+1) normalized expression of marker transcripts. Gene expression values from Dataset 2 T cells align with gene expression values from Dataset 1 T cells (Supplementary Figure 1h-p), demonstrating successful harmonization of Dataset 1 and Dataset 2. **(k)** T cells from Dataset 2, projected into reference B (Supplementary Figure 2) by Symphony, colored by sample source. **(l)** Dataset 2 T cells as in (k), colored by Sample type. **(m)** Dataset 2 T cells projected into reference B, colored by reference B annotations assigned via k-nearest-neighbors (k=5)

#### Supplementary Figure 5

**a**

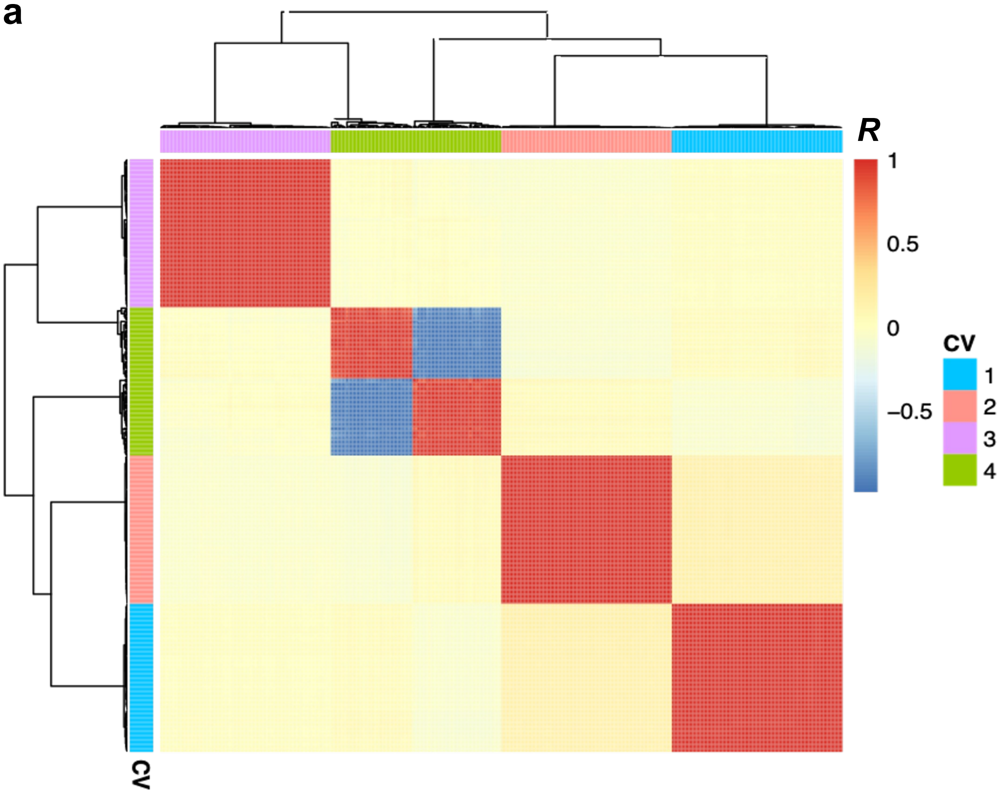


**b**

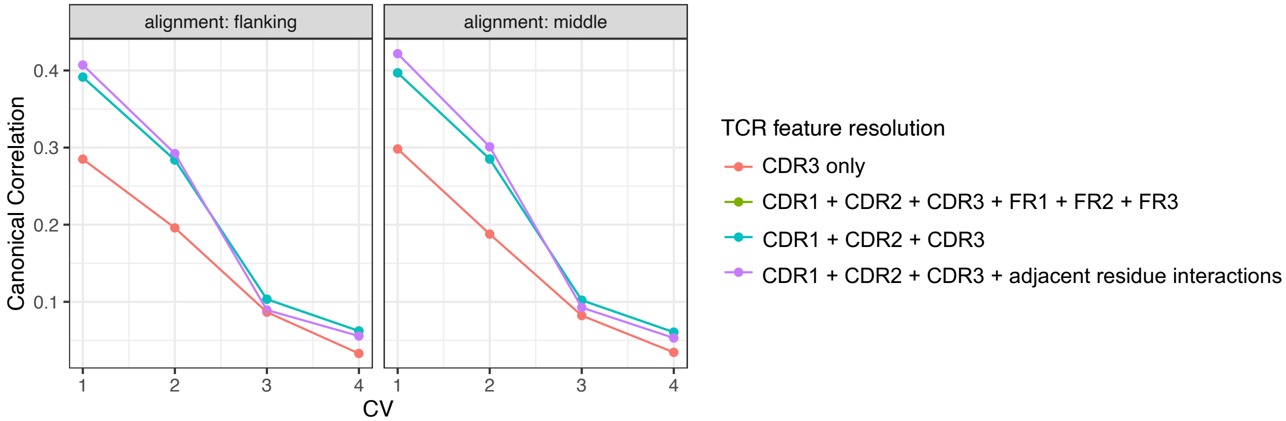


**Supplementary Figure 5. (a)** To assess whether our rCCA solution was sensitive to the choice of representative cell per T cell clone, we re-selected one cell at random to represent each expanded T cell clone and re-ran regularized Canonical Correlation Analysis (rCCA) 1000 times. We computed correlations between each pair of re-runs of rCCA with respect to the 1225 TCR feature loadings for canonical variates (CVs) 1-4. Heatmap shows correlation between re-run i (row) and re-run j (column). Dendograms depict hierarchical clustering of rows and columns through R package “pheatmap.” Pearson’s *R* values near 1 and -1 for all CV-matched comparisons indicate that rCCA applied to these data is robust to choice of representative cell per T cell clone. **(b)** To align residues from Complimentary Determining Region 3 (CDR3) sequences of different lengths, we considered two schemes: “flanking” and “middle.” For the “flanking” scheme, shorter CDR3 sequences align to the flanking residues of longer CDR3 sequences. For the “middle” scheme, shorter CDR3 sequences align to the middle residues of longer CDR3 sequences. We tested the two CDR3 alignment schemes, as well as four different TCR feature resolutions, by applying rCCA to Dataset 1 and examining the resultant canonical correlations. We observed the strongest canonical correlations when we included CDR1, 2, and 3 from both the alpha and beta chain, excluded framework (FR) residues, and included interactions between adjacent CDR3 residues (see Methods).

#### Supplementary Figure 6

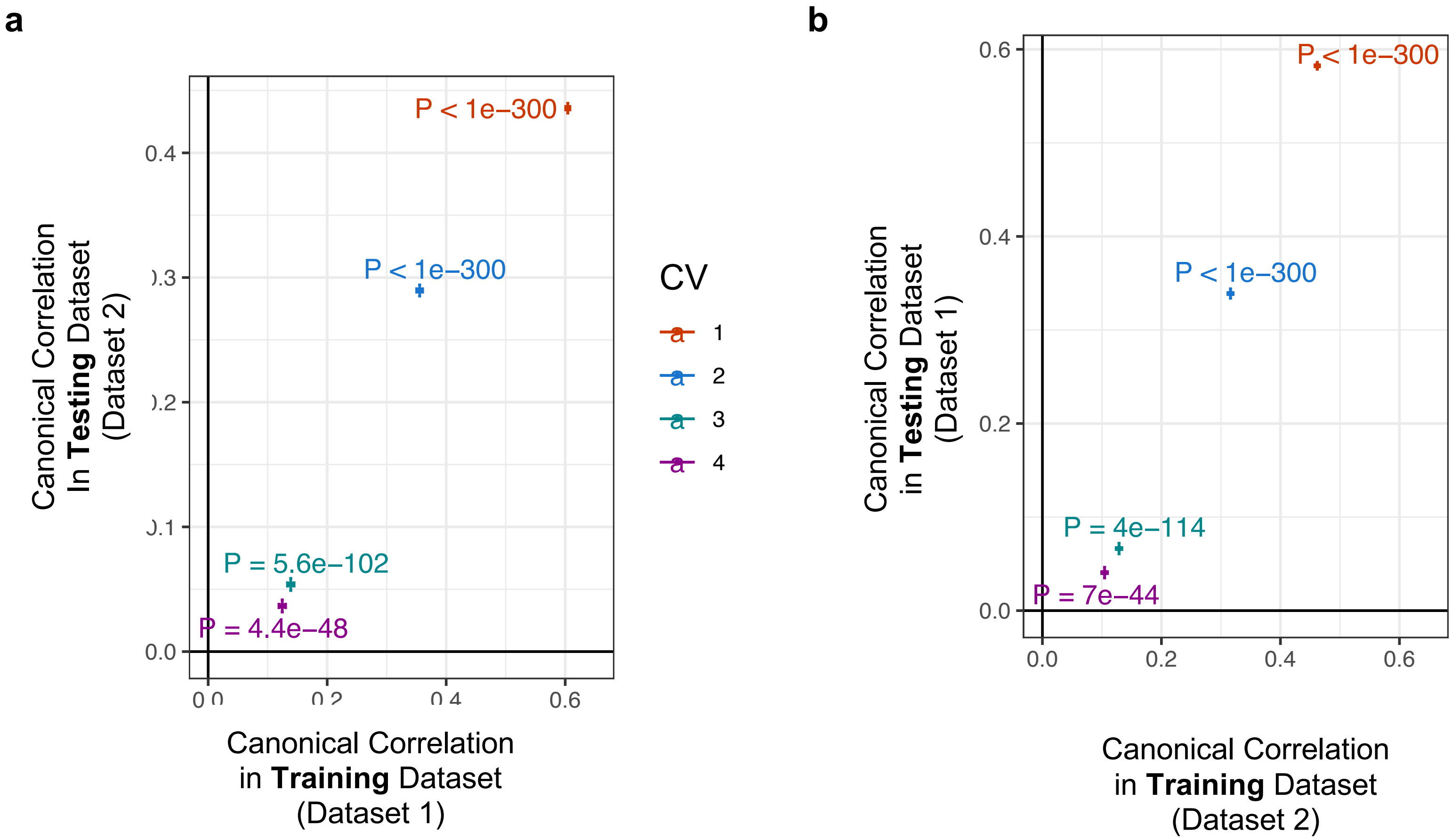


**Supplementary Figure 6. (a)** We used rCCA to learn weights on TCR features and T cell state features in Dataset 1, and then used these weights to score cells in Dataset 2. For each CV, we observed a significant correlation between the TCR-based score and the TCS-based score across cells in Dataset 2. *P* value computed by Pearson's product=moment correlation test (two-sided). **(b)** Analogous to **(a)**, such that weights are learned using Dataset 2 and tested in Dataset 1.

### Supplementary Figure 7

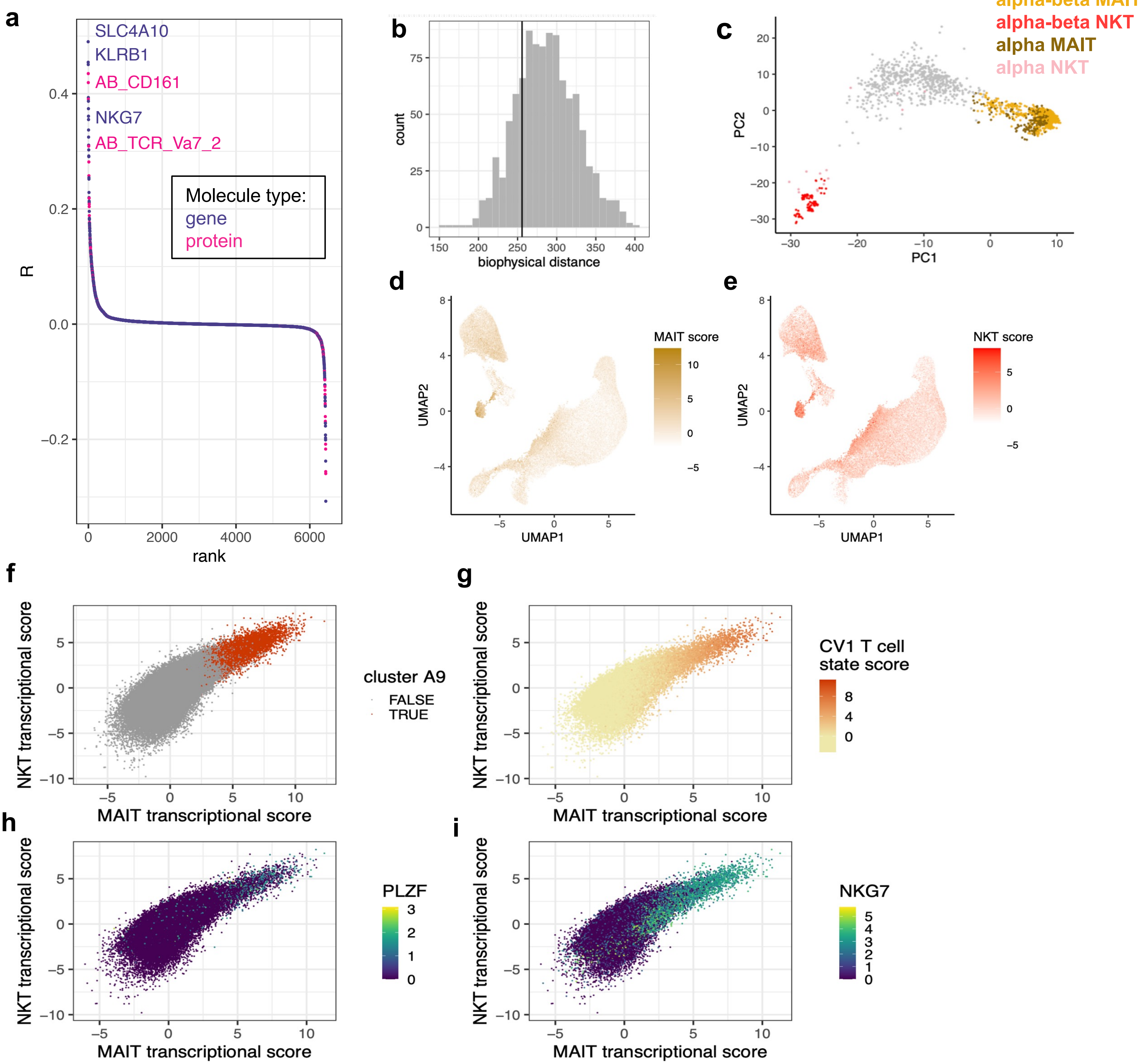


**Supplementary Figure 7. (a)** Correlations between CV1 T cell state score and expression of each variable gene (purple) as well as each TotalSeq surface protein (pink) in Dataset 1. **(b)** Histogram depicts distribution of biophysical distance values observed through 1000 random samples of Dataset TCRs that are not canonical MAIT and NKT TCRs (see Supplementary Note 1). Black vertical line denotes biophysical distance between canonical MAIT and NKT TCRs in Dataset 1. **(c)** To examine the TCR composition of T cells in cluster A9, we conducted principal component (PC) analysis with TCR features, restricted to cells in cluster A9 (Methods). PC1 and PC2 scores suggested three types of TCRs in cluster A9, two of which corresponded to canonical MAIT and NKT TCR genes, respectively. **(d-e)** To examine the transcriptional state of MAIT-like TCRs and NKT-like TCRs separately, we fit two logistic regressions: one to find the optimal linear combination of batch-corrected gene expression PC scores for canonical MAIT TCR usage, the other for canonical NKT TCR usage (Methods). MAIT transcriptional scores (d) are fitted values from the first model, while NKT transcriptional scores (e) are fitted values from the second model. **(f)** MAIT transcriptional scores (x-axis) and NKT transcriptional scores (y-axis) for T cells in Dataset 1. As expected from their similar patterns in (d-e), MAIT transcriptional score and NKT transcriptional score are highly correlated, and both are maximal in transcriptional cluster A9. **(g)** Dataset 1 T cells as in (f). CCA CV1 tracks closely with both the MAIT gene expression phenotype and the NKT gene expression phenotype. **(h)** Dataset 1 T cells as in (f). log(CP10K +1) *PLZF* expression increases with the MAIT, NKT, and CV1 transcriptional scores. **(i)** Dataset 1 T cells as in (f). log(CP10K +1) *NKG7* expression increases with the MAIT, NKT, and CV1 transcriptional scores.

#### Supplementary Figure 8

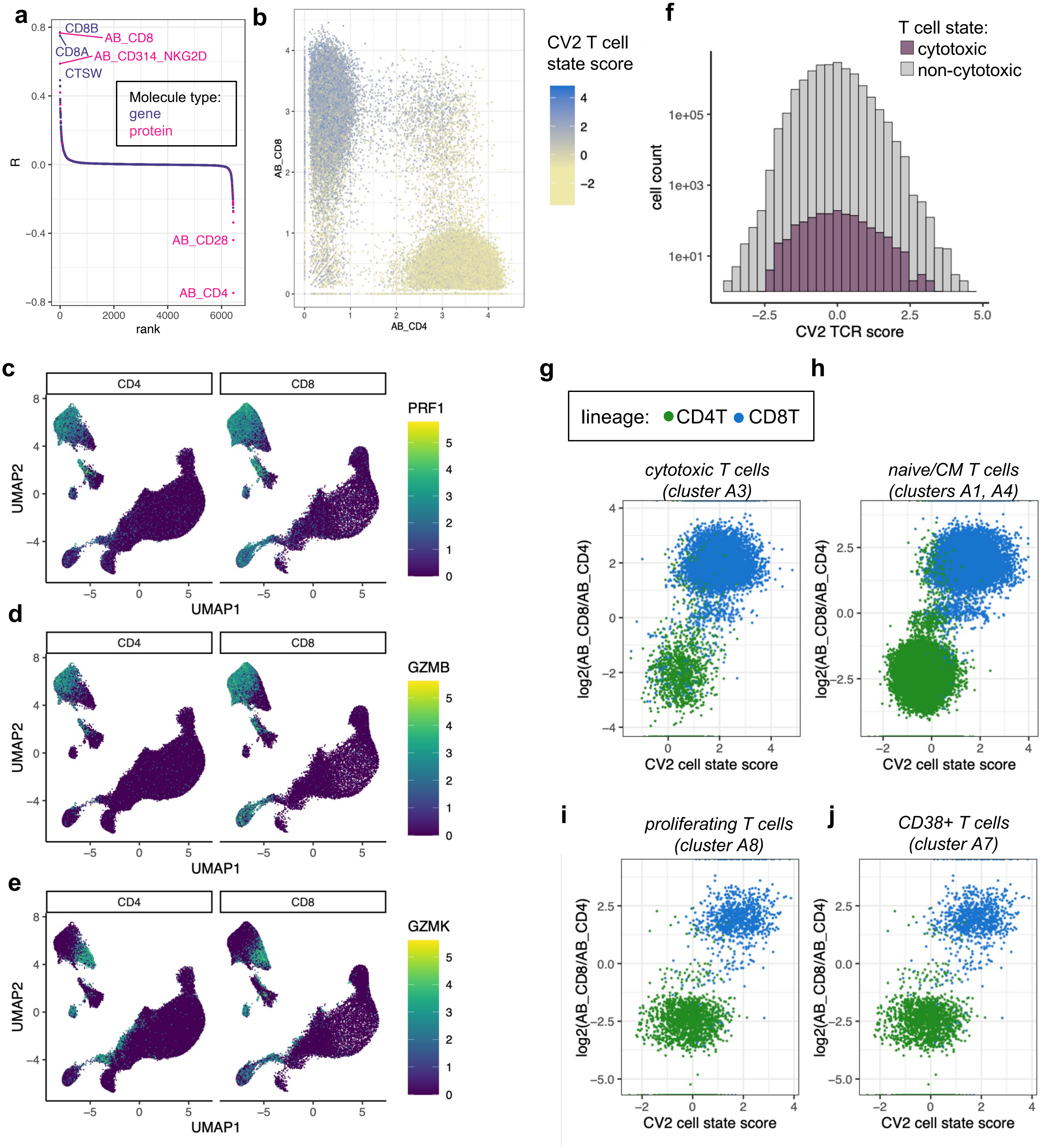


**Supplementary Figure 8. (a)** Correlations between CV2 T cell state score and expression of each variable gene (purple) as well as each TotalSeq surface protein (pink) in Dataset 1. **(b)** T cells from Dataset 1, plotted by CLR-normalized UMI counts for CD4 (x-axis) and CD8 (y-axis) TotalSeq antibodies. The CV2 T cell state score is lower in CD4 compared to CD8 T cells. **(c)** T cells from Dataset 1, colored by log(CP10K + 1) normalized expression of *PRF1*, separated into CD4 (left) and CD8 (right) populations (CD4 versus CD8 lineage annotated via clusters B1-B9, see Methods). **(d-e)** Dataset 1 T cells as in (c), for other cytotoxicity markers *GZMB* and *GZMK*. These markers confirmed that both CD4 and CD8 cytotoxic T cells localized to transcriptional cluster A3. **(f)** Among CD4 T cells, CV2 TCR scores were roughly equivalent between cytotoxic and non-cytotoxic cells. **(g)** Among cytotoxic T cells, the CV2 cell state score delineates CD4 and CD8 populations (CD4 versus CD8 lineage annotated via clusters B1-B9, see Methods). **(h-j)** Dataset 1 T cells as in (g), for naive/CM, proliferating, and CD38+ T cells, respectively.

### Supplementary Figure 9

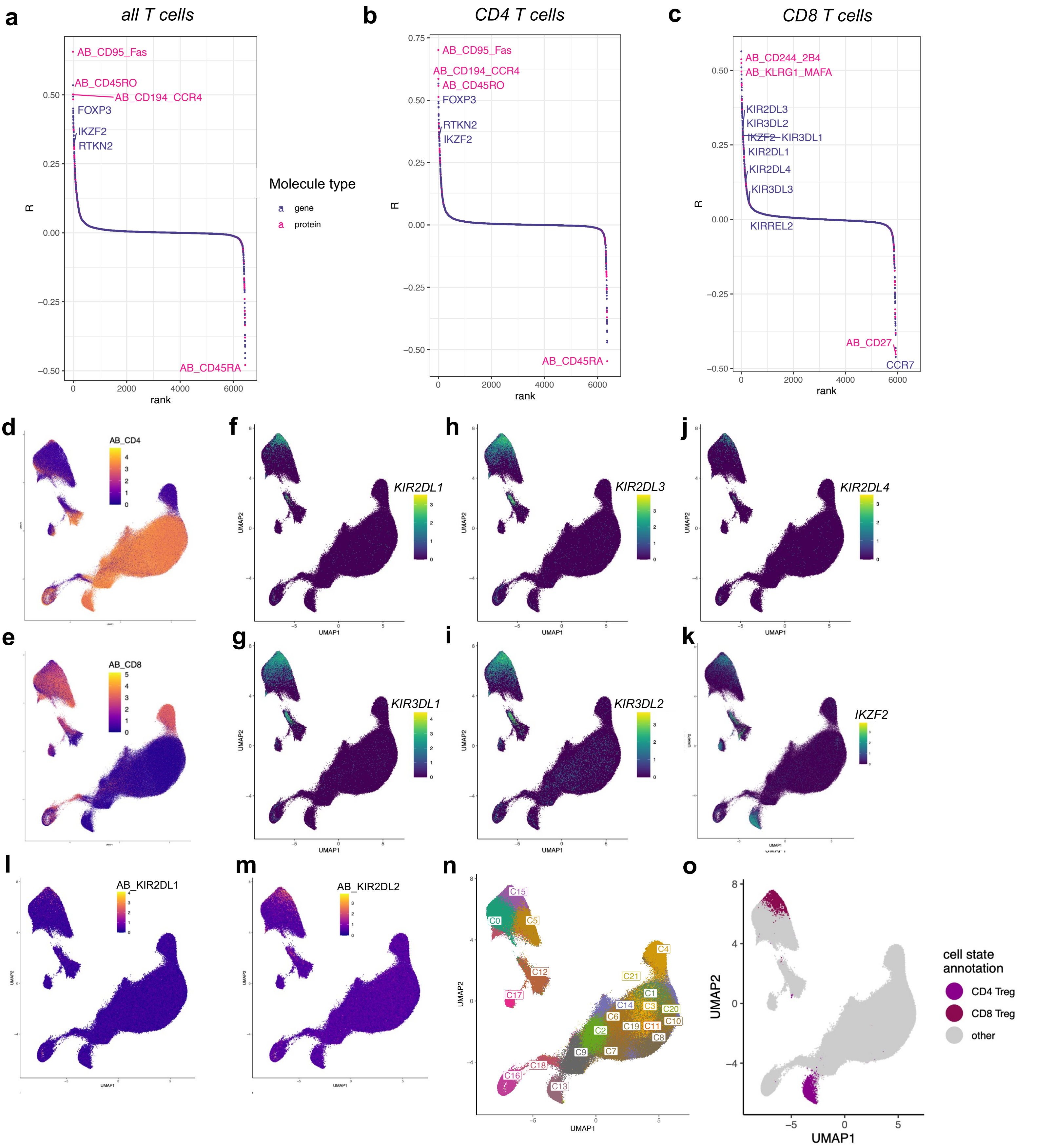


**Supplementary Figure 9. (a)** Correlations between CV3 T cell state score and expression of each variable gene (purple) as well as each TotalSeq surface protein (pink) in Dataset 1. **(b)** Correlations as in (a), restricted to CD4 T cells. **(c)** Correlations as in (a), restricted to CD8 T cells. **(d-e)** UMAP of Dataset 1 T cells, colored by CLR-normalized protein expression of CD4 and CD8, respectively (TotalSeq antibodies). **(f-k)** UMAP of Dataset 1 T cells, colored by log(CP10K +1) normalized expression of select transcripts. **(l-m)** UMAP of Dataset 1 T cells, colored by CLR-normalized protein expression of KIR2DL1 and KIR2DL2, respectively (TotalSeq antibodies). **(n)** Louvain clustering of Dataset 1 T cells at resolution 2.0. **(o)** Cell annotations with respect to Treg state for Dataset 1 T cells. Cells were labeled “CD8 Treg” if they belonged to cluster 15 (SFig 9n) and “CD4 Treg” if they belonged to cluster A6 (Fig 1a).

#### Supplementary Figure 10

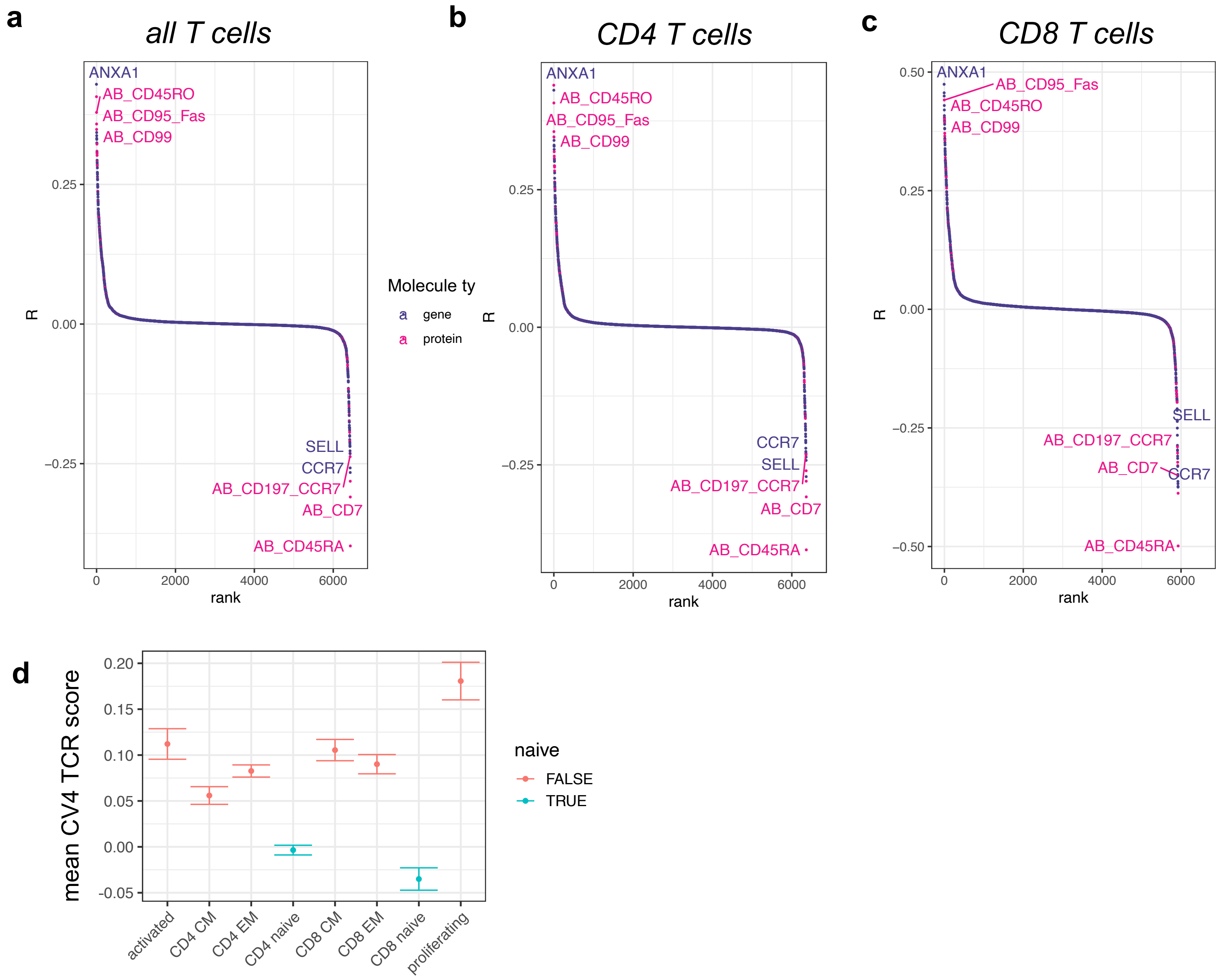


**Supplementary Figure 10. (a)** Correlations between CV4 T cell state score and expression of each variable gene (purple) as well as each TotalSeq surface protein (pink) in Dataset 1. **(b)** Correlations as in (a), restricted to CD4 T cells. **(c)** Correlations as in (a), restricted to CD8 T cells. **(d)** Mean CV4 TCR score across T cell populations in Dataset 1. Error bars denote 95% confidence intervals.

#### Supplementary Figure 11

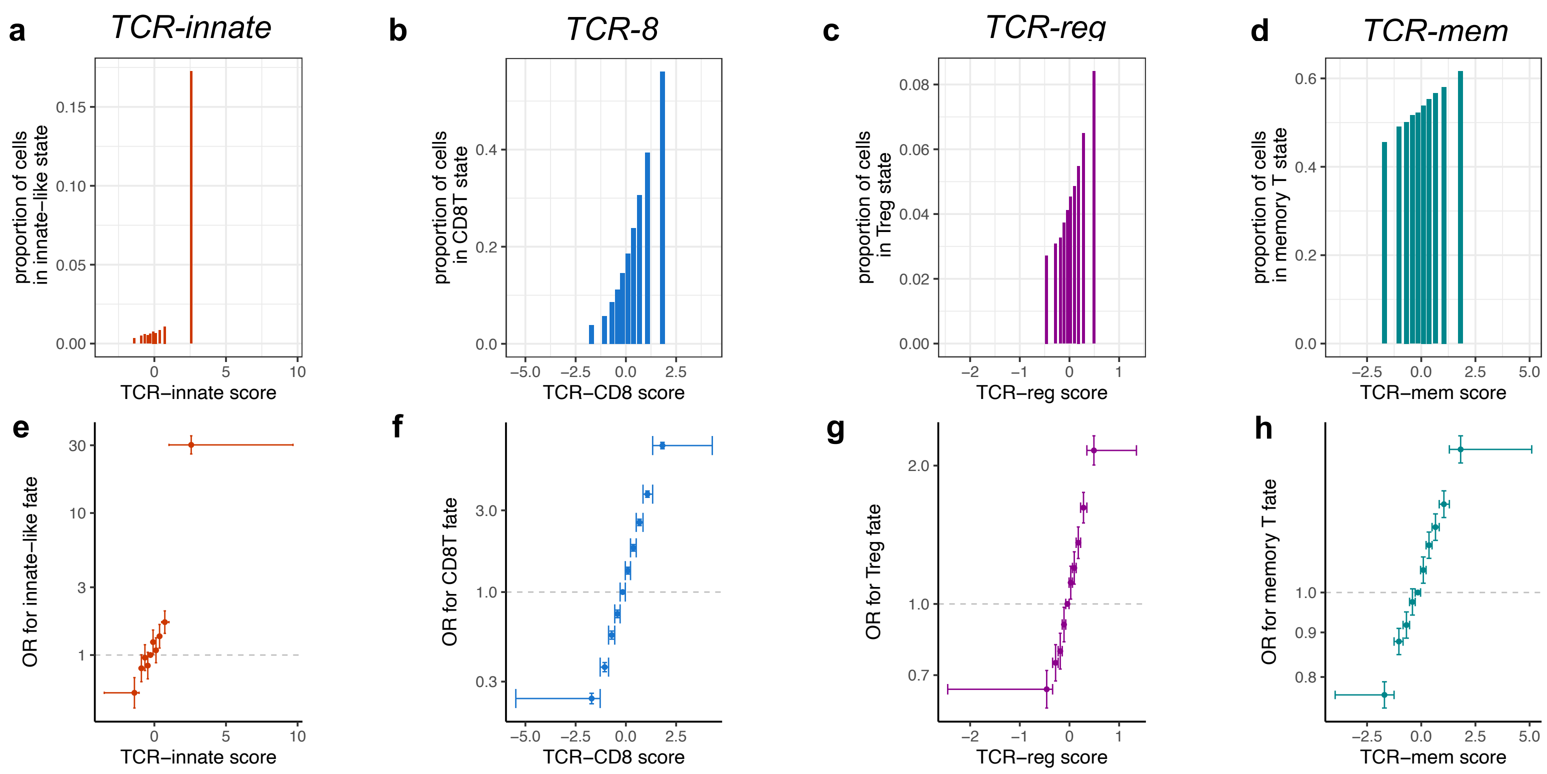


**Supplementary Figure 11. (a-d)** Proportion of T cell clones from the training subset of Dataset 1 and Dataset 2 observed in each T cell state of interest within each decile of its corresponding TCR score. **(e)** Each point represents a decile of the TCR-innate score in the training subset of Dataset 1 and Dataset 2, with a horizontal bar spanning from its minimum to maximum value. We compute the odds ratio (OR, y-axis) for *PLZF*$^{high}$ T cell state for T cells in each decile compared to T cells in the fifth decile. 95% CIs (error bars) and *P* values computed via mixed-effects logistic regression. **(f-h)** T cells as in (e), for CD8T, Treg, and memory T cell states, and their corresponding TCR scores.

#### Supplementary Figure 12

amino acid category:
hydrophobic
hydrophobic/aromatic
basic
acidic
neutral

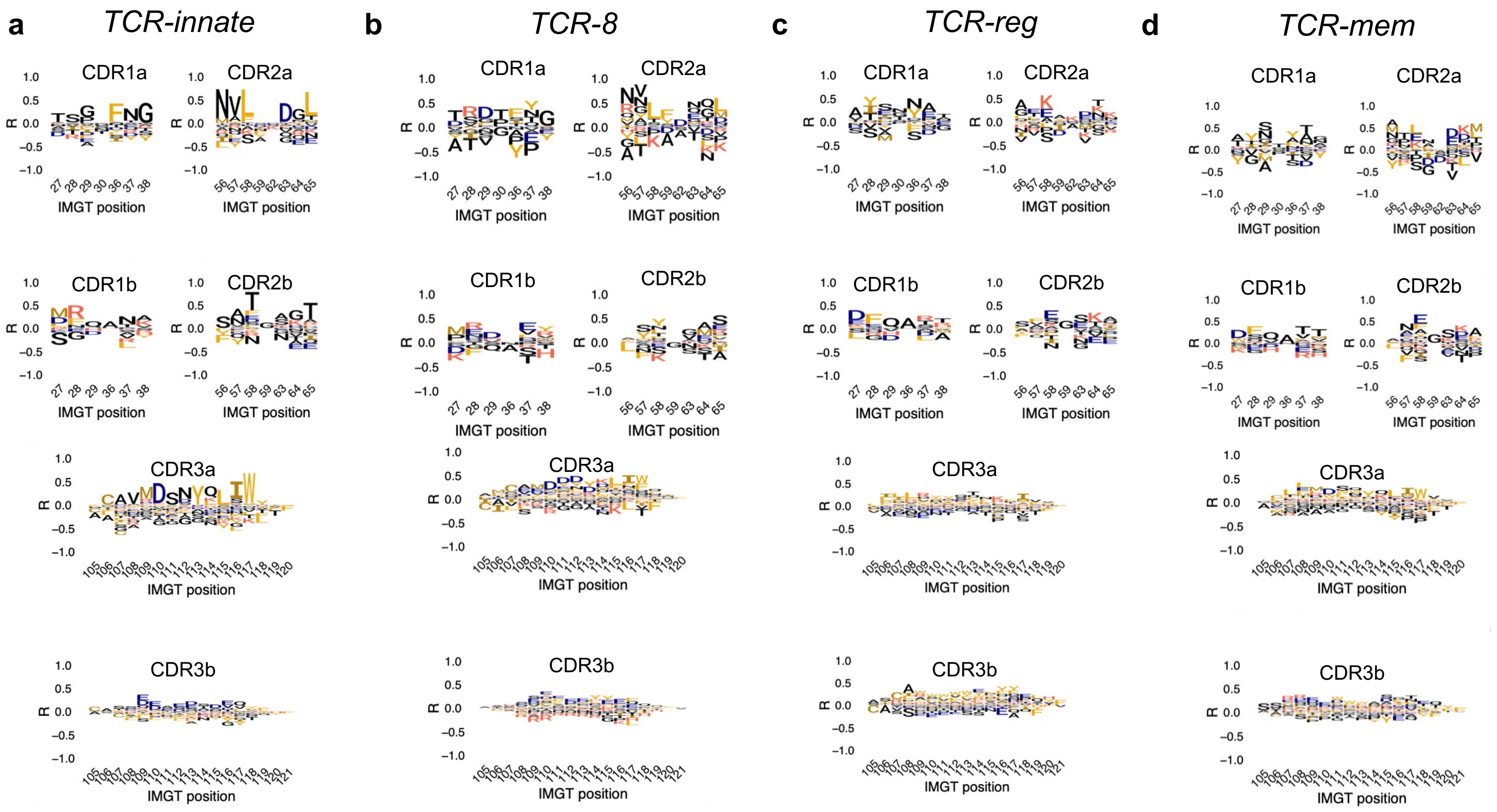


**Supplementary Figure 12.** TCR features for **(a)** TCR-innate, **(b)** TCR-CD8, **(c)** TCR-reg, and **(d)** TCR-mem, visualized as marginal correlations to each amino acid in each complementarity-determining region (CDR). Specifically, using training observations from Dataset 1 and Dataset 2, we computed the correlation between each TCR score value and the presence (=1) or absence (=0) of each amino acid in each position of the TCR.

### Supplementary Figure 13

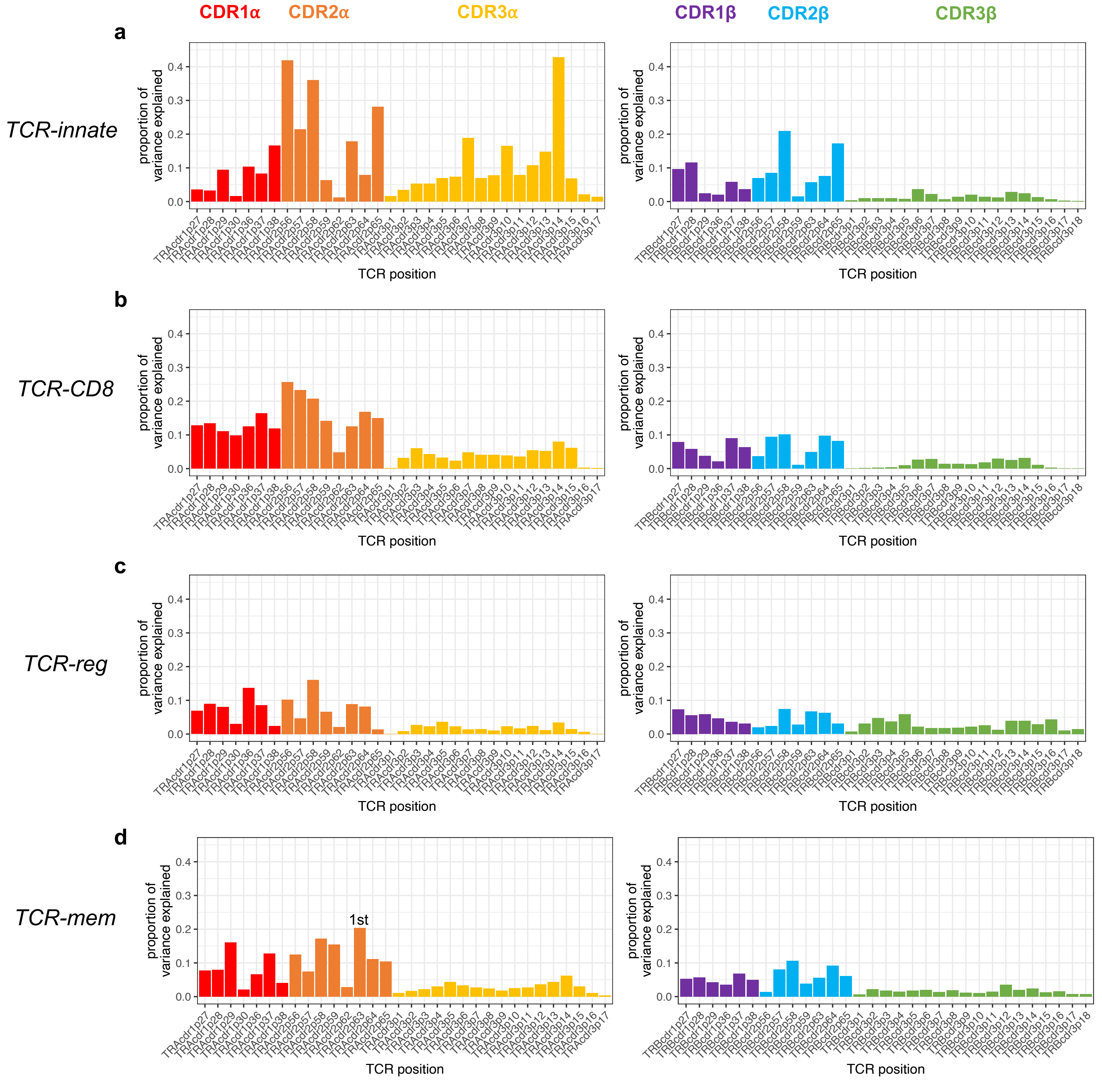


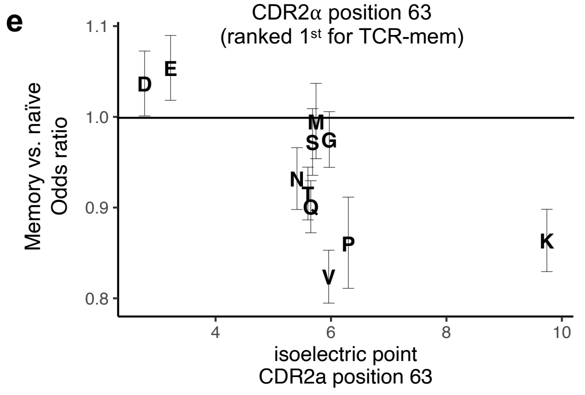


**Supplementary Figure 13.** To investigate the relative contributions of each TCR position to each TCR scoring function, we quantified variance explained through linear regression. Specifically, we used the training observations from Datasets 1 and 2 to fit a linear model for each TCR score at each TCR position. TCR score was the response variable, and amino acid identity was the categorical predictor variable. A "gap" was included as a possible amino acid value. Each bar denotes the variance explained by one such linear model for **(a)** TCR-innate, **(b)** TCR-CD8, **(c)** TCR-reg, and **(d)** TCR-mem. **(e)** We used mixed-effects logistic regression to estimate the effect of each amino acid at TCRa p63. Memory versus naïve state was the response variable. The reference group was T cells with an amino acid gap at TCRa p63. For each amino acid observed at TCRa position 63, we plot the odds ratio for memory state compared to this reference group. Error bars denote 95% confidence intervals.

### Supplementary Figure 14

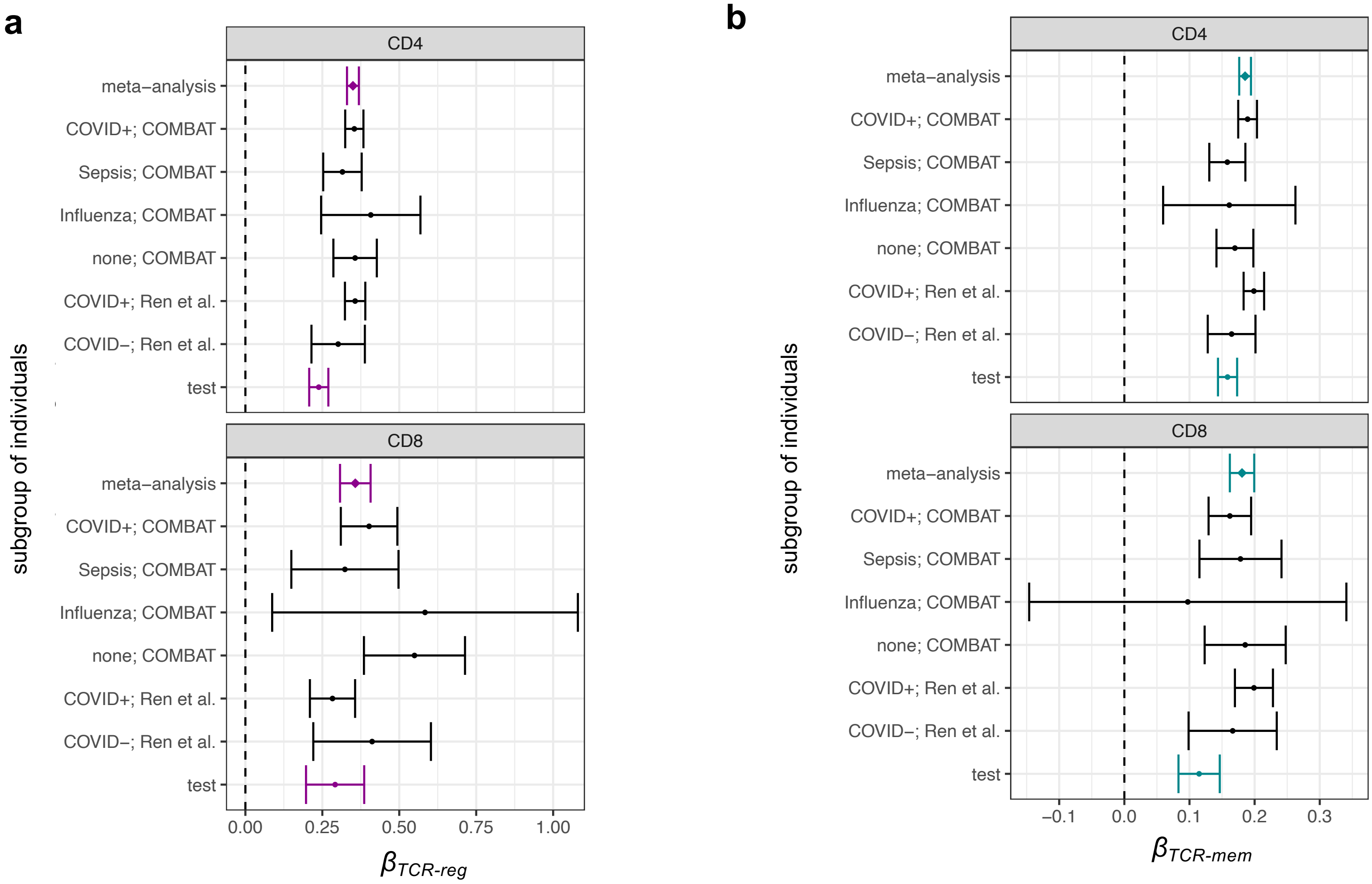


**Supplementary Figure 14. (a)** Forest plot depicting the association between TCR-reg and Treg state for T cells from each subset of individuals, further stratified by CD4 lineage (top) and CD8 lineage (bottom). CD4 lineage and CD8 lineage are designated via clusters B1-B9 (Supplementary Figure 2). **(b)** Forest plot depicting association between TCR-mem and memory state for T cells stratified as in a. $\beta_{TCR\text{-}reg}$ and $\beta_{TCR\text{-}mem}$ are computed via mixed-effects logistic regression, with the TCR score scaled to mean 0 variance 1 in the training set. Error bars denote 95% confidence intervals. The “test” subset is the 66 individuals from Dataset 1 and Dataset 2 selected at random to be held out from TCR score training. Meta-analytic $\beta_{TCR\text{-}reg}$ and $\beta_{TCR\text{-}mem}$ are estimated by fixed-effects inverse-variance-weighted meta-analysis, applied to the 6 training subsets.

### Supplementary Figure 15

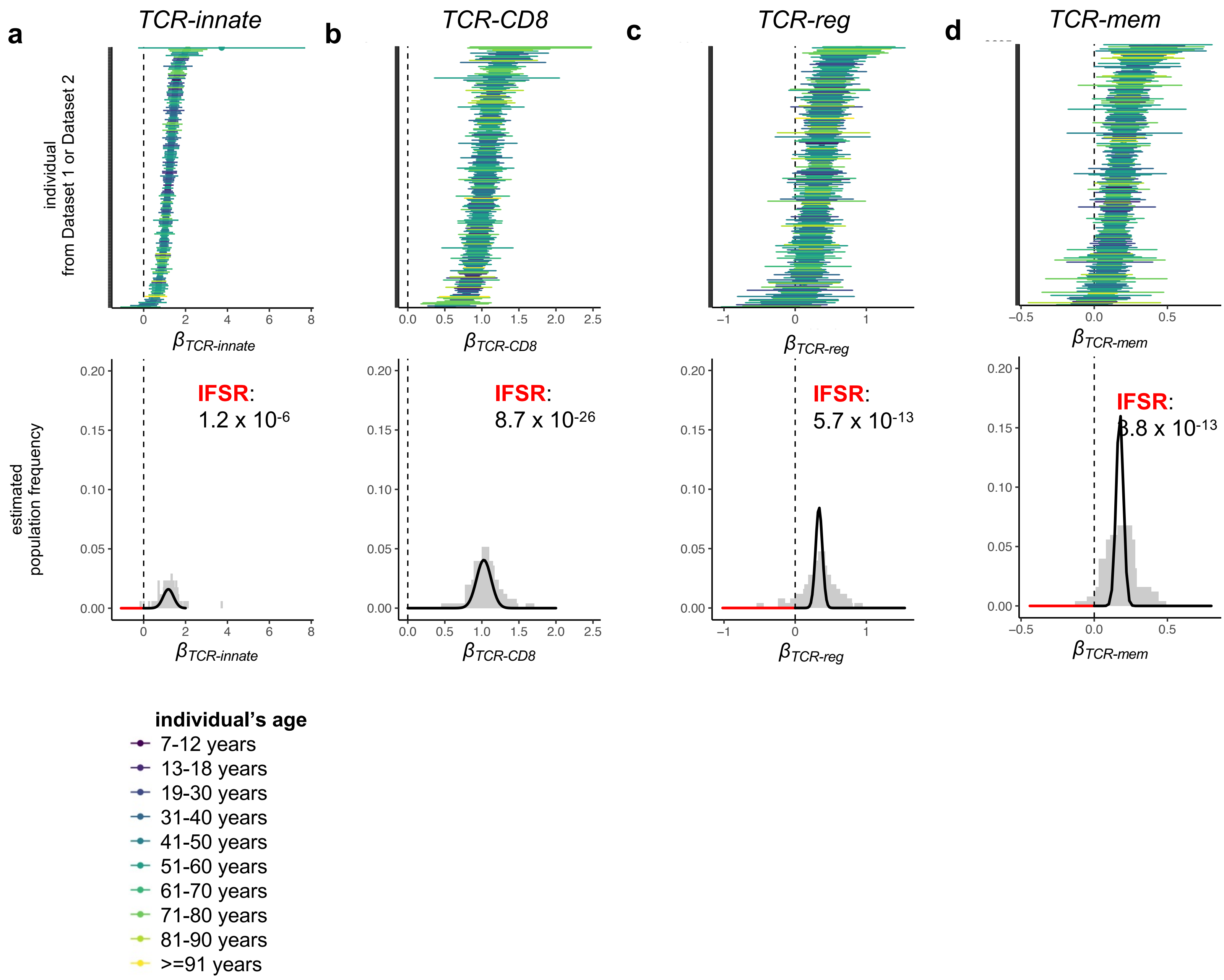


**Supplementary Figure 15. (a)** Forest plot above shows 95% CIs for $\beta_{TCR\text{-}innate}$ for Dataset 1 and Dataset 2 T cells stratified by individual. $\beta_{TCR\text{-}innate}$ estimates are computed via logistic regression for innate-like transcriptional fate, done separately in each individual. Histogram below shows the same $\beta_{TCR\text{-}innate}$ estimates in gray, with the estimated distribution of $\beta_{TCR\text{-}innate}$ overlaid in black. Parameters for the distribution of $\beta_{TCR\text{-}innate}$ are estimated via random effects meta-analysis, implemented by R package metafor. Red-shaded area corresponds to the local false sign rate (lFSR), the estimated proportion of individuals for whom TCR-innate does not demonstrate a positively signed association to innate-like transcriptional fate. **(b)** Analogous to (a), for TCR-CD8 and CD8 T cell fate. **(c)** Analogous to (a), for TCR-reg and Treg cell fate. **(d)** Analogous to (a), for TCR-mem and memory T cell state.

### Supplementary Figure 16

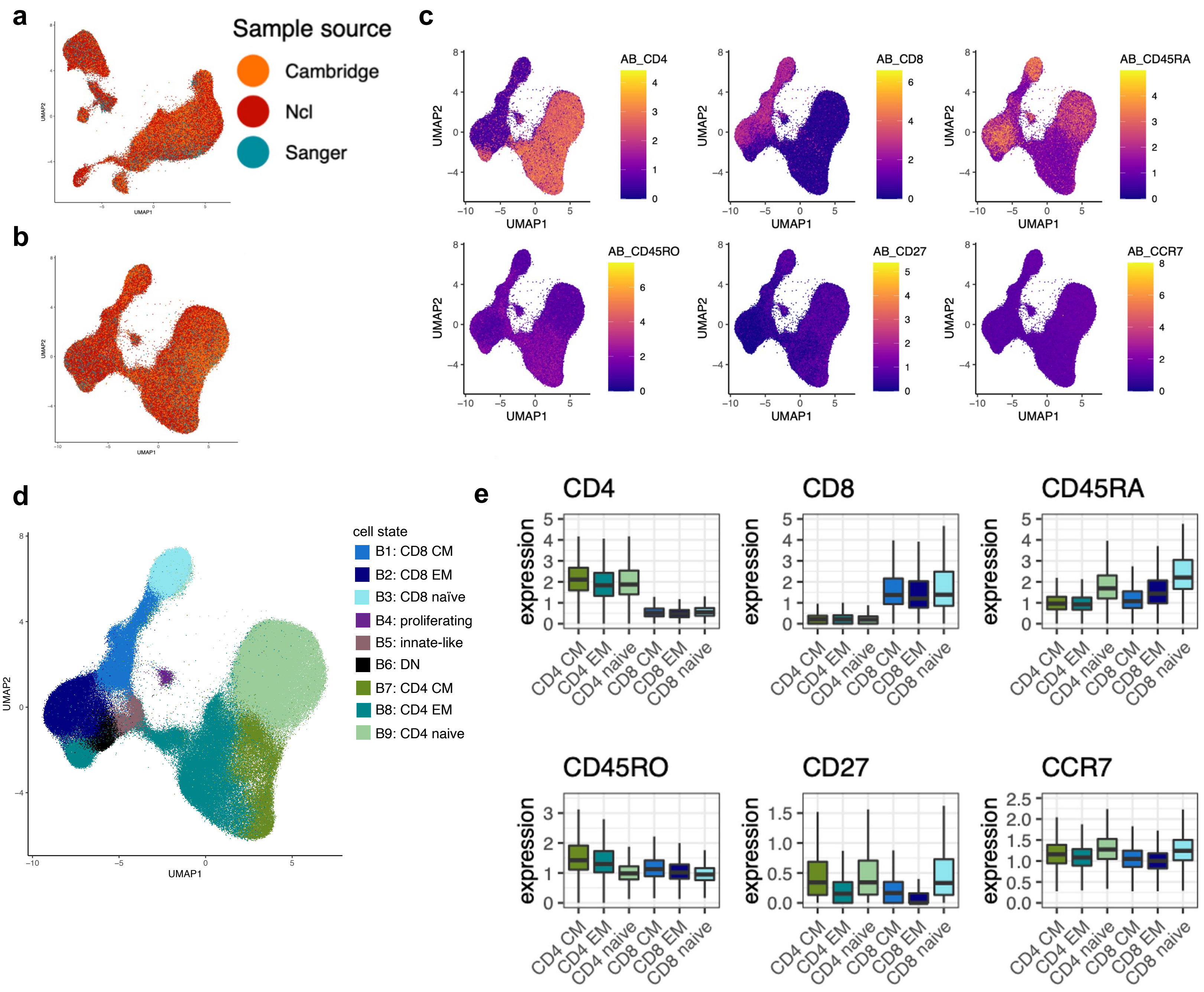


**Supplementary Figure 16. (a)** T cells from Dataset 3, projected into reference A (Supplementary Figure 1) by Symphony. **(b)** T cells from Dataset 3, projected into reference B (Supplementary Figure 2) by Symphony. For both projections, the only single-cell modality used from Dataset 3 is mRNA. **(c)** Dataset 3 T cells projected into reference B, colored by CLR-normalized surface protein expression values, which are masked from the projection process. **(d)** Dataset 3 T cells colored by reference B annotations assigned via Symphony and k-nearest-neighbors (k=5). **(e)** CLR-normalized protein expression (y-axis) measured in Dataset 3 is consistent with our cell state annotations, even though our annotation pipeline does not use protein information from Dataset 3. After learning mRNA-protein covariation from Datasets 1 and 2, our annotation pipeline can impute protein-defined cell states based on mRNA alone in new datasets.

#### Supplementary Figure 17

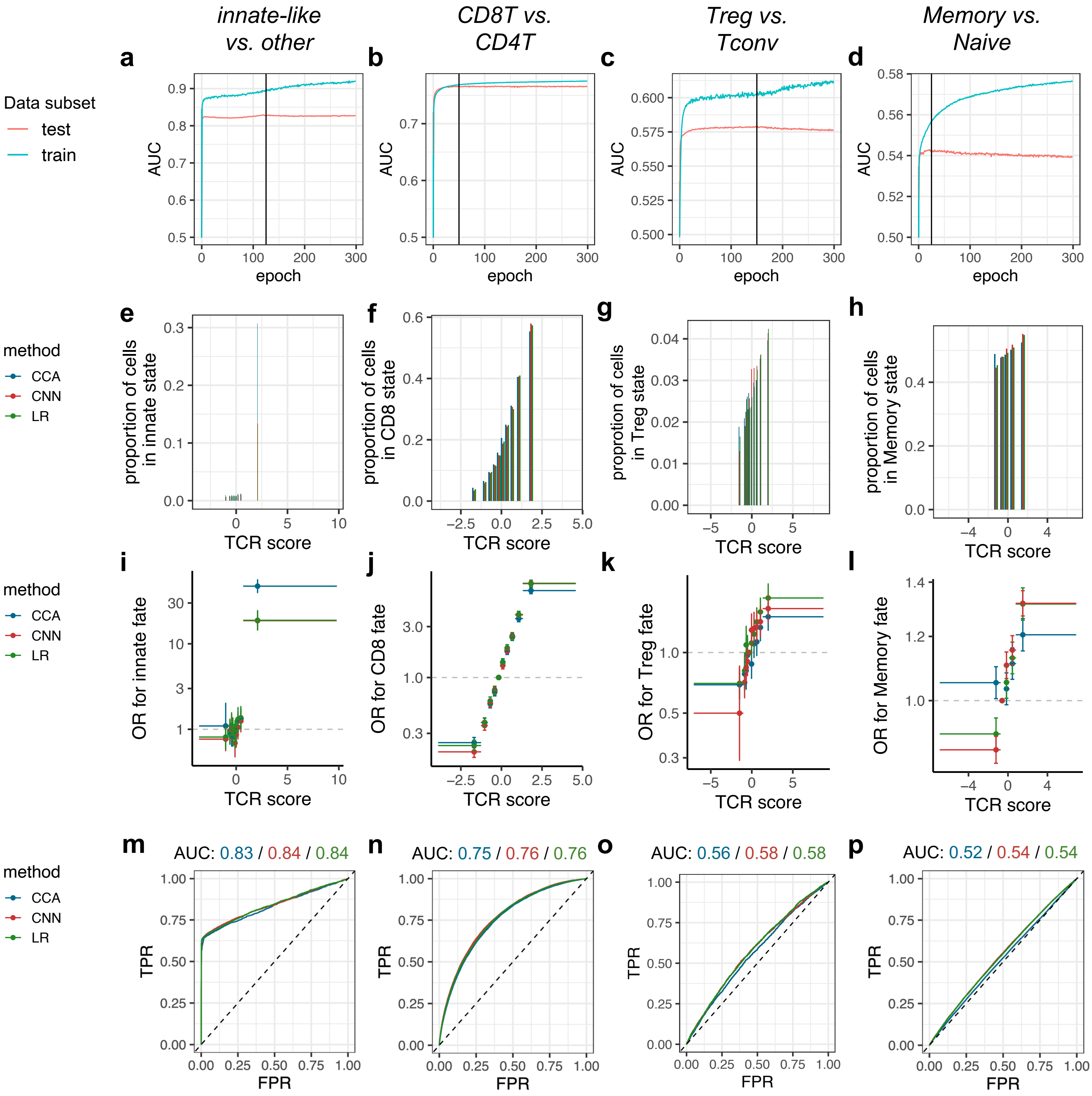


**Supplementary Figure 17.** With Dataset 3 as an external validation cohort, we compared TCR scoring functions from rCCA (blue), ridge-regularized logistic regression (green), and Convolutional Neural Networks (CNN, red). All three scoring functions were trained on the same observations (70% of clones from Dataset 1 and Dataset 2). For both training and testing data, expanded T cell clones were de-duplicated by picking one representative cell at random. **(e)** Each bar represents a decile of TCR score computed by one of the methods. Y-axis denotes the proportion of T cell clones from Dataset 3 observed in the innate-like (*PLZF*$^{high}$, cluster A9) T cell state. **(f-h)** Analogous to (e), for CD8 T cell fate, Treg fate, and memory fate, respectively. **(i)** Each point represents a decile of TCR score computed by one of the methods, with a horizontal bar spanning from its minimum to maximum value. We compute the odds ratio (OR, y-axis) for the innate-like (*PLZF*$^{high}$, cluster A9) T cell state for T cells in each decile compared to T cells in the fifth decile. 95% CIs (error bars) and *P* values computed via mixed-effects logistic regression. **(j-l)** Analogous to (e), for CD8 T cell fate, Treg fate, and memory fate, respectively. **(m)** Receiver operating characteristic (ROC) curve for TCR-based classifiers of innate-like T cell fate. **(n-p)** Analogous to (m), for CD8 T cell fate, Treg fate, and memory fate, respectively.

### Supplementary Figure 18

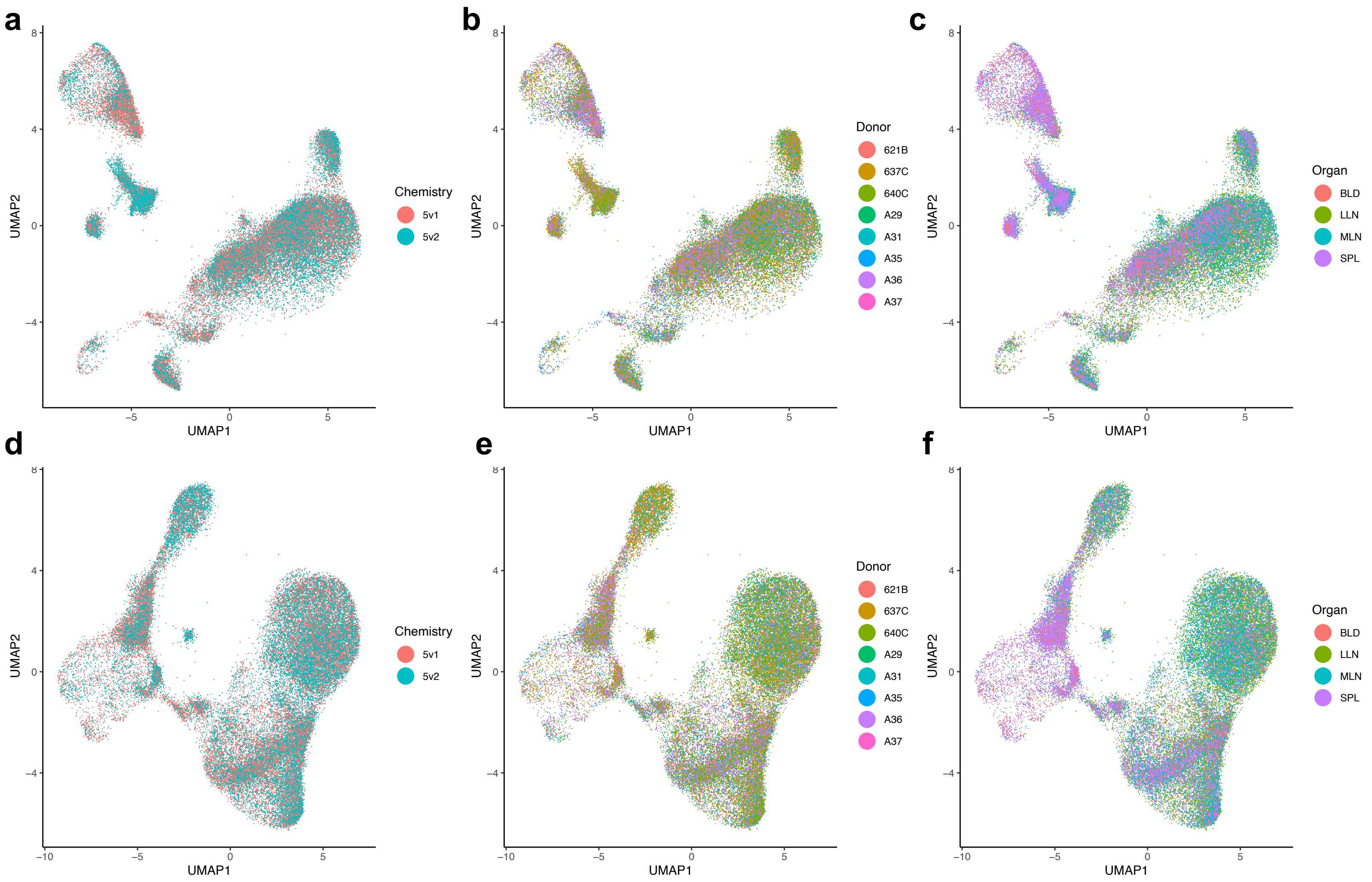


**Supplementary Figure 18. (a-c)** Dataset 4 T cells projected into reference A (Supplementary Figure 1) by Symphony (Methods). **(d-f)** Dataset 4 T cells projected into reference B (Supplementary Figure 2) by Symphony (Methods).

### Supplementary Figure 19

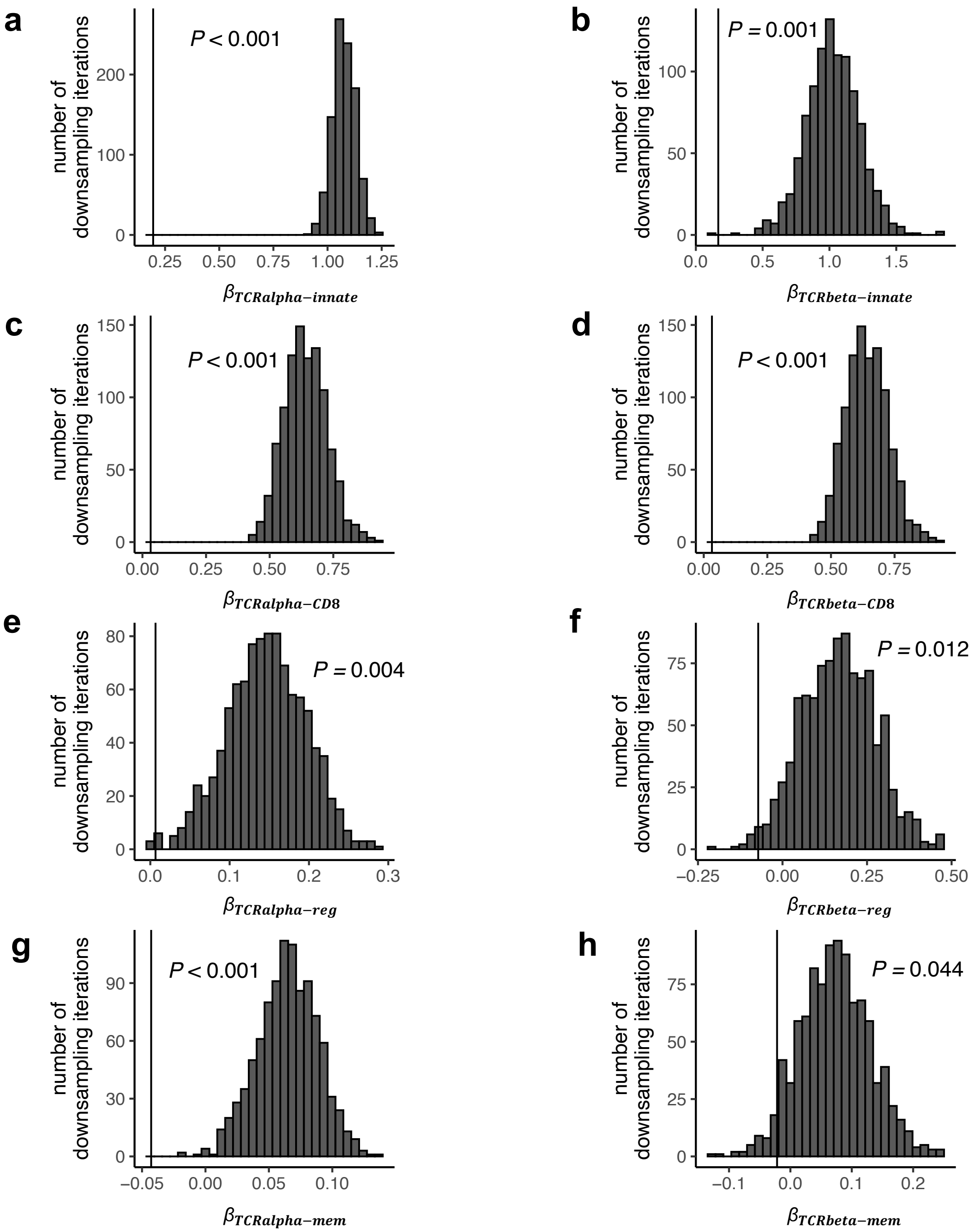


**Supplementary Figure 19. (a)** Histogram of $\beta_{TCRalpha-innate}$ estimates through 1000 iterations of down-sampling productive TCRs in Dataset 4 to match the number of nonproductive TCRs in Dataset 4. We estimate $\beta_{TCRalpha-innate}$ by mixed effects logistic regression, predicting innate-like T cell state based on the TCR scoring function TCRalpha-innate. Black vertical line marks estimate for $\beta_{TCRalpha-innate}$ when applied to nonproductive TCRs. Empirical *P* value tests the hypothesis that the association between TCRalpha-innate and innate-like T cell state is the same for nonproductive and productive TCRs. **(b)** Analogous to (a), for $\beta_{TCRbeta-innate}$ and innate-like T cell state, **(c)** Analogous to (a), for $\beta_{TCRalpha-CD8}$ and CD8T cell state, **(d)** Analogous to (a), for $\beta_{TCRbeta-CD8}$ and CD8T cell state, **(e)** Analogous to (a), for $\beta_{TCRalpha-reg}$ and Treg cell state, **(f)** Analogous to (a), for $\beta_{TCRbeta-reg}$ and Treg cell state, **(g)** Analogous to (a), for $\beta_{TCRalpha-mem}$ and memory T cell state, **(h)** Analogous to (a), for $\beta_{TCRbeta-mem}$ and memory T cell state.

#### Supplementary Figure 20

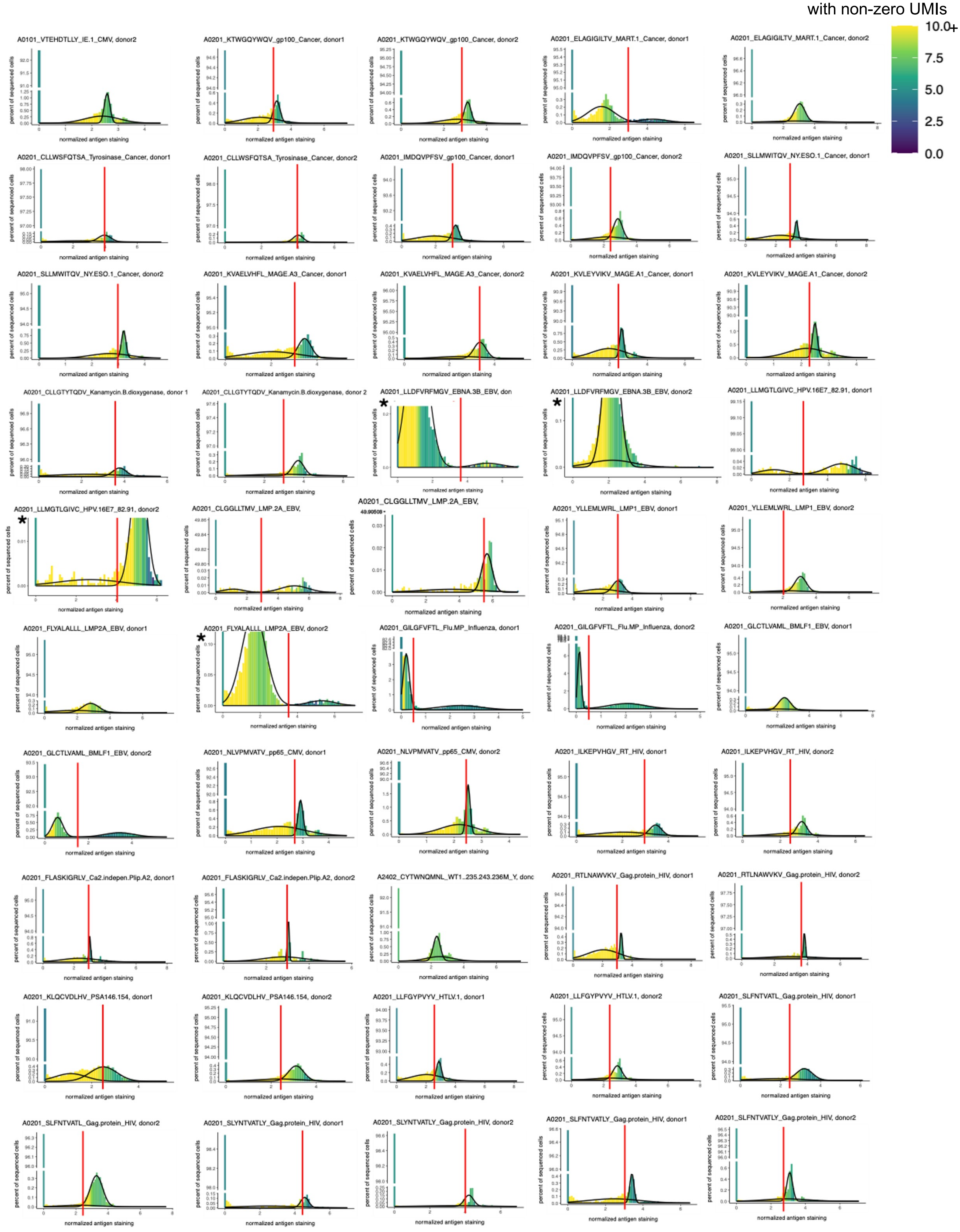

### Supplementary Figure 20

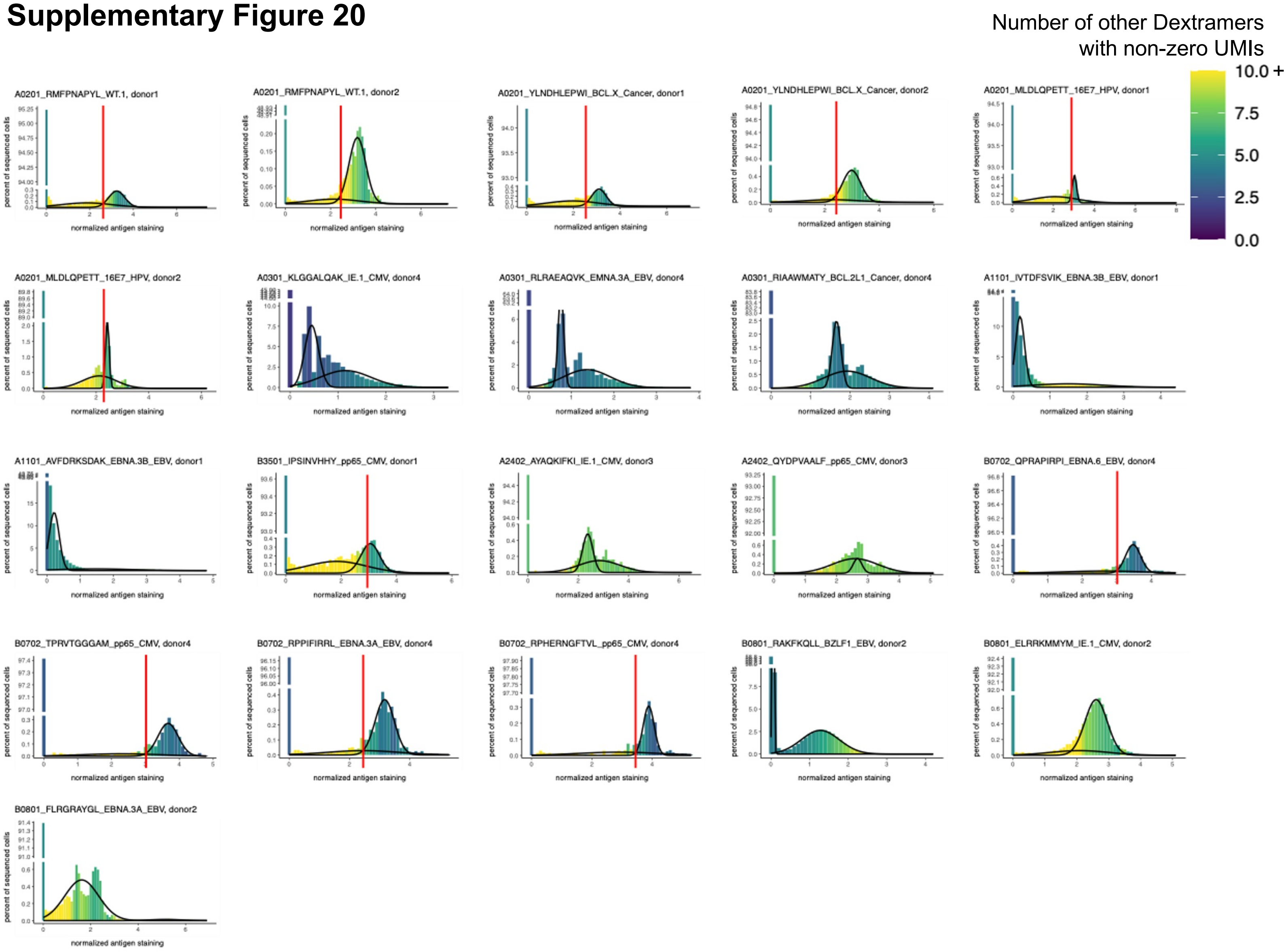


**Supplementary Figure 20.** Each plot represents one HLA-matched Dextramer-donor pair from Dataset 5. Histogram displays the distribution of UMI counts for the given Dextramer, normalized by a custom procedure based on negative binomial regression residuals (see Methods). Black outlines denote the distributions inferred by a Gaussian mixture model (two components, R package “mclust” v5.4.8). Red vertical lines mark the manual gates we set to delineate antigen-binding T cells after careful visual inspection. For plots lacking a red vertical line, no antigen-binding population was discernible.

### Supplementary Figure 21

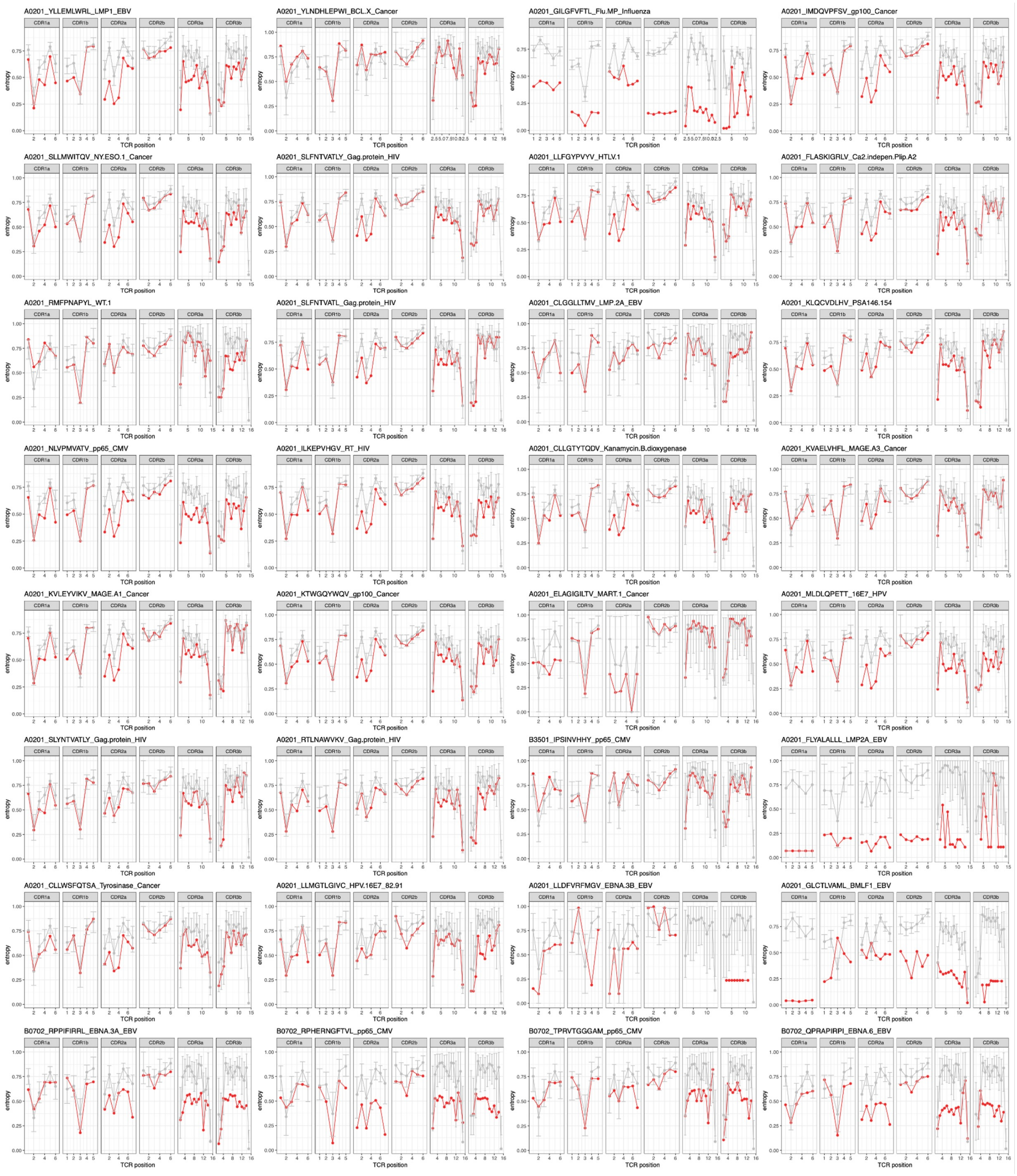


**Supplementary Figure 21.** Each plot (of six panels) represents one antigen-specific T cell population from Dataset 5. In red, we plot amino acid entropy at each TCR position (log2-normalized Shannon entropy) for the group of T cells inferred to recognize the given antigen. In gray, we show the same entropy calculations for 1000 random samples of a matched number of cells from Dataset 5. The point denotes the mean; the error bar denotes the minimum and maximum entropies observed in the 1000 random samples.

#### Supplementary Figure 22

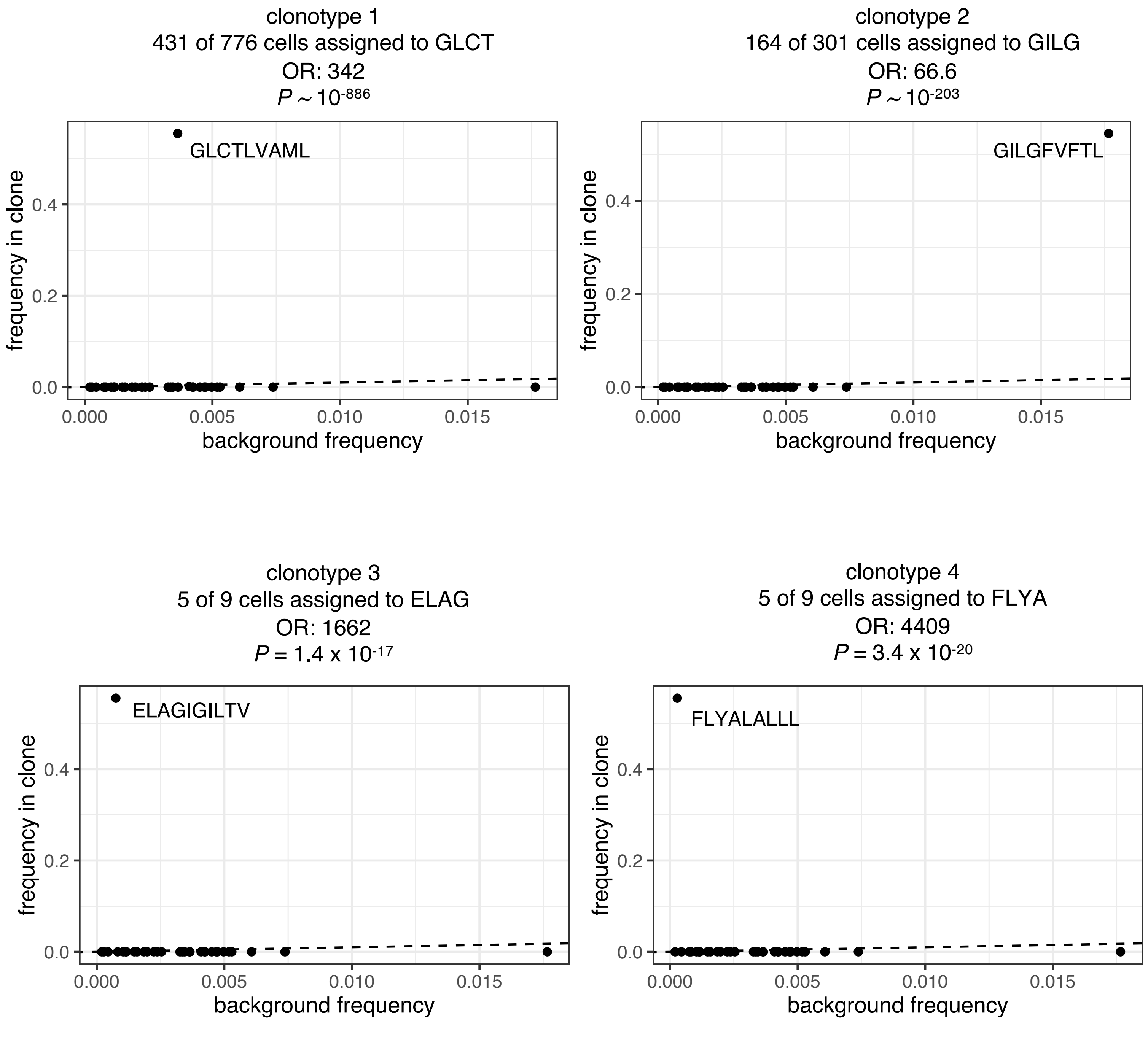


**Supplementary Figure 22**. Each plot represents an expanded T cell clone from Dataset 5. Each point represents a pMHC Dextramer assignment. We compare the frequency of each Dextramer assignment in the the expanded clone (y-axis) to the frequency of its assignment in the entire dataset (x-axis). We report the odds ratio (OR) for the Dextramer with the greatest frequency in the expanded clone [odds(Dextramer j) for T cells in clone i]/[odds(Dextramer j) for T cells not in clone i]). *P* values are computed by hypergeometric test.

#### Supplementary Figure 23

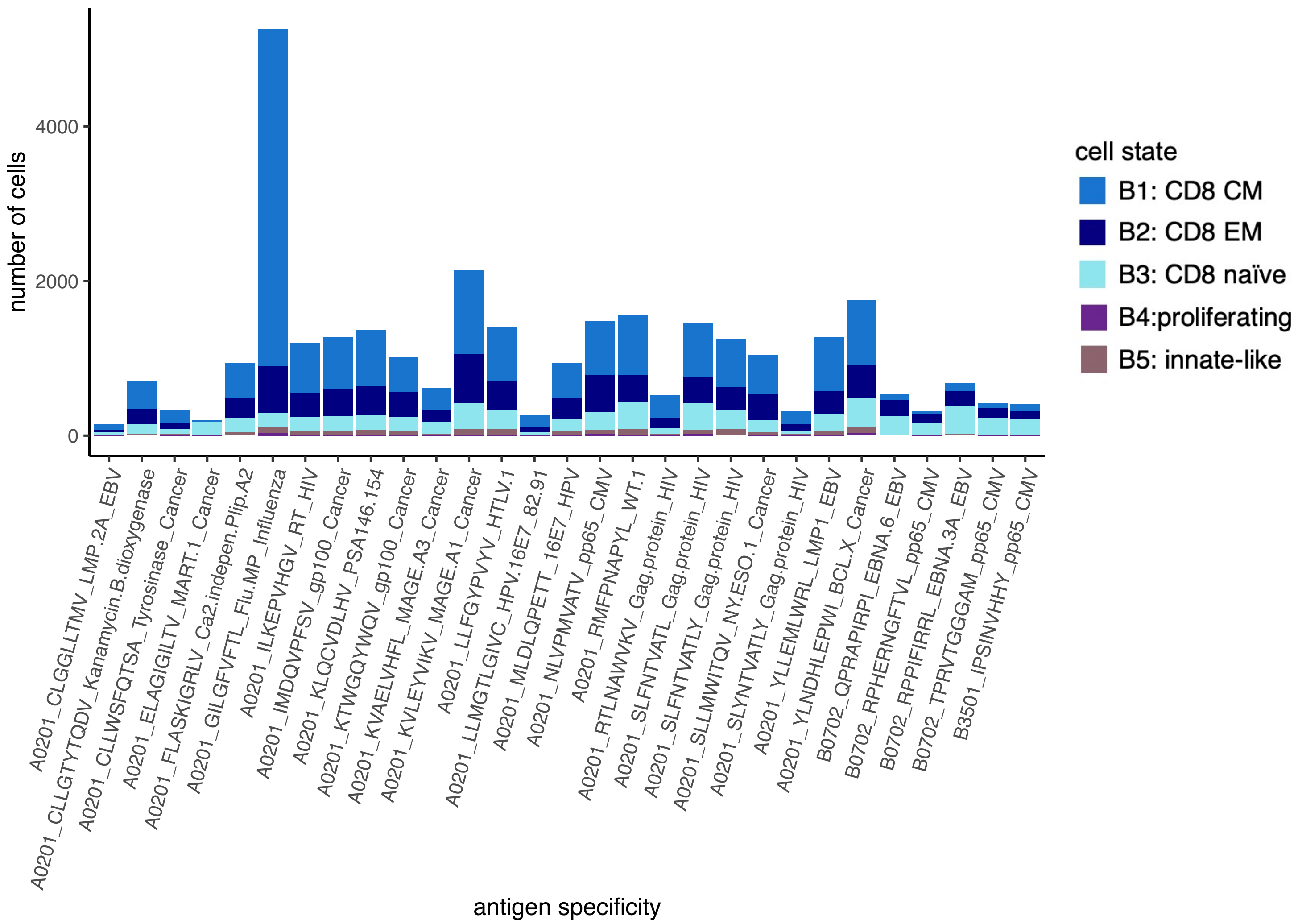


**Supplementary Figure 23.** Cell counts among each of the antigen-specific populations in Dataset 5.

#### Supplementary Figure 24

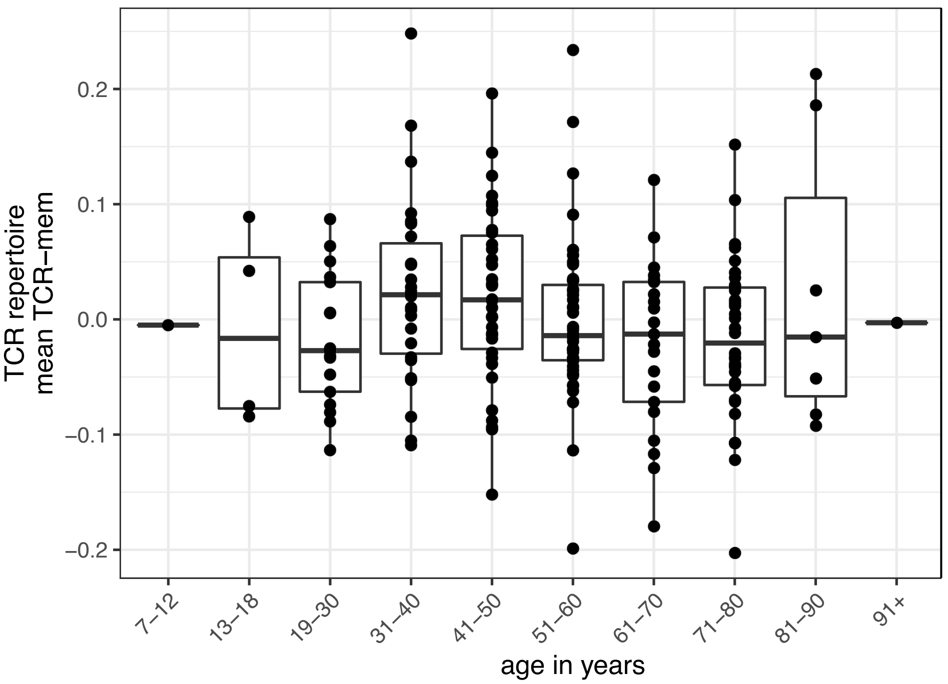


**Supplementary Figure 24**. Mean TCR-mem score for each individual's TCR repertoire sample (y-axis), plotted against the individual's age bracket (x-axis). Data consists of training observations from Dataset 1 and Dataset 2.

#### Supplementary Figure 25

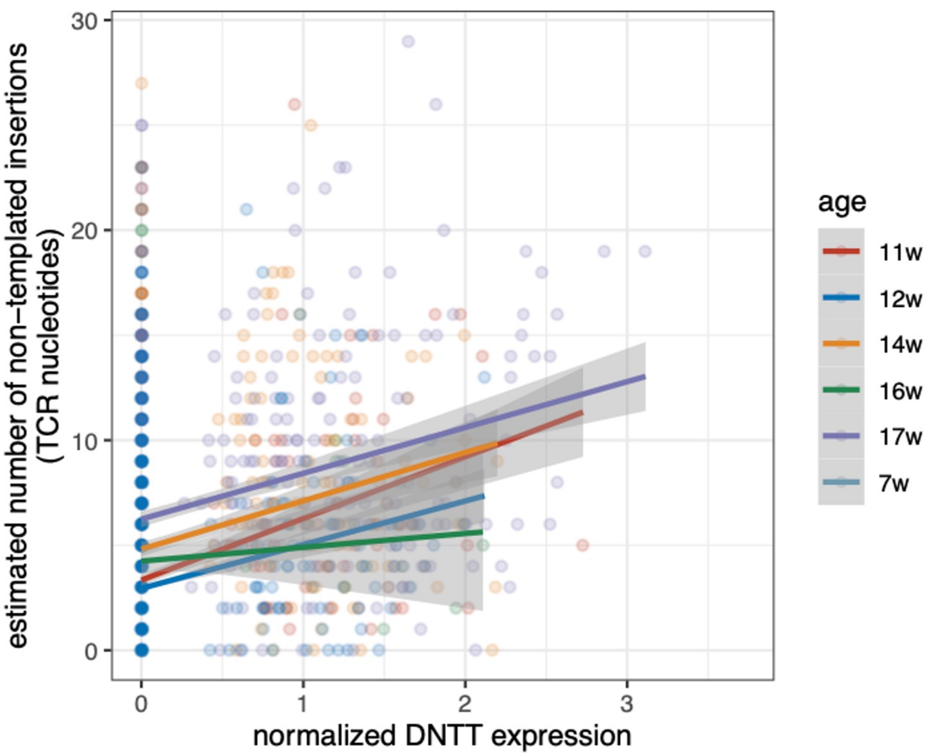


**Supplementary Figure 25.** Number of TCR nucleotide insertions estimated by IGoR (summed across the alpha and beta chains) for each prenatal thymic T cell (y-axis) in Dataset 6, plotted against *DNTT* expression (log(CP10K+1)). *DNTT* encodes Terminal Deoxynucleotidyl Transferase (TdT), the enzyme responsible for non-templated TCR nucleotide insertions. Line-of-best-fit computed for each gestational age category ("w" = post-conception weeks) by a linear model (geom_smooth(method="lm") from R package "ggplot2", v3.3.6).

#### Supplementary Figure 26

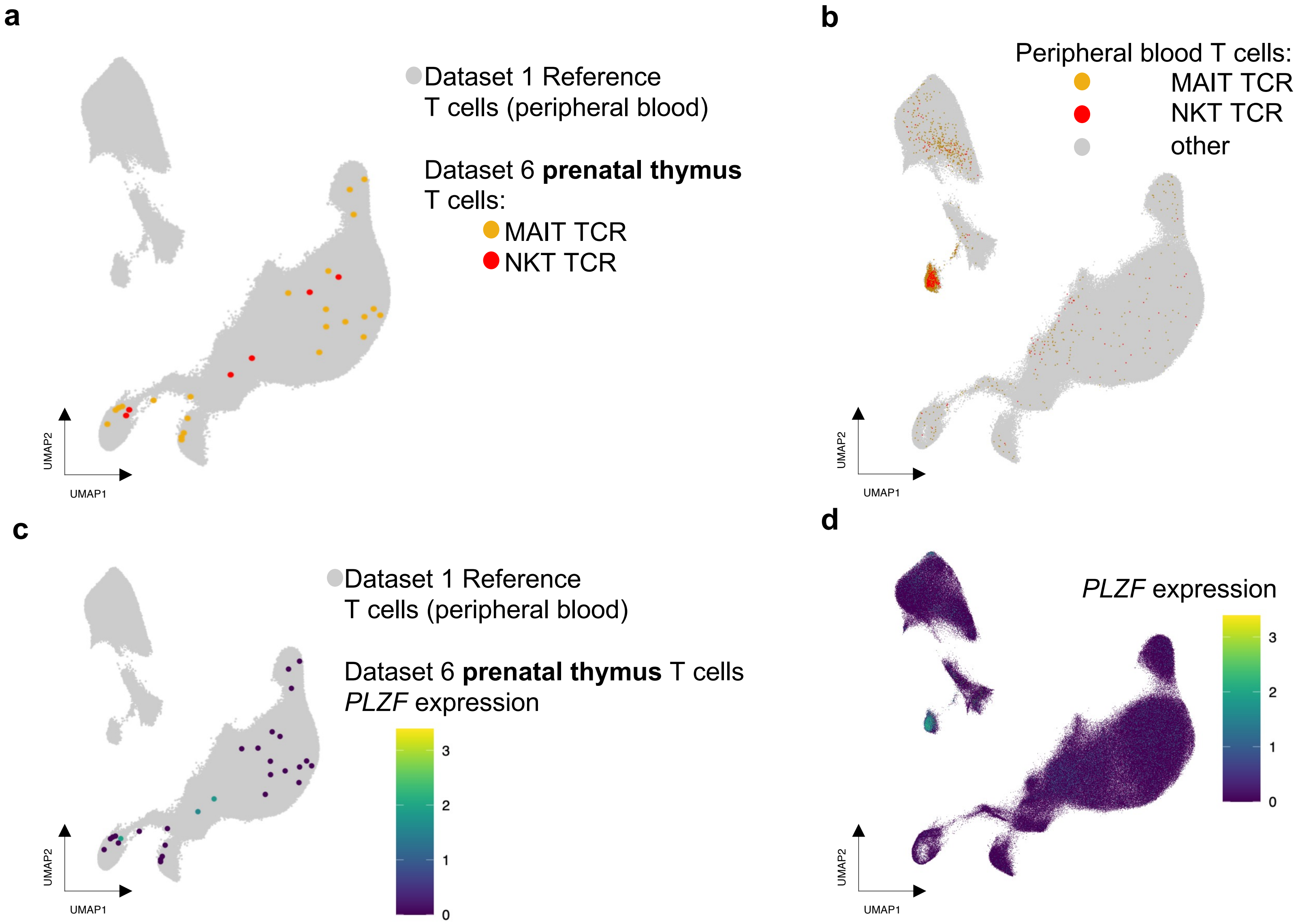


**Supplementary Figure 26. (a)** Dataset 6 (prenatal thymic) T cells with canonical MAIT TCRs (colored yellow, *TRAV1-2* and select *TRAJ* and *TRBV* genes, see Methods) or NKT TCRs (colored red, *TRAV10-TRAJ18-TRBV25*) projected into our T cell state reference UMAP constructed from Dataset 1 (colored gray). **(b)** UMAP of Dataset 1 T cells, colored yellow if their paired TCR uses canonical MAIT TCR genes, red if their paired TCR uses canonical NKT TCR genes, and gray otherwise. **(c)** Dataset 6 (prenatal thymic) T cells colored by *PLZF* expression (log(CP10K+1)) and projected into our T cell state reference UMAP. **(d)** UMAP of Dataset 1 T cells, colored by *PLZF* expression (log(CP10K+1)).
